# Transcriptomic Hallmarks of Mortality Reveal Universal and Specific Mechanisms of Aging, Chronic Disease, and Rejuvenation

**DOI:** 10.1101/2024.07.04.601982

**Authors:** Alexander Tyshkovskiy, Daria Kholdina, Kejun Ying, Maria Davitadze, Adrian Molière, Yoshiyasu Tongu, Tomoko Kasahara, Leonid M Kats, Anastasiya Vladimirova, Alibek Moldakozhayev, Hanna Liu, Bohan Zhang, Uma Khasanova, Mahdi Moqri, Jeremy M. Van Raamsdonk, David E. Harrison, Randy Strong, Takaaki Abe, Sergey E. Dmitriev, Vadim N. Gladyshev

## Abstract

Health is strongly affected by aging and lifespan-modulating interventions, but the molecular mechanisms of mortality regulation remain unclear. Here, we conducted an RNA-seq analysis of mice subjected to 20 compound treatments in the Interventions Testing Program (ITP). By integrating it with the data from over 4,000 rodent tissues representing aging and responses to genetic, pharmacological, and dietary interventions with established survival data, we developed robust multi-tissue transcriptomic biomarkers of mortality, capable of quantifying aging and change in lifespan in both short-lived and long-lived models. These tools were further extended to single-cell and human data, demonstrating common mechanisms of molecular aging across cell types and species. Via a network analysis, we identified and annotated 26 co-regulated modules of aging and longevity across tissues, and developed interpretable module-specific clocks that capture aging- and mortality-associated phenotypes of functional components, including, among others, inflammatory response, mitochondrial function, lipid metabolism, and extracellular matrix organization. These tools captured and characterized acceleration of biological age induced by progeria models and chronic diseases in rodents and humans. They also revealed rejuvenation induced by heterochronic parabiosis, early embryogenesis, and cellular reprogramming, highlighting universal signatures of mortality, shared across models of rejuvenation and age-related disease. They included *Cdkn1a* and *Lgals3*, whose human plasma levels further demonstrated a strong association with all-cause mortality, disease incidence and risk factors, such as obesity and hypertension. Overall, this study uncovers molecular hallmarks of mammalian mortality shared across organs, cell types, species and models of disease and rejuvenation, exposing fundamental mechanisms of aging and longevity.

## INTRODUCTION

Aging is associated with the systemic accumulation of damage, leading to a gradual loss of function, deterioration of organismal health, and an increased mortality rate^1^. However, these manifestations of aging can be influenced by numerous genetic, dietary, and pharmacological lifespan-shortening and longevity interventions, ranging from Hutchinson-Gilford progeria syndrome (HGPS)^2–5^ and the *Klotho* knockout (KO) model^6,7^ to caloric restriction (CR)^8^, rapamycin^9–11^, and dwarf models associated with growth hormone deficiency^12–14^. Since 2004, the Interventions Testing Program (ITP)^15^ has evaluated the effects of over 50 compounds and combinations on the survival of genetically heterogeneous UM-HET3 female and male mice. It established that more than 10 treatments can effectively extend lifespan in female and/or male mice, including rapamycin^9,10,16,17^, acarbose^18–20^, canagliflozin^21^, 17-α-estradiol^18,19,22^, protandim^18^, captopril^23^, glycine^24^, nordihydroguaiaretic acid (NDGA)^25,26^, astaxanthin^27^, and meclizine^27^, along with two combinations of compounds, rapamycin with metformin^18^ and rapamycin with acarbose^23^. Previous studies by our group and others focused on the identification of molecular signatures of aging^28–32^ or lifespan-regulating models^33–36^. However, a unified analysis of mortality-associated mechanisms driven by mammalian aging and by various lifespan-shortening and longevity interventions has been lacking.

Rapid accumulation of high-throughput data, including transcriptomics, proteomics, and DNA methylation (DNAm) allowed for the development of chronological epigenetic^37–41^, transcriptomic^42–45^, and proteomic^46^ clocks that can predict the age of humans or animals based on the molecular profiles of their tissues and individual cell types^47,48^. Recently, the first pan-mammalian chronological epigenetic clock was developed^49^, highlighting the existence of conserved molecular biomarkers of aging across organs and species. However, since age-related signatures reflect not only detrimental but also neutral and adaptive changes^41^, biomarkers of mortality and clinical measurements provide a more direct evaluation of an aggregated level of damage (i.e., biological age) and allow to predict health outcomes more effectively^50^. Previously, DNAm was used to develop blood-specific clocks that estimate human time to death and healthspan^50,51^. However, current mortality-associated clocks are limited to human blood cells, and their application is further restricted by the low amount of single-cell DNAm data and low interpretability of features (methylation of individual CpG sites) used to train these models.

In contrast, detailed characterization of mammalian genes makes transcriptomic models highly interpretable and widely applicable to various models of aging and longevity interventions. Implementation of transcriptomic tools is further facilitated by wide availability of single-cell RNA-seq (scRNA-seq) data, allowing to examine the level of molecular damage across cell types and characterize cells most vulnerable to a given treatment. Therefore, the development of mortality transcriptomic clocks based on gene expression profiles and their functional components across organs and mammalian species could reveal universal and specific molecular mechanisms of the established and novel models of healthspan regulation, rejuvenation and aging.

To fill this gap, we performed RNA sequencing (RNA-seq) of tissues of mice subjected to 20 interventions with known effects on lifespan tested by the ITP. By integrating these and publicly available transcriptomic data encompassing 4,539 rodents, we developed robust mammalian mortality multi-tissue clocks and deconstructed them into functional modules associated with aging and longevity. We demonstrated their utility in assessing longevity and detrimental mechanisms of progeria and aging-related diseases. By including human data, we constructed multi-species multi-tissue transcriptomic clocks of chronological age and mortality. These tools allowed us to gain numerous novel insights into longevity, rejuvenation, and chronic diseases, revealing shared and model-specific signatures of mortality. Using them, we also established universality of the aging-related damage accumulation paradigm across cell types and identified a compound with a substantial rejuvenation effect that resembles heterochronic parabiosis. Finally, we developed an interactive online platform TACO for preprocessing of RNA-seq data and implementation of the constructed transcriptomic clocks.

## RESULTS

### Gene expression regulation of mammalian aging and longevity

To identify gene expression hallmarks of maximum lifespan and mortality, we performed RNA-seq of liver samples from 170 22-month-old UM-HET3 female and male mice subjected to 20 pharmacological treatments tested by the ITP^16,17,21–23^ together with the corresponding age- and sex-matched controls (Supplementary Table 1A; Fig. 1A). These interventions included those with no statistically significant effect on mouse lifespan, such as minocycline^16^ and nicotinamide riboside (NR)^22^, as well as those leading to extension of lifespan in males and/or females, such as the antidiabetic drug canagliflozin^21^, antihypertensive angiotensin-converting enzyme (ACE) inhibitor captopril^23^, late-life treatment with rapamycin^16^, middle- and late-life treatment with a ‘non-feminizing’ derivative of estrogen 17-α-estradiol^22^, as well as a whole-life and middle-life treatment with a combination of rapamycin and acarbose^23^, one of the most effective pharmacological longevity interventions in mammals up to date. We further expanded our dataset with 12 liver samples from 4-6-month-old male and female mice. This allowed us to assess molecular mechanisms of lifespan regulation and aging and integrate these features into a single mortality model.

**Fig. 1.**
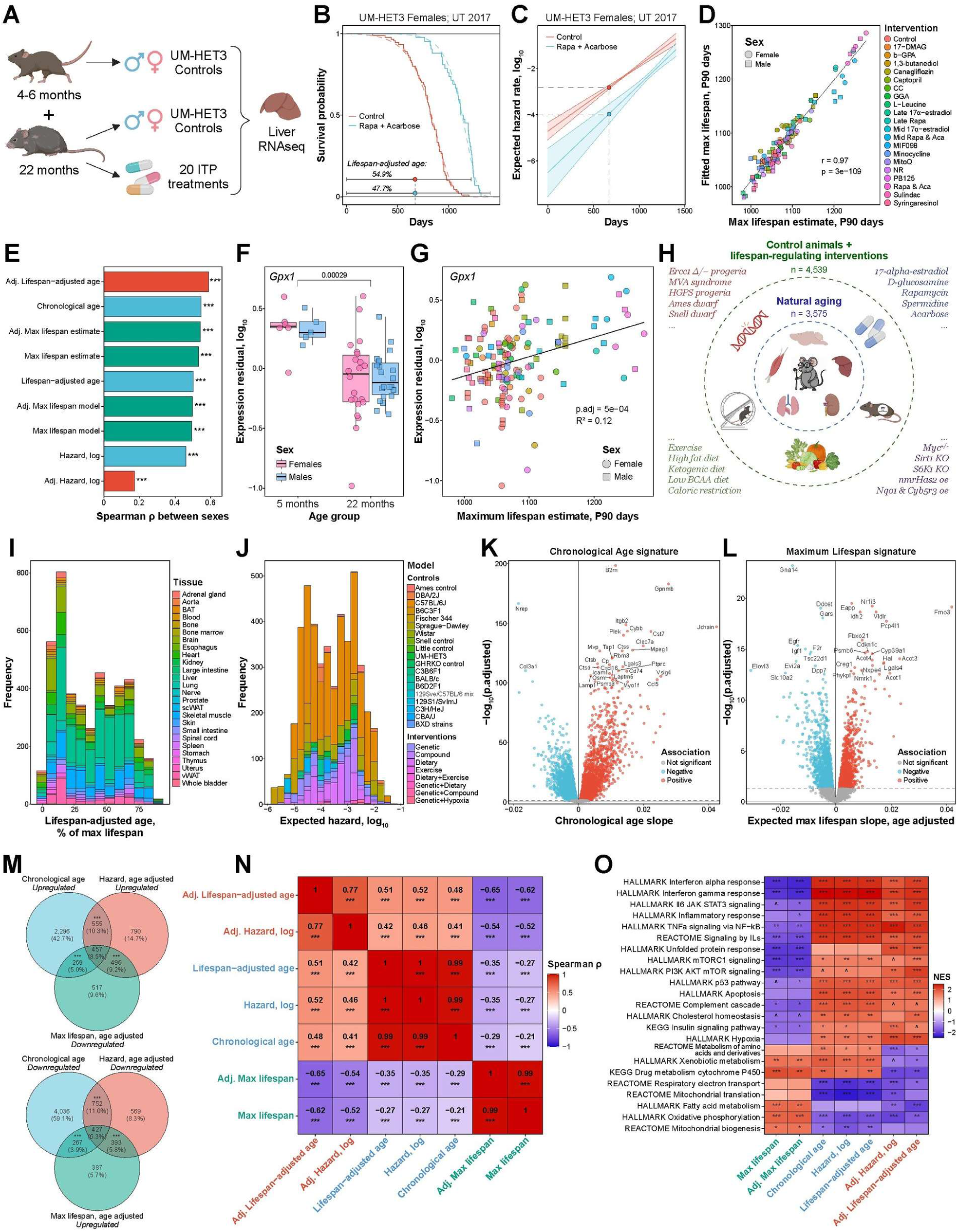
Gene expression signatures reflect molecular mechanisms of mammalian aging, maximum lifespan, and mortality. **A.** Overview of samples from the Interventions Testing Program (ITP) subjected to RNA-seq. **B.** Example of a fitted survival curve of ITP mice. UM-HET3 control females and females subjected to rapamycin and acarbose treatment are shown from the University of Texas 2017 ITP cohort. Estimates of lifespan-adjusted age for control and treated animals of the same chronological age are shown in the text. **C.** Expected all-cause mortality curves (in log scale) of animals from (B), fitted with a Gompertz model. 95% confidence intervals are shown with shaded areas. Dotted lines reflect estimates of expected hazard for age-matched control and treated organisms. **D.** Estimates of expected maximum lifespan (90^th^ percentile) for various ITP cohorts derived from survival data (x axis) and from fitted Gompertz models (y axis). Pearson correlation coefficient and p-value are shown in text. Rapa: Rapamycin; Aca: Acarbose; CC: Candesartan Cilexetil; GGA: Geranylgeranyl Acetone; NR: Nicotinamide Riboside; Late: Late-life; Mid: Mid-life. **E.** Correlation between male and female gene expression signatures of lifespan (green bars), chronological age, lifespan-adjusted age and mortality unadjusted (blue) and adjusted for chronological age (red) from ITP data. For each trait, union of top 1000 associated genes in females and males (with the lowest p-value) was used to estimate Spearman’s correlation coefficient. Asterisks reflect BH-adjusted p-values. **F.** Expression of *Gpx1* adjusted for sex in young (left) and old (right) control UM-HET3 mice. BH adjusted p-value corresponding to age difference in expression is shown in text. **G.** Association between expected maximum lifespan of ITP cohorts estimated from survival data (x axis) and expression of *Gpx1* adjusted for sex (y axis). BH adjusted p-value is shown in text. **H.** Overview of transcriptomic data from mice and rats aggregated in the study. Number of samples for control (blue) and all animals (green) corresponding to various strains, sexes, ages and lifespan-regulating interventions is provided. Icons reflect most represented organs and types of interventions. Examples of lifespan-shortening and -extending interventions included in the dataset at the level of gene expression and survival are shown. KO: Knockout; oe: overexpression. **I.** Distribution of lifespan-adjusted ages (x axis) and tissues (shown by color) for all samples covered in the aggregated meta-dataset (n=4,539). BAT: Brown Adipose Tissue; scWAT: Subcutaneous White Adipose Tissue; vWAT: Visceral White Adipose Tissue. **J.** Distribution of expected all-cause mortality (in log scale, x axis), available strains and types of interventions (shown by color) for all samples in the meta-dataset (n=4,539). **K.** Gene expression signatures of chronological age in control animals identified from the whole meta-dataset (n=3,575). Slope of association and BH-adjusted p-value (in log scale) are shown on x and y axis, respectively. Top genes associated with age are shown in the text. **L.** Gene expression signatures of expected maximum lifespan adjusted for chronological age identified from the whole meta-dataset (n=4,539). Maximum lifespans were estimated with Gompertz model based on survival data for the corresponding sex, strain and intervention. **M.** Venn diagrams of genes significantly associated (BH-adjusted p-value < 0.05) with chronological age (blue), expected mortality (red) and maximum lifespan (green) adjusted for age. Statistical significance of pairwise overlaps was assessed with Fisher’s exact test and is shown with asterisks. **N.** Correlation matrix of normalized enrichment scores (NES) of pathways associated with signatures of maximum lifespan (green), chronological age, lifespan-adjusted age and mortality unadjusted (blue) and adjusted for chronological age (red), identified from meta-dataset. NES were estimated with GSEA. Correlation coefficient and statistical significance are indicated with text and asterisks, respectively. **O.** Functional enrichment (GSEA) of transcriptomic signatures of maximum lifespan (green), chronological age, lifespan-adjusted age and mortality unadjusted (blue) and adjusted for chronological age (red), identified from meta-dataset. Statistical significance is shown with asterisks. Only functions significantly enriched by at least one signature are shown (BH-adjusted p-value < 0.05). The whole list of enriched functions is in Supplementary Table 2B. ^ p.adj < 0.1; * p.adj < 0.05; ** p.adj < 0.01; *** p.adj < 0.001.

To calculate expected all-cause mortality rate for each animal presented in our data, we fitted survival curves obtained from original ITP studies with Gompertz model (Fig. 1B) separately for each cohort, sex, site, and experimental group. Since every ITP compound was tested across 3 independent sites and survival of control animals was tested across multiple cohorts, we aggregated parameters of individual Gompertz functions for each sex and experimental group using a mixed-effect model to increase the reliability of mortality estimates (Extended Data Fig. 1A). As expected, aggregated Gompertz models produced intermediate estimates of hazard rate and survival across studies (Extended Data Fig. 1A-B). We then calculated the logarithm of hazard rate for each sample in our dataset by providing its chronological age to the corresponding aggregated and cohort-specific mortality models and taking the average of the resulting two estimates (see Methods). This algorithm allowed us to estimate expected mortality rate for every mouse based on its age, sex, cohort, and treatment data (Fig. 1C). As a complementary metric of biological age, we also calculated lifespan-adjusted age, defined as a chronological age divided by the maximum lifespan (99.9^th^ percentile) of the respective mouse model estimated with the Gompertz function (Fig. 1B). This metric, therefore, can be considered as a fraction of maximum lifespan that has passed for the provided animal. 90^th^ percentile lifespan estimates predicted by Gompertz models for each experimental group, sex, and cohort were well correlated with the corresponding lifespan estimates derived from survival data (Pearson’s r = 0.97) (Fig. 1D), suggesting that this mortality model captures differences in survival dynamics across treatments and sexes, and can be used to estimate expected hazard rate and maximum lifespan.

Principal component analysis (PCA) performed on the filtered and normalized dataset revealed separation of samples by sex according to the first principal component (Extended Data Fig. 1C). To identify transcriptomic biomarkers of maximum lifespan, expected mortality and lifespan-adjusted age, we utilized a regression model and tested if gene log-expression is associated with these features after adjustment for sex and site. The same approach was employed to detect signatures of chronological age between young (4-6 months) and old (22 months) control samples. Using a False Discovery Rate (FDR) threshold of 0.05, we revealed 1272, 1457, 868 and 700-795 genes significantly associated with chronological age, lifespan-adjusted age, expected hazard rate (in log scale) and maximum lifespan, respectively (Extended Data Fig. 1D-F). Age-related gene expression changes demonstrated statistically significant weak negative correlation with biomarkers of lifespan (Spearman rho between −0.05 and −0.2), both before and after adjustment for chronological age (Extended Data Fig. 1G), in agreement with our previous findings^28^. At the same time, signatures of lifespan-adjusted age and mortality rate demonstrated a stronger negative association with lifespan (Spearman rho between −0.13 and −0.37) and positive association with age (Spearman rho > 0.93), confirming that these metrics better reflect detrimental changes associated with aging and shorter lifespan. Regression analysis performed separately for males and females revealed a significant positive correlation between sexes for all estimated signatures (Fig. 1E). This suggests that molecular mechanisms of aging and longevity are generally shared across sexes, despite the presence of sex-specific molecular and lifespan-extending effects induced by many interventions, such as 17-α-estradiol and canagliflozin^21,22,33^.

Remarkably, one of the top genes positively associated with maximum lifespan (p.adjusted = 5.10^−4^) and negatively associated with chronological age and expected mortality (p.adjusted = 2.9.10^−4^) was *Gpx1*, encoding the selenoprotein glutathione peroxidase 1 (Fig. 1F-G). Accordingly, overexpression of this gene was shown to ameliorate age-related kidney pathologies in old mice, such as glomerulosclerosis and interstitial fibrosis^52^. In addition, we observed a similar association with age, mortality, and lifespan for several other genes involved in antioxidative response, including multiple cytochrome P450 genes (e.g., *Cyp2f2*, Extended Data Fig. 2A-B), and genes involved in fatty acid metabolism (e.g., *Acsm1*, Extended Data Fig. 2C-D). To characterize pathways, enriched for the identified gene expression signatures, we performed gene set enrichment analysis (GSEA) (Supplementary Table 2A). As at the level of genes, the functional signature of chronological age was positively correlated with mortality and lifespan-adjusted age (Spearman rho > 0.94) and negatively correlated with lifespan (Spearman rho < −0.31), both before and after adjustment for age (Extended Data Fig. 2E). Interestingly, correlations at the level of pathways were generally stronger than at the level of genes, suggesting that longevity interventions may affect the expression of different individual genes but ultimately lead to similar functional outcomes. Pathways positively associated with lifespan and negatively with mortality, both before and after adjustment for age, included oxidative phosphorylation, mitochondrial translation, xenobiotic metabolism, and fatty acid metabolism, while interferon signaling and hemostasis demonstrated the opposite associations (Extended Data Fig. 2F).

To characterize universal molecular mechanisms of rodent aging, mortality, and lifespan regulation that are reproduced across tissues, species, genetic backgrounds and interventions, we expanded our dataset with publicly available gene expression data from tissues of control healthy mice and rats with known chronological age (Fig. 1H). Thus, we integrated 3,575 samples from 26 tissues, covering a substantial range of rodent ages, from 4 to 943 days (Extended Data Fig. 3A). We also introduced 964 transcriptomic samples from mice subjected to various pharmacological, genetic, dietary, and environmental interventions with extending, shortening, or neutral effects on lifespan that have the associated published survival data, including HGPS progeria model^2^, caloric restriction^8^, spermidine^53^, acarbose^18,19,33^, pregnancy-associated plasma protein-A (PAPP-A) KO^54^, Ames and Snell dwarf models^12,55^, S6K1 deletion^56^, hypoxia^57^, and others (Fig. 1H). Overall, our meta-dataset included 4,539 mouse and rat gene expression profiles representing 26 tissues, 79 unique interventions, 96 independent studies, and various platforms (Supplementary Table 1B). Utilizing published survival data, we fitted cohort-specific and aggregated Gompertz mortality models for every presented strain, sex, and intervention using the algorithm explained above. We then used them to calculate expected hazard rate and lifespan-adjusted age for every sample in our gene expression dataset (Fig. 1I-J). Similar to the ITP data, the Gompertz function was able to capture differences in mortality dynamics across various rodent models, confirmed by a high correlation between the estimate of 90^th^ percentile lifespan predicted by the model and the lifespan estimate derived from the corresponding survival data (r = 0.995) (Extended Data Fig. 3B). As expected, hazard rate, as a composite metric, was positively correlated with chronological age (Extended Data Fig. 3D) but was also affected by different genetic backgrounds, progeria models, high-fat diets and lifespan-extending interventions.

Using aggregated data, we searched for robust multi-tissue transcriptomic biomarkers of aging, mortality, and maximum lifespan. To reduce variation in gene expression caused by batch effect and differences in baseline expression across organs and sexes, for every subset of samples corresponding to a particular dataset, tissue, and sex we centered their gene expression profiles around the median expression profile of a randomly chosen control group (see Methods). We then applied linear mixed-effect model, including data source and tissue as random terms and sex as a fixed term, to identify gene expression changes associated with difference in chronological age, lifespan-adjusted age, maximum lifespan (99.9^th^ percentile), and expected hazard rate (in log scale). Using FDR threshold of 0.05, we detected 3098 statistically significant signatures of maximum lifespan and 9059 to 9167 signatures of chronological age, lifespan-adjusted age and mortality at the level of gene expression (Fig. 1K). Interestingly, after adjustment for chronological age, 3213, 4099, and 4439 genes demonstrated statistically significant association with maximum lifespan, lifespan-adjusted age, and expected hazard rate, respectively (Fig. 1L; Extended Data Fig. 3C), pointing to systemic regulation of mortality and longevity at the molecular level.

Among top genes, whose expression is negatively associated with maximum lifespan of rodents and positively correlated with their mortality after adjustment for age, we found *Igf1* (p.adjusted < 3.6.10^−13^ for age-adjusted lifespan and mortality signatures), encoding insulin-like growth factor-1, a well-known regulator of lifespan in multiple species (Fig. 1L)^58^. Previously, we have shown that this gene is consistently downregulated in tissues of long-lived mammalian species^28^, supporting its crucial role in the longevity control both across and within species. Another gene negatively correlated with lifespan and positively associated with both mortality and chronological age was *Ddost* (p.adjusted < 3.7.10^−8^ for signatures of age-adjusted mortality and lifespan signatures; p.adjusted = 3.5.10^−4^ for aging signature), involved in processing of advanced glycation end products (AGEs)^59^. Its expression is elevated in people with Alzheimer’s disease (AD)^60^ and can serve as a prognostic biomarker of poor survival for patients with hepatocellular carcinoma (HCC)^61^ and gliomas^62^.

Among genes positively associated with lifespan and negatively correlated with mortality, we identified flavin-containing monooxygenase 3 gene *Fmo3* (p.adjusted < 3.9.10^−8^ for age-adjusted lifespan and mortality signatures), an established inhibitor of mTOR and activator of autophagy^63^. Its overexpression in mice was shown to ameliorate several aging-related hallmarks, reducing levels of pro-inflammatory cytokine interleukin-6, total cholesterol, triglyceride, and markers of senescence. Interestingly, *Fmo3* expression is also slightly upregulated with age (p.adjusted = 0.011; Extended Data Fig. 3E), while *Igf1* is downregulated in tissues of aged organisms (p.adjusted = 8.3.10^−7^), further supporting the idea that age-related biomarkers include both detrimental and compensatory molecular changes^41^. We also detected strong positive association with rodent lifespan and negative correlation with hazard rate for *Nmrk1* (p.adjusted < 1.3.10^−13^ for age-adjusted lifespan and mortality signatures) (Fig. 1L; Extended Data Fig. 3C), encoding a nicotinamide riboside kinase 1, an enzyme involved in biosynthesis of NAD^+^ precursor NMN^64^, whose association with aging and longevity is well established^28,65^.

Consistent with the ITP data, genes associated with chronological age had statistically significant co-directional overlap with biomarkers of age-adjusted mortality and counter-directional overlap with lifespan signatures (Extended Data Fig. 3F). This was evident both at the level of up- and downregulated genes (Fig. 1M), supporting the hypothesis that age-associated transcriptomic changes are generally prognostic of poor health outcomes even after adjustment for the organism’s age. We also detected significant correlations between these signatures, both at the level of individual genes (Extended Data Fig. 3G) and enriched functions (Fig. 1N). As expected, signatures of lifespan-adjusted age and expected mortality exhibited a stronger association with maximum lifespan, at the same time maintaining high correlation with chronological age.

GSEA revealed that genes negatively associated with maximum lifespan and positively associated with aging and mortality rate, both before and after adjustment for chronological age, were involved in interferon and interleukin signaling, p53 pathway, complement cascade, and mTOR and insulin signaling, while genes involved in oxidative phosphorylation and mitochondrial biogenesis showed the opposite associations (Fig. 1O; Supplementary Table 2B). We also detected significant positive association with lifespan and negative association with age-adjusted mortality for genes related to fatty acid metabolism and xenobiotic metabolism, supporting their role in regulation of longevity^33,35,66,67^ and clearance of accumulated molecular damage^1^.

### Rodent multi-tissue transcriptomic clocks of aging and mortality

To develop quantitative multi-tissue biomarkers of mammalian aging and mortality at the level of gene expression, we utilized the aggregated dataset to train machine learning models that predict chronological age, lifespan-adjusted age, and expected hazard rate of tissue samples. For the chronological clock, we trained an Elastic Net (EN) linear model with K-Fold cross-validation on the complete gene expression profiles of 3,575 samples from 26 tissues of healthy mice and rats not subjected to any interventions. Most organs in the dataset were well distributed across age groups, with the top represented tissues being liver (750 samples), skeletal muscle (521 samples), brain (499 samples) and kidney (275 samples) (Extended Data Fig. 3A). As a response variable, we used chronological age divided by the species maximum recorded lifespan, according to AnAge^68,69^, in a linear way or with log-log transformation^49^. We also tested 2 normalization methods of gene expression: scaling and YuGene normalization^70^, shown to be effective for standardization of transcriptomic data from different sources^71^ (see Methods). The trained model was able to predict absolute chronological age of animals on the test set with R^2^ = 0.88, Pearson’s r = 0.94 and mean absolute error (MAE) = 2.2 months (Fig. 2A). To get robust estimates of model quality, we performed nested cross-validation by randomly choosing training and test set 10 times and calculating median accuracy of the model across iterations (Extended Data Fig. 4A). Log-log transformation of age resulted in a higher quality of the transcriptomic clock (median Pearson’s r = 0.938, median MAE = 2.11 months), while scaling and YuGene normalization methods produced comparable accuracy.

**Fig. 2.**
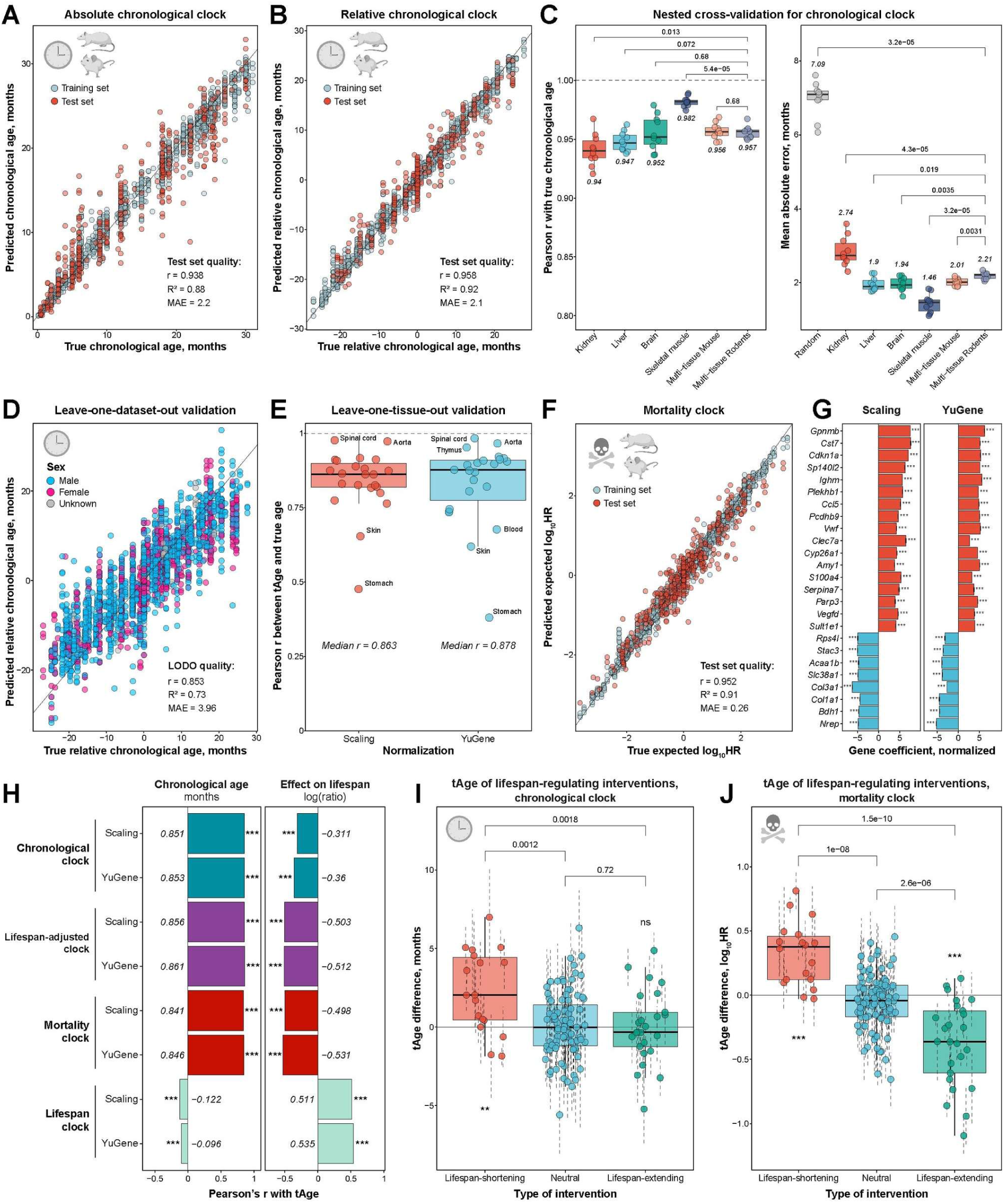
Multi-tissue transcriptomic clocks capture mortality-associated changes caused by aging and lifespan-regulating interventions. **A-B.** Accuracy of prediction of absolute (A) and relative (B) chronological age with multi-tissue transcriptomic clocks trained with Elastic Net (EN) model on mice and rats (with scaling normalization). Training and test sets are denoted by color. Pearson correlation coefficient, R^2^ and mean absolute error (MAE) for test set are shown in the text. **C.** Accuracy of predictions of relative chronological age on 10 randomly chosen test sets for tissue-specific (kidney, liver, brain, skeletal muscle) and mouse or rodent multi-tissue EN transcriptomic clocks. Pearson correlation coefficient and MAE are shown on left and right panels, respectively. Median estimate of quality across 10 runs is provided in text. MAE of random age prediction is shown in grey. Pairwise comparison of accuracy for various models was performed with Wilcoxon signed-rank test, corresponding BH-adjusted p-values are shown in text. **D.** Leave-one-dataset-out (LODO) accuracy of relative chronological age prediction with rodent multi-tissue EN transcriptomic clock (with YuGene normalization). Sex of samples is depicted with color. Pearson correlation coefficient, R^2^ and mean absolute error (MAE) are shown in the text. **E.** Quality of relative chronological age prediction for novel tissues not included in the training set with rodent multi-tissue EN transcriptomic clock. Pearson correlation between real and predicted age was estimated for each organ after training on remaining tissues (leave-one-tissue-out procedure) with scaling (left) and YuGene (right) normalization methods. Median correlation coefficient across tissues is shown in the text. **F.** Accuracy of prediction of relative expected hazard ratio (HR) in log scale with rodent multi-tissue EN transcriptomic clock (with YuGene normalization). Training and test sets are denoted by color. Pearson correlation coefficient, R^2^ and mean absolute error (MAE) for test set are shown in the text. **G.** Top 25 genes with the highest average absolute coefficients in rodent multi-tissue EN mortality clocks trained on data subjected to scaling (left) or YuGene (right) normalization across 10 randomly chosen training sets. Data are mean normalized coefficients ± SE. **H.** Quality of prediction of chronological age (left) and effect on lifespan induced by various interventions (right) for rodent multi-tissue EN transcriptomic clocks of chronological age, lifespan-adjusted age, mortality and expected maximum lifespan. Pearson correlation coefficient was calculated between predicted transcriptomic age and real chronological age (months) or effect on lifespan (log-ratio between treated and control animals) in independent datasets. Median Pearson’s r and statistical significance are shown with text and asterisks, respectively. **I-J.** Accuracy of prediction of intervention effect on expected lifespan for rodent multi-tissue EN transcriptomic clocks of chronological age (I) and mortality (J) (with YuGene normalization). For every type of interventions (lifespan-shortening, neutral or lifespan-extending), dots reflect mean tAge differences between treated and control samples for a given intervention, dataset and sex. Statistical significance of deviation from zero for lifespan-shortening and lifespan-extending interventions and pairwise comparison of tAges across the groups were assessed with mixed effect models. Corresponding BH-adjusted p-values are shown with asterisks and text, respectively. Data are mean tAge differences ± SE. * p.adj < 0.05; ** p.adj < 0.01; *** p.adj < 0.001.

To further improve predictive power of the clock by reducing batch effect, we calculated relative gene expression compared to a randomly chosen reference group within each dataset and tissue, and trained the clock to predict difference in chronological age based on differences in gene expression profiles (see Methods; Fig. 2B). Indeed, this approach resulted in an even higher R^2^ on test sets (median Pearson’s r = 0.957 and median R^2^ = 0.92 across 10 iterations) (Fig. 2C; Extended Data Fig. 4A), demonstrating quality of age prediction comparable to epigenetic clocks^37,39,40,49^. Finally, to enhance applicability of the model, we trained it with Bayesian Ridge (BR) model that provides transcriptomic age (tAge) point estimate together with a credible interval (CI) reflecting the level of uncertainty for model’s prediction on the given data, which may be particularly useful when model is applied to novel interventions and datasets with small sample sizes. The accuracy of BR clocks was similar to EN models (median Pearson’s r = 0.953, median R^2^ = 0.91) (Extended Data Fig. 4A), suggesting that they can also be employed for assessing aging-associated molecular phenotypes.

In addition to the rodent multi-tissue model, we trained chronological clocks on individual tissues, including kidney, liver, brain and skeletal muscle, as well as a multi-tissue model based only on mouse samples (Fig. 2C; Extended Data Fig. 4B). The resulting clocks had comparable accuracy (median Pearson’s r between 0.94 and 0.982), suggesting that individual tissues may be integrated in a multi-tissue model without substantial loss of quality. We also applied other machine learning algorithms, including Support Vector Machines (SVM), Light Gradient Boosting Machine (LightGBM), Random Forest and K-Nearest Neighbors (KNN), and observed similar or worse accuracy compared to the EN model (Extended Data Fig. 4C).

To validate the clock’s ability to predict chronological age of samples from independent datasets, we performed leave-one-dataset-out (LODO) test, iteratively excluding each dataset from the training set and passing it to the model trained on remaining datasets. In agreement with previous results, we observed higher quality of age prediction across independent datasets for the relative chronological clock (Pearson’s r = 0.853, MAE = 3.96 months; Fig. 2D), which significantly outperformed absolute clocks (Extended Data Fig. 4E-F), confirming that differentiation of gene expression profiles is effective for the reduction of tissue- and dataset-associated batch effects.

To examine if the multi-tissue model can be applied to tissues not used during training, we performed leave-one-tissue-out test. Remarkably, relative chronological clock was able to predict age dynamics in all tested unseen tissues (Extended Data Fig. 4D) with the median Pearson’s r = 0.878 (Fig. 2E), suggesting that the model indeed captures systemic aging-associated molecular hallmarks shared across mammalian organs.

To develop an integrated model encompassing molecular mechanisms of aging and longevity, we utilized the complete transcriptomic meta-dataset that includes healthy animals and rodents subjected to lifespan-shortening, neutral or longevity interventions together with the corresponding survival data (Fig. 1I-J). Using these data, we trained rodent multi-tissue transcriptomic clocks of lifespan-adjusted age and expected mortality rate (Fig. 2F; Extended Data Fig. 5A). Like chronological clocks, relative models demonstrated higher quality on test sets than absolute clocks (Median Pearson’s r = 0.954 and 0.943 for lifespan-adjusted age and mortality, respectively), while EN and BR models showed comparable accuracy (Extended Data Fig. 5D-G). When applied to independent datasets in LODO test, multi-tissue transcriptomic clocks of lifespan-adjusted age and mortality also showed similar quality (Pearson’s r = 0.848 and 0.81, respectively; Extended Data Fig. 6A-B).

In agreement with the signature analysis, clock coefficients were in general positively correlated across models of chronological age, lifespan-adjusted age, and expected mortality (Extended Data Fig. 5B-C), with lifespan-adjusted and mortality clocks exhibiting the highest similarity. Among top features with positive coefficients selected by chronological and mortality models across 10 random iterations, we observed genes *Gpnmb*, *Cst7*, and *Cdkn1a*, encoding a type I transmembrane glycoprotein, endosomal/lysosomal cathepsin inhibitor, and cell cycle inhibitor, respectively (Fig. 2G; Extended Data Fig. 4G). Remarkably, these genes have been previously shown to be prognostic biomarkers of individual diseases. Thus, glycoprotein nonmetastatic melanoma protein B (GPNMB), involved in regulation of cell differentiation, regeneration and inflammation^72^, was shown to have elevated levels in patients with neurodegenerative diseases, including Alzheimer’s disease^73^, Gaucher disease^74^, and amyotrophic lateral sclerosis^75^. Cystatin F regulates cell cytotoxicity, while its induction is also associated with development of brain pathologies, such as a prion disease^76^ and AD in mice and humans^77^, presumably through the impairment of lysosomal function in microglia. Finally, cyclin-dependent kinase inhibitor p21 (CDKN1A), a member of the p53 pathway, is a well-established marker of cellular senescence, DNA damage, and tumor progression^78,79^. Interestingly, *Cst7* and *Cdkn1a* expression across rodent tissues demonstrated positive association with both aging and mortality after adjustment for age (Extended Data Fig. 3E), suggesting that these factors are robust molecular hallmarks of impaired health. In contrast, top features with negative coefficients in mortality clock included several genes associated with cellular differentiation and epithelial-mesenchymal transition (EMT), such as *Nrep*, *Col1a1* and *Col3a1* (Fig. 2G), presumably reflecting the overall stem cell exhaustion observed with aging and in short-lived mouse models^80,81^. Thus, neuronal regeneration related protein (NREP, of P311) plays a crucial role in wound healing and scar formation^82^. Accordingly, expression of these genes showed significant negative association with both chronological age and mortality after adjustment for age (Extended Data Fig. 3E).

To test if the developed molecular models of biological age capture both the effects of aging and lifespan-regulating interventions, we assessed the correlation of transcriptomic clock predictions on independent datasets separately with chronological age and with the ratio of expected maximum lifespan between age-matched control animals and mice subjected to the intervention. To have a positive control for the second test, we trained gene expression model to directly predict lifespan (i.e., lifespan clock), utilizing changes in expected maximum lifespan as a response variable (Extended Data Fig. 6C). Interestingly, addition of chronological age to the list of features for this model didn’t improve its quality (median Pearson r across test sets is 0.885 and 0.886 for lifespan model with and without chronological age, respectively), suggesting that chronological age differences were well reflected by gene expression data (Extended Data Fig. 6D). As expected, predictions of chronological clocks demonstrated a high positive correlation with age and a mild negative correlation with the effect on maximum lifespan (Pearson r ≈ 0.85 and −0.36, respectively), whereas the lifespan clock showed the opposite (Pearson r ≈ −0.12 and 0.54, respectively) (Fig. 2H; Extended Data Fig. 6E). At the same time, lifespan-adjusted and mortality clocks demonstrated high correlation with both chronological age and the effect on lifespan (Pearson r ≈ 0.85 and −0.53, respectively). Remarkably, when we analyzed short-lived and long-lived models separately, we observed that the multi-tissue chronological clock was able to distinguish detrimental interventions from other groups (p.adjusted = 0.0012) and predicted higher tAge for these models (p.adjusted = 0.003) (Fig. 2I). However, it didn’t capture the effect of lifespan-extending interventions (p.adjusted > 0.7), in agreement with previous studies showing that established longevity interventions typically don’t produce a strong anti-aging effect on the gene expression profile^28,36^. In contrast, transcriptomic clocks of lifespan-adjusted age and especially clocks of expected mortality distinguished both short-lived and long-lived models (p.adjusted < 6.10^−4^ and < 3.10^−6^ for lifespan-adjusted and mortality clocks, respectively), predicting higher and lower tAge for detrimental and longevity interventions, respectively (p.adjusted < 6.9.10^−5^ and < 1.3.10^-5^ for lifespan-adjusted and mortality clocks, respectively) (Fig. 2J; Extended Data Fig. 6F). Remarkably, the mortality clock predicted lifespan-modulating models even better than lifespan clock (Extended Data Fig. 6G), suggesting that aging and longevity-affecting models complement each other improving the overall quality of molecular biomarkers of health.

### Co-regulated gene expression modules of aging and longevity

To deconstruct transcriptomic hallmarks of aging and longevity into independent co-regulated components, we employed weighted gene co-expression network analysis (WGCNA)^83^. To focus on modules associated with aging- and lifespan-associated changes and reduce the impact of batches, tissues, and sexes, we centered all gene expression profiles in our meta-dataset around median profile of randomly selected reference group from the same dataset, tissue, and sex (see Methods). Genes in the resulting dataset formed several distinct co-regulated clusters (Extended Data Fig. 7A), suggesting that the signatures of aging and longevity may be separated into components. WGCNA revealed 28 co-regulated modules, each composed of 30 to 630 genes (Fig. 3A; Extended Data Fig. 8). Most modules identified by separately analyzing male and female samples demonstrated significant overlap, confirming that gene clustering was largely not driven by sex-specific differences in gene expression (Extended Data Fig. 7B). Following filtering of genes within each module (see Methods; Extended Data Fig. 7C), we performed a functional enrichment analysis. Remarkably, we found that most gene clusters were enriched for specific sets of pathways, with very little overlap across modules (Fig. 3B; Extended Data Fig. 9-10; Supplementary Table 4). Thus, turquoise, blue, and greenyellow modules were enriched for genes associated with inflammatory response and innate immune system, mitochondrial translation and respiratory electron transport, and extracellular matrix (ECM) organization and EMT, respectively. To validate the biological meaning of the identified gene network components, we estimated partial correlation between 1^st^ principal components (PCs) of modules after adjustment for chronological age (Extended Data Fig. 7D). As expected, modules annotated with similar functions had a higher correlation and clustered together (Fig. 3C). For example, modules associated with inflammatory response, interferon signaling and T cell signaling formed a connected higher-order cluster, while metabolism-related modules were organized in a different interconnected cluster. To examine the association of transcriptomic components with aging and longevity, we estimated correlation of their 1^st^ PCs with chronological age and expected maximum lifespan (Fig. 3C-D). In agreement with our previous studies, modules of immune response demonstrated a strong positive association with aging and a negative correlation with maximum lifespan^28,33^. In contrast, modules of mitochondrial function, oxidative phosphorylation, and lipid metabolism were positively correlated with longevity and negatively associated with age, highlighting their role in the regulation of healthspan^33,35,66,84,85^. Overall, most modules demonstrated opposite associations with aging and lifespan. However, several transcriptomic gene sets, including modules of heat stress response, translation, and ECM organization and EMT, showed co-directional associations with chronological age and maximum lifespan, potentially reflecting adaptive aging-related hallmarks. Finally, some modules, such as those involved in protein processing and mTOR signaling, demonstrated association with longevity but no significant correlation with aging (Fig. 3D).

**Fig. 3.**
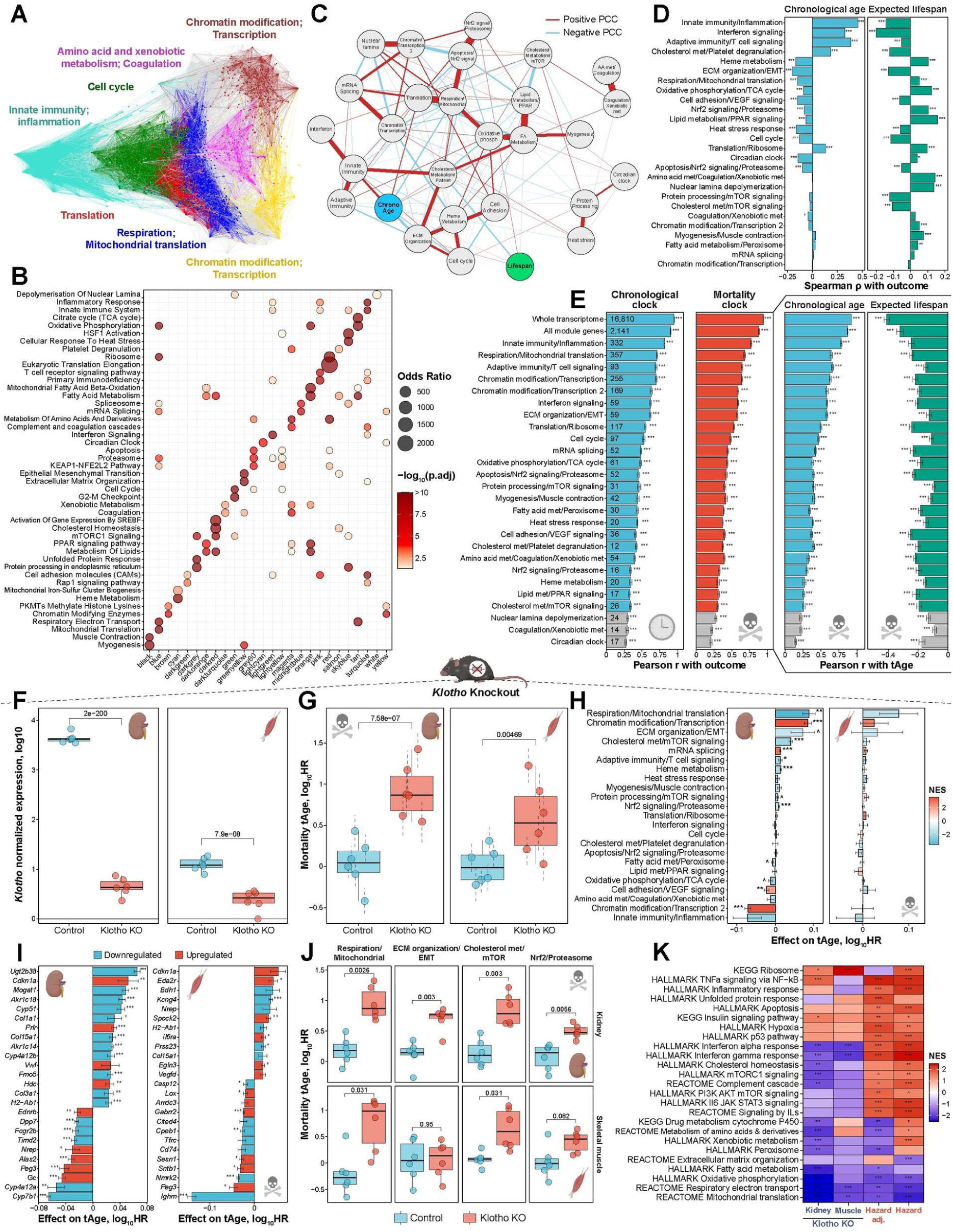
Co-regulated gene expression modules characterize components of mammalian aging and longevity. **A.** Gene co-expression network based on spectral embedding with several modules identified with weighted gene co-expression network analysis (WGCNA). Representative enriched functions for each module are shown in the text with the corresponding color. **B.** Functional enrichment of co-regulated gene expression modules identified with WGCNA. Modules and pathways are shown in columns and rows, respectively. Size of dots and their color reflect odds ratio and BH-adjusted p-value, respectively. Only enrichments with p.adjusted < 0.05 are shown with dots. The whole list of enriched functions is in Supplementary Table 4A. **C.** Partial correlation network of 1^st^ principal components (PCs) of co-regulated modules (grey dots), chronological age (blue) and expected maximum lifespan (green). Sign and scale of partial correlation coefficient (PCC) are indicated by color and line width. Modules are named after representative enriched functions. The dictionary between modules and functions is in Supplementary Table 4B. AA: Amino acid; Chrono: Chronological; FA: Fatty acid. **D.** Spearman correlation between 1^st^ PC of module and chronological age (blue) or expected maximum lifespan after adjustment for age (green). Statistical significance is reflected with asterisks. **E.** Quality of rodent multi-tissue Elastic Net (EN) transcriptomic clocks of chronological age (blue) and expected mortality (red) trained on various co-regulated modules (left), and correlation of mortality clock predictions with chronological age (blue) and expected maximum lifespan adjusted for age (green) (right) on 10 randomly chosen test sets. All available genes (1^st^ row), all genes associated with at least one module (2^nd^ row) and genes associated with individual modules (other lines) were used to train the clocks. Number of genes included in feature set for each clock is shown with text. Statistical significance is denoted with asterisks. Module clocks with Pearson correlation coefficient with outcome < 0.3 are shown in grey and excluded from the subsequent analysis. **F.** Normalized expression of *Klotho* (in log scale) in kidney (left) and skeletal muscle (right) of 8-week-old control mice and age-matched mice with *Klotho* knockout (KO). BH-adjusted p-value is shown in text. **G.** Transcriptomic age (tAge) for control and *Klotho* KO mice in kidney (left) and muscle (right) estimated with rodent multi-tissue Bayesian Ridge (BR) mortality clock trained on all genes excluding *Klotho*. tAges between the groups were compared with mixed effect model, corresponding BH-adjusted p-values are shown in text. Data are tAges ± SE. **H.** Contributions of gene expression modules to tAge difference between control and *Klotho* KO mice estimated with the rodent multi-tissue Elastic Net (EN) mortality clock. Individual modules are shown in rows and named after representative functions. Positive and negative values reflect pro- and anti-mortality changes in *Klotho* KO mice compared to age-matched controls, respectively. Statistical significance for each module was assessed with the two-sample unpaired t-test and indicated with asterisks. Bars are colored based on normalized enrichment scores (NES) from gene set enrichment analysis (GSEA), reflecting if genes associated with a particular module are generally up-(red) or downregulated (blue) in *Klotho* KO mice. Modules significantly enriched for up- or downregulated genes (BH-adjusted p-value < 0.05) are visualized with thick bars. Data are means ± SE. **I.** Top genes driving pro-(positive) or anti-mortality (negative) transcriptomic changes in kidneys (left) and skeletal muscles (right) of *Klotho* KO mice compared to age-matched controls, according to the rodent multi-tissue EN clock. Top 25 genes with the highest absolute effect on tAge difference (logFC * clock coefficient) are shown. Genes up- and downregulated in *Klotho* KO mice are colored in red and blue, respectively. Statistical significance of logFC for each gene is indicated with asterisks. Data are mean tAge difference ± SE. **J.** Mortality tAge in control and *Klotho* KO mice estimated with representative module-specific multi-tissue mortality clocks. Difference between the groups was assessed with the two-sample unpaired t-test, and corresponding BH-adjusted p-values are shown in text. Results for all module-specific mortality and chronological clocks are in Extended Data Fig. 11G. **K.** Functional enrichment (GSEA) of gene expression changes induced in *Klotho* KO mice, and signatures of mortality. Only functions significantly enriched by at least one signature are shown (BH-adjusted p-value < 0.05). The whole list of enriched functions is in Supplementary Table 5A. HR: Hazard Ratio; ECM: Extracellular matrix; EMT: Epithelial-Mesenchymal Transition; met: metabolism. ^ p.adj < 0.1; * p.adj < 0.05; ** p.adj < 0.01; *** p.adj < 0.001.

To develop quantitative biomarkers of mortality based on the identified independent components of gene expression network, for every module we trained relative rodent multi-tissue transcriptomic clocks of chronological age and expected mortality (Fig. 3E). Clocks trained on all genes included in at least one module (2,141 genes) demonstrated high quality comparable to the clocks trained on the whole transcriptome (median Pearson r across test sets = 0.88-0.9), suggesting that together the identified co-regulated components explain most of the association between gene expression and mortality. Most individual module-specific clocks were able to predict response variables in test sets with moderate accuracy (median Pearson’s r across modules = 0.41-0.44). After filtering out modules with median Pearson’s r < 0.3, we obtained 23 functionally annotated module-specific multi-tissue clocks of chronological age and mortality (Fig. 3E, left). Moreover, all mortality clocks were positively correlated with chronological age and negatively correlated with expected maximum lifespan in test samples (Fig. 3E, right), confirming that mortality models trained on gene sets from individual modules preserved the composite nature of this metric. Besides, transcriptomic ages predicted with module-specific clocks maintained the overall structure of the network observed at the level of the 1^st^ PCs (e.g., outputs of clocks trained on inflammatory, adaptive immunity, and interferon signaling modules formed an interconnected cluster) (Extended Data Fig. 7E-F), suggesting that predictions of the clocks indeed represent mortality-associated changes of specific cellular pathways and may be used to obtain interpretable data on molecular mechanisms of aging and longevity in the given animal model.

### Molecular mechanisms of mortality induced by *Klotho* knockout

To validate the developed biomarkers and characterize molecular mechanisms of mortality associated with an established progeria model, we performed RNA-seq on the kidney and skeletal muscle samples of *Klotho* gene knockout mice and healthy age-matched controls (n=6 per group). *Klotho* is an established pro-longevity gene involved in several lifespan-associated pathways. It was shown to inhibit IGF1/mTOR signaling, upregulate antioxidant enzymes through activation of Nrf2 signaling, and block TGF-β leading to prevention of fibrosis and stem cell exhaustion^86^. *Klotho* overexpression extends mouse lifespan^87^, whereas its deficiency results in progeria, limiting mouse maximum lifespan to 4-5 months^6,7^.

*Klotho* was highly expressed in the kidneys of control mice (Fig. 3F), consistent with previous data^86^, while its expression was barely detected in both kidneys and skeletal muscles of KO mice. To get an unbiased estimate of transcriptomic age, we retrained multi-tissue clocks by excluding *Klotho* gene from the feature set and applied the models to control and progeria samples. In agreement with survival data, we detected a statistically significant increase of transcriptomic age in both organs of *Klotho* KO mice, according to chronological, lifespan-adjusted, and mortality clocks (Fig. 3G; Extended Data Fig. 11A-B). Interestingly, a more profound effect was observed in kidneys where the gene is usually produced (Fig. 3G). Among top genes driving the pro-aging and pro-mortality effect of this model was *Cdkn1a*, upregulated in both tissues (Fig. 3I; Extended Data Fig. 11C-D), consistent with the role of Klotho in the prevention of cellular senescence^86^. Overall, gene expression changes contributing to a higher transcriptomic age were similar between kidneys and skeletal muscles (Pearson’s r = 0.39) (Extended Data Fig. 11D).

We then examined top transcriptomic modules that contributed to the increased molecular damage in *Klotho* KO mice after adjustment for the effect of all other modules. To do that, we applied EN multi-tissue clocks trained on all genes included in at least one module and summed up contributions of each module for control and *Klotho* KO samples, followed by pairwise comparison between the groups. Top modules driving the pro-mortality effect of this progeria model in kidneys included those associated with respiration and mitochondrial translation, ECM organization and EMT, cholesterol metabolism and mTOR signaling, and Nrf2 signaling and proteasome function (Fig. 3H), in agreement with the established mechanisms of Klotho activity from the previous studies^86^. Respiration/mitochondrial translation and ECM/EMT modules also had the highest effect on increased tAge in the muscle, according to chronological and mortality clocks (Extended Data Fig. 11E). Interestingly, the module of inflammation and innate immunity was downregulated in both kidneys and skeletal muscles of *Klotho* KO mice, and accordingly had a negative effect on mortality and chronological tAge, suggesting that this cellular pathway was not responsible for the pro-mortality phenotype in this progeria model (Fig. 3H; Extended Data Fig. 11E).

To identify all functional components exhibiting the damaging effect of *Klotho* KO, we applied module-specific multi-tissue chronological and mortality clocks. In agreement with previous results, clocks trained on respiration and mitochondrial translation, and cholesterol metabolism and mTOR signaling gene sets demonstrated a statistically significant increase of tAge in both tissues according to chronological and mortality clocks (Fig. 3J; Extended Data Fig. 11F-G), confirming that these cellular pathways are shared hallmarks of this progeria model across organs. Besides, we observed a statistically significant pro-mortality effect for ECM organization and EMT, and Nrf2 signaling and proteasome modules in kidneys (Fig. 3J), while skeletal muscle was characterized by a significantly aged profile of myogenesis and muscle contraction component (Extended Data Fig. 11F-G). This data suggests that *Klotho* KO is characterized by both shared mechanisms of aging across organs and tissue-specific effects highlighted by co-regulated modules of the gene network. Again, immunity-related modules didn’t exhibit increased tAge in the *Klotho* KO model (Extended Data Fig. 11G), although they were among most accurate module-specific clocks (Fig. 3E), suggesting that inflammation and interferon signaling were not elevated in the examined progeria model. Finally, to establish the direction of expression changes for the identified mortality-associated pathways in *Klotho* KO mice, we performed functional enrichment (Fig. 3K; Supplementary Table 5A). In agreement with the module analysis, we observed a strong statistically significant downregulation of genes associated with respiratory electron transport and mitochondrial translation induced by *Klotho* KO in both organs, and these pathways were also negatively correlated with mortality, both before and after adjustment for chronological age, providing further evidence for impairment of mitochondrial function induced by this genetic intervention. Thus, the identified co-regulated components of the gene expression network provide interpretable characterization of molecular mechanisms responsible for the observed change of biological age induced by an intervention.

### Transcriptomic aging of individual cell types

To examine if multi-tissue transcriptomic hallmarks identified in bulk tissues reflect molecular mechanisms of aging and mortality within individual cell types, we utilized murine droplet scRNA-seq data collected by Tabula Muris Consortium^88^. This dataset includes gene expression profiles of single cells obtained from various tissues of mice of different age and sex. To increase coverage of single cell data for the clock application, we employed a metacell approach, pooling read counts from a randomly selected number of cells corresponding to the same biological sample (Fig. 4A). To validate this method, we selected tissues with at least 7 samples covering more than 15 months of mouse lifespan, pooled together reads from all available cells for every tissue and animal and applied the mouse multi-tissue chronological clock (Extended Data Fig. 12A-B). The clock was able to capture age dynamics with high quality for all examined mouse organs (median Pearson’s r = 0.92; Pearson’s r for pooled data = 0.86). To test how metacell coverage affects clock quality, we reran the analysis using different numbers of cells for metacell aggregation (Fig. 4B-C). Dependence between cell coverage and clock quality followed a saturation model, reaching median Pearson’s r = 0.9 (95% of the maximum value) starting from approximately 100 cells (∼1 mln reads per metacell), while already 25 cells (200-250 thousand reads per metacell) resulted in median Pearson’s r = 0.84, corresponding to 89% of the maximum value.

**Fig. 4.**
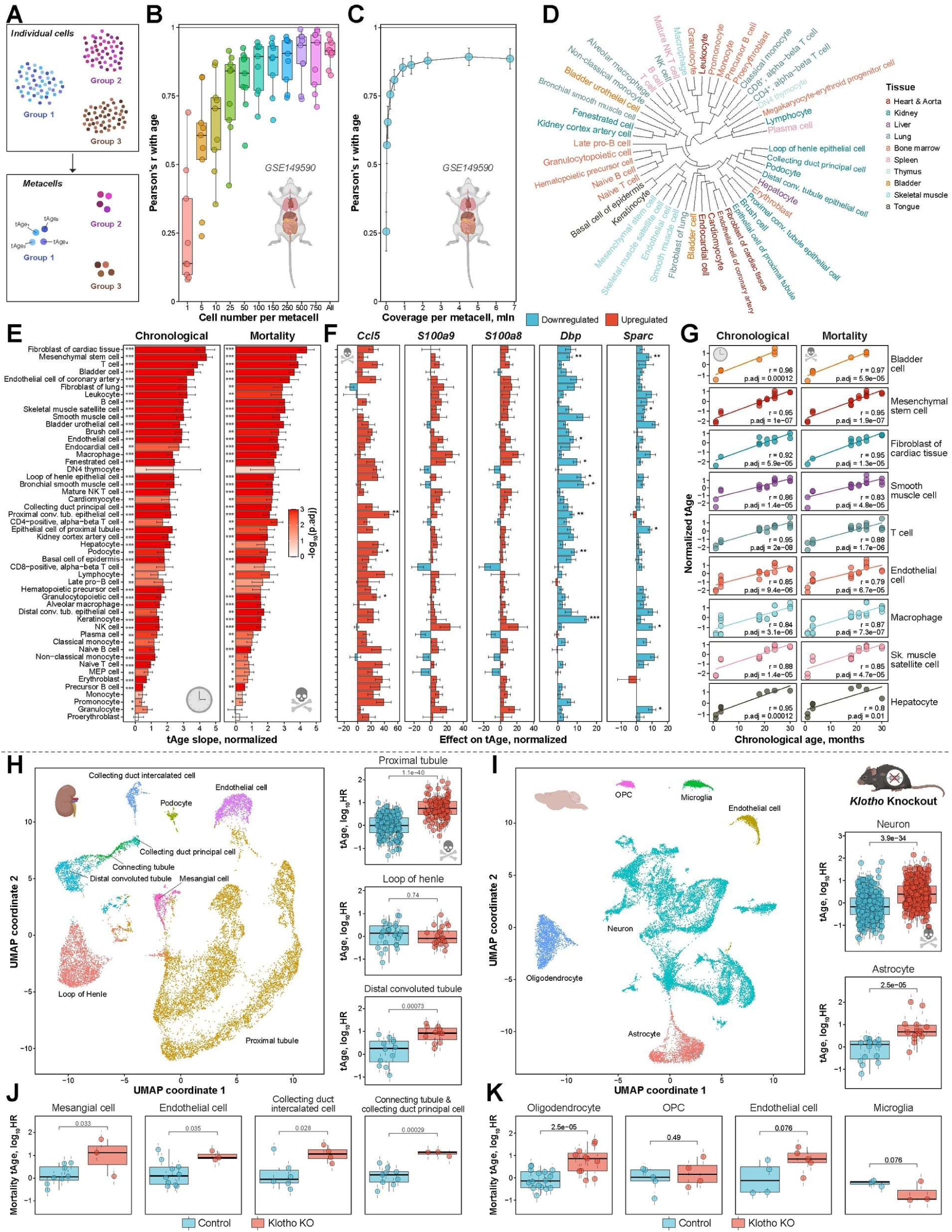
Single-cell RNA-seq data reveal conserved mechanisms of aging and mortality across cell types in naturally aged and short-lived mice. **A.** Scheme of a metacell analysis of transcriptomic age of individual cells. Gene expression profiles of randomly chosen cells corresponding to the same type and experimental group are pooled together and passed to clocks. **B.** Dependence between quality of age prediction with mouse multi-tissue Elastic Net (EN) chronological transcriptomic clock and number of cells from a sample aggregated in a single metacell. Dots reflect individual tissues from the droplet Tabula Muris Senis scRNA-seq dataset. **C.** Dependence between quality of chronological age prediction with mouse multi-tissue EN chronological clock and mean coverage of aggregated metacells. Data are mean Pearson correlation coefficient between chronological ages and predicted transcriptomic ages (tAges) across tissues in Tabula Muris Senis data ± SE. **D.** Hierarchical clustering of weighted aging-associated gene expression signatures of individual cell types. Union of top 1000 genes associated with age (with the lowest p-value) was used to estimate Pearson’s correlation coefficient for each pair of cell types. Weighted signatures were calculated as (age slope * chronological clock coefficient). Complete hierarchical method based on correlation distance was used for clustering. Cell types are colored according to tissue origin. **E.** Dependence between chronological age and transcriptomic age estimated with mouse multi-tissue EN chronological (left) and mortality (right) clocks for individual cell types from Tabula Muris Senis dataset (in rows). Data are normalized slopes ± SE. Statistical significance of association was assessed with linear model. Asterisks reflect corresponding BH-adjusted p-values. MEP: Megakaryocytic-erythroid progenitors. **F.** Top genes contributing into increased molecular age across cell types according to mouse multi-tissue EN mortality clock. Top 5 genes with the highest average effect on normalized tAge (normalized slope * clock coefficient) are shown. Cell types are labelled on E. Genes up- and downregulated with age are colored in red and blue, respectively. Statistical significance of slope for each gene is denoted with asterisks. Data are normalized tAge slope ± SE. **G.** Molecular age dynamics in representative cell types estimated with mouse multi-tissue chronological (left) and mortality (right) clocks. Pearson correlation coefficient and corresponding BH-adjusted p-value are shown in text. Sk. muscle: Skeletal muscle. **H-I.** UMAP of filtered kidney (H) and brain (I) cells from 8-week-old control and *Klotho* knockout (KO) mice. Cell type annotation is denoted with color. OPC: Oligodendrocyte Precursor Cells. **J-K.** Mortality tAge difference between metacells of control and age-matched *Klotho* KO mice representing various kidney (J) and brain (K) cell types, estimated with the rodent multi-species Bayesian Ridge (BR) mortality clock trained on all genes excluding *Klotho*. tAges between the groups were compared with mixed effect model, and BH-adjusted p-values are shown in text. Data are tAges ± SE. HR: Hazard Ratio. * p.adj < 0.05; ** p.adj < 0.01; *** p.adj < 0.001.

To examine if multi-tissue biomarkers of chronological age and mortality capture age dynamics within individual cell types, we applied the metacell procedure and transcriptomic clocks separately to every cell type. Interestingly, out of 50 examined cell types (Fig. 4D), 47 and 46 of them demonstrated a statistically significant increase of tAge with mouse age according to chronological and mortality clocks, respectively (Fig. 4E, Extended Data Fig. 12C). They included fibroblasts, hepatocytes, muscle cells, endothelial cells, macrophages, lymphocytes, etc. (Fig. 4G). Surprisingly, even stem cells, such as mesenchymal stem cells and skeletal muscle satellite cells, exhibited substantial pro-aging and pro-mortality gene expression changes with age, suggesting that aging is associated with systemic universal molecular changes shared across most cell types and tissues.

Indeed, gene expression changes contributing to increased tAge with time were generally positively correlated across cell types (Extended Data Fig. 13A). Top shared pro-aging and pro-mortality transcriptomic hallmarks included upregulation of several inflammation-associated genes, such as a pro-inflammatory chemokine *Ccl5*^89^ and modulators of leukocyte recruitment and cytokine secretion *S100a8* and *S100a9*^90^, as well as downregulation of the circadian rhythm regulator *Dbp*^91–93^ and a modulator of ECM synthesis, differentiation, and wound healing *Sparc*^94^ (Fig. 4F; Extended Data Fig. 13B). Despite the overall similarity in aging-associated gene expression changes across cell types, clustering revealed several groups of cell types that shared closely related aging profiles (Fig. 4D). In general, cell types aggregated according to their tissue origin, with the exception of blood cells, most of which formed a single cluster even though they were derived from different organs. This suggests that aging is accompanied by both global hallmarks of damage accumulation shared across tissues and local environment effects specific for neighboring groups of cells. Detailed exploration of differences in aging across mammalian cell types may provide insights for the development of effective longevity therapies targeted to specific subsystems of the organism.

### Effect of *Klotho* KO on gene expression hallmarks of mortality across cell types

To further characterize molecular mechanisms of the *Klotho* KO progeria model and test if it induces systemic pro-aging effect shared across multiple cell types, we performed single-nucleus RNA-seq (snRNA-seq) of kidney and brain tissues from an 8-week-old control mouse and an age-matched *Klotho* KO animal. UMAP revealed distinct clusters of cells in each organ, representing various cell types (Fig. 4I; Extended Data Fig. 14A). The brain was mostly composed of neurons, along with astrocytes, oligodendrocytes, microglia, endothelial cells, and oligodendrocyte precursor cells (OPC) (Fig. 4I). The majority of kidney cells were proximal tubule epithelial cells, accompanied by mesangial cells, endothelial cells, podocytes, epithelial cells from loop of Henle, distal convoluted tubule, and connecting tubule, as well as principal and intercalated cells of collecting duct (Extended Data Fig. 14A). To decrease noise and chances of getting a mixed signal from multiple cell types, we filtered kidney data ensuring that cells from every annotated cell type form a single cluster on the UMAP (Fig. 4H). Interestingly, most kidney cell types demonstrated a clear separation between control and *Klotho* KO mice, while brain cells formed homogeneous clusters (Extended Data Fig. 14B-C), confirming that the kidney is more affected by this genetic manipulation. Indeed, we observed a substantially higher snRNA level of *Klotho* in kidney cells of control mice, and it was significantly downregulated in all cell types of progeroid mice (Extended Data Fig. 14D). In contrast, brain cells exhibited lower expression of this gene in control animals, and its downregulation in *Klotho* KO mice was statistically significant only for neurons and oligodendrocytes, although a similar trend was observed for all cells except for OPC (Extended Data Fig. 14E).

We then aggregated metacells separately for each cell type of control and *Klotho* KO mice using the total coverage of 200K reads per metacell and applied multi-tissue chronological and mortality clocks trained on all genes except for *Klotho*. The mortality clock revealed a statistically significant acceleration of tAge induced by *Klotho* KO in all presented kidney cell types, except for the loop of Henle cells (Fig. 4J). The chronological clock showed similar effects for most cell types, although not all of them reached statistical significance (Extended Data Fig. 14F). In the brain, both mortality and chronological clocks detected a significant increase of tAge in neurons, astrocytes, and oligodendrocytes, as well as marginal significance in endothelial cells (p.adjusted = 0.076-0.079) (Fig. 4K; Extended Data Fig. 14G). Interestingly, microglia cells demonstrated a marginally significant decrease of tAge according to the mortality clock (p.adjusted = 0.076), consistent with the previous data showing that *Klotho* KO results in systemic downregulation of genes associated with inflammatory response across organs (Fig. 3H,K).

To further characterize molecular mechanisms of the progeria, we focused on the most representative cell types in the kidney and brain, being proximal tubule cells and neurons, respectively. Genes driving aging- and mortality-associated effect of *Klotho* KO in proximal tubule cells were similar to those in the bulk kidney (Pearson’s r = 0.57) (Extended Data Fig. 15A-C), including upregulated *Cdkn1a* and *Cp,* the latter encoding ceruloplasmin glycoprotein involved in iron metabolism and shown to be elevated in multiple disease models, including HCC, glioma, heart failure, and cerebral ischemia^95–98^. At the same time, although neurons also exhibited an acceleration of tAge in *Klotho* KO model, it was associated with distinct molecular signatures of mortality, not significantly correlated with those in proximal tubules (Pearson’s r = 0.02) and weakly correlated with those in skeletal muscle (Pearson’s r = 0.11) (Extended Data Fig. 15A-C). Among top gene drivers of the pro-mortality molecular phenotype of neurons, we detected downregulated *Nrep* and upregulated cell cycle regulator *Junb*, capable of inducing senescence and stem cell niche delpetion^99,100^.

Despite distinct mortality-associated signatures of *Klotho* KO at the level of individual genes, our module analysis revealed several shared molecular mechanisms of progeria between proximal tubule cells and neurons (Extended Data Fig. 15D-E). Thus, modules of respiration and mitochondrial translation, and cholesterol metabolism and mTOR signaling were among top contributors to biological age acceleration in both cell types, in agreement with bulk transcriptomic data. At the same time, the inflammatory module had a negative effect on mortality tAge in both proximal tubule cells and neurons, and the corresponding genes were significantly downregulated, especially in neurons. Accordingly, mortality tAge of neurons and proximal tubules estimated with the module-specific inflammatory clock also demonstrated a trend towards decline in progeroid mice (Extended Data Fig. 15F). These findings are in line with the observed decreased biological age of microglia cells in *Klotho* KO mice (Fig. 4K), suggesting that the overall inflammatory response was diminished by introduced genetic manipulation. Therefore, this progeria model shows that lifespan-regulating interventions may produce contrasting effects on mortality across different cell types and cellular components, emphasizing the systemic nature of mammalian mechanisms of aging and longevity.

### Molecular mechanisms of biological age deviation in models of chronic disease

To examine molecular mechanisms of mortality induced by established models of aging-related diseases, we preprocessed 9 publicly available datasets with gene expression profiles of tissues from animals representing models of Alzheimer’s disease (AD) (brain and pineal gland)^101^, chronic kidney disease (CKD) (kidney)^102^, diabetic neuropathy (DN) (kidney)^103^, ischemic stroke (IS) (heart, brain and microglia cells)^104–106^, nonalcoholic steatohepatitis (NASH) (liver)^107^, and hepatocellular carcinoma (HCC) (liver)^108^, along with age-matched controls. Multi-tissue chronological and mortality clocks revealed a statistically significant acceleration of biological age in all examined disease models and organs, except for HCC (Fig. 5A; Extended Data Fig. 16). Interestingly, transcriptomic age of heart and microglia in the model of ischemic stroke demonstrated an accumulating effect with time following disease induction. Thus, mortality tAge of the heart was elevated already 1 day after the stroke but was even further increased on day 3 (Fig. 5A). The brain, being a primary target of IS, exhibited an even higher increase in biological age 3 days after disease induction, both in young and old animals. At the same time, microglia cells demonstrated an elevated mortality tAge only 14 days after the stroke in both young and old animals, suggesting that various components of the organism may respond to disease models with different rates. Moreover, on day 14 biological age of microglia derived from the ipsilateral hemisphere, where IS was induced, was significantly higher than that from the contralateral hemisphere (p.adjusted < 0.046 for mortality clock) (Fig. 5A), highlighting the effect of the local environment on the dynamics of aging- and mortality-associated molecular mechanisms.

**Fig. 5.**
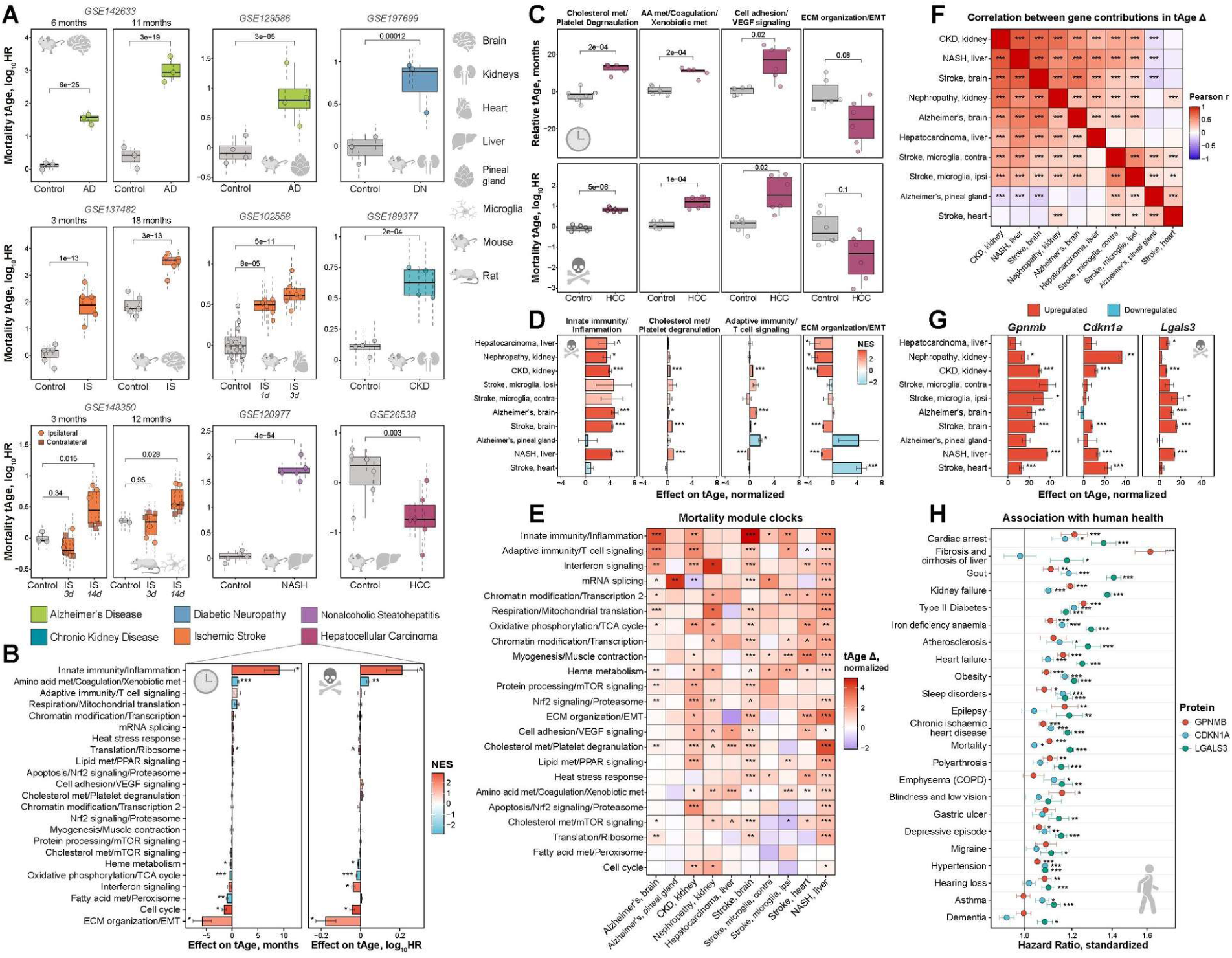
Aging-related diseases in mammals exhibit common molecular and functional changes associated with mortality. **A.** Mortality transcriptomic age (tAge) of tissues from control rodents (grey) and age-matched animals representing different models of age-related diseases (indicated with color), as assessed with the rodent multi-tissue Bayesian Ridge (BR) mortality clock. tAges between the groups were compared with mixed effect model, and corresponding BH-adjusted p-values are shown in the text. Species and organs are depicted with icons. GEO IDs for all datasets are provided in text. Data are tAges ± SE. **B.** Contributions of transcriptomic modules to chronological (left) and mortality (right) tAge difference between livers from control mice and age-matched animals with spontaneous hepatocellular carcinoma, estimated with rodent multi-tissue Elastic Net (EN) chronological and mortality clocks. Individual modules are shown in rows and named after representative functions. Positive and negative values reflect pro- and anti-aging and mortality changes in carcinoma compared to control livers, respectively. Statistical significance for each module was assessed with ANOVA or linear regression (for stroke microglia model) and indicated with asterisks. Bars are colored based on normalized enrichment scores (NES) from gene set enrichment analysis (GSEA), reflecting if genes associated with a particular module are generally up- (red) or downregulated (blue) in carcinoma samples. Modules significantly enriched for up- or downregulated genes (BH-adjusted p-value < 0.05) are visualized with thick bars. Data are means ± SE. **C.** Chronological (up) and mortality (bottom) tAge in control and hepatocellular carcinoma livers estimated with representative module-specific multi-tissue clocks. Difference between the groups was assessed with the two-sample unpaired t-test, and corresponding BH-adjusted p-values are shown in text. **D.** Top modules with the highest absolute contribution to the normalized tAge difference between control and diseased animals across age-related disease models. Color and thickness schemes are identical to (B). **E.** Normalized mortality tAge difference between control and age-matched diseased animals assessed with all module-specific multi-tissue mortality clocks. Color and asterisks reflect size and statistical significance (BH-adjusted p-values) of tAge difference between control and disease groups, assessed with the ANOVA or linear regression (for stroke microglia data). Increased and decreased tAges in disease models are shown in red and blue, respectively. **F.** Correlation between weighted gene expression signatures of mortality (logFC * clock coefficient) across various models of age-related diseases, according to the rodent multi-tissue EN mortality clock. The union of top 500 differentially expressed genes (with the lowest p-value) were used to estimate Pearson’s correlation coefficient for each pair of signatures. Statistical significance is indicated with asterisks. **G.** Top genes contributing to increased molecular age across age-related diseases according to rodent multi-tissue EN mortality clock. Top 3 genes with the highest average effect on normalized tAge (normalized slope * clock coefficient) are shown. Genes up- and downregulated in disease samples are colored in red and blue, respectively. Statistical significance of expression change for each gene is denoted with asterisks. Data are normalized tAge difference ± SE. **H.** Association of GPNMB, CDKN1A and LGALS3 protein concentration in human plasma with mortality and various diseases and risk factors, based on UK Biobank. Hazard Ratio (HR) for protein concentration normalized by standard deviation across patients is shown on x axis. Statistical significance is indicated with asterisks. Data are exp(log_10_HR ± SE). HR: Hazard Ratio; AD: Alzheimer’s disease; DN: Diabetic Neuropathy; IS: Ischemic Stroke; CKD: Chronic Kidney Disease; NASH: Nonalcoholic Steatohepatitis; HCC: Hepatocellular Carcinoma; ipsi: ipsilateral; contra: contralateral; ECM: Extracellular matrix; EMT: Epithelial-Mesenchymal Transition; met: metabolism. ^ p.adj < 0.1; * p.adj < 0.05; ** p.adj < 0.01; *** p.adj < 0.001.

Interestingly, HCC samples showed a significantly lower mortality tAge compared to control liver samples from age-matched animals (Fig. 5A). Many genes driving this signal were related to cellular dedifferentiation, ECM organization and cell cycle, such as *Plekhb1*, *Col1a1*, *Col3a1*, *Col4a1*, *Ube2c*, *Cdk1*, and *Mki67* (Extended Data Fig. 17A-B). Accordingly, modules of EMT/ECM organization and cell cycle had the highest negative statistically significant effect on aging- and mortality-associated molecular profile of HCC (Fig. 5B), confirming that tumors resemble some features of young cells, such as elevated proliferation, dedifferentiation, ECM remodeling, and increased deposition of collagen^109–112^. This was further supported by the module-specific EMT/ECM organization clock that showed a marginally significant decrease of chronological and mortality tAge in HCC (p.adjusted = 0.08-0.1) (Fig. 5C). At the same time, several other modules, such as those associated with inflammatory response, and amino acid and xenobiotic metabolism, contributed to the acceleration of biological age in tumor samples (Fig. 5B,D). Interestingly, most module-specific clocks displayed elevated tAge in hepatocellular carcinoma (Fig. 5C,E; Extended Data Fig. 18A), indicating that although tumors exhibit some anti-aging features, they are accompanied by accelerated aging of other cellular components and, therefore, don’t fully resemble the young organismal state. Examples of such established pro-aging hallmarks of cancer include inflammation and dysregulation of energy metabolism and mitochondrial function^109^. In agreement with that, functional enrichment revealed multiple shared hallmarks between signatures of HCC and signatures of mortality, such as a statistically significant upregulation of genes associated with inflammatory response, interferon signaling, p53 pathway and apoptosis, and downregulation of genes involved in oxidative phosphorylation and mitochondrial translation (Extended Data Fig. 18B; Supplementary Table 5B).

To examine if different age-related diseases share similar molecular signatures of aging and mortality, we estimated pairwise correlations between gene expression changes contributing to increased tAge in each of the models. Remarkably, the majority of examined pathologies exhibited overall similar aging- and mortality-associated signatures (Fig. 5F; Extended Data Fig. 17C). The top module contributing to the acceleration of biological age for most diseases was the inflammatory component (Fig. 5D), in agreement with the crucial role of chronic inflammation in progression of age-related pathologies^80,113,114^. Module-specific clocks reflecting various branches of immune response were also among those predicting the highest difference in tAge between healthy and diseased animals across the models (Fig. 5E; Extended Data Fig. 18A), followed by clocks representing mRNA splicing, chromatin modification/transcription, and energy metabolism (respiration/mitochondrial translation, oxidative phosphorylation, etc.). Accordingly, genes related to inflammatory response, interferon signaling, and apoptosis were upregulated in response to the majority of aging-related diseases, while genes involved in oxidative phosphorylation and respiratory electron transport were downregulated, resembling changes associated with mortality and aging (Extended Data Fig. 18B; Supplementary Table 5B). The only 2 models that didn’t display an upregulation of inflammatory response and positive contribution of inflammatory module to biological age were AD in the pineal gland and ischemic stroke in the heart tissue, which explains clustering of their mortality-associated profiles separately from most other pathologies (Fig. 5F; Extended Data Fig. 17C). Instead, pro-aging mechanisms in these models were mainly driven by the EMT/ECM organization module (Fig. 5D). Interestingly, this module had a negative effect on mortality tAge for many examined diseases, which may reflect ECM remodeling due to tissue fibrosis that often accompanies chronic inflammation^115,116^.

Since various pathologies were characterized by overall similar mortality-associated molecular profiles, we looked at top genes with the highest average contribution to acceleration of biological age across the models. Remarkably, some genes, including *Gpnmb*, *Cdkn1a*, and *Lgals3,* were significantly upregulated (p.adjusted < 0.05) in at least 5 models and had a strong pro-mortality effect shared across diseases (Fig. 5G). One of these factors, CDKN1A (p21), is a marker of intracellular damage and senescence^78,79^. On the other hand, GPNMB and galectin-3, being members of the inflammatory transcriptomic module, are involved in the modulation of immune response^72,117^ and are elevated in patients with certain diseases, including AD^73^ and arrhythmogenic cardiomyopathy^118^, while downregulation of galectin 3 in mouse CKD model is further associated with amelioration of renal inflammation and fibrosis^119^. Our study suggests that these genes are not only consistent multi-tissue biomarkers of aging and mortality but are also robust signatures of age-related diseases. To test if these factors also reflect health deterioration in humans, we utilized plasma proteomic data collected for more than 50,000 patients from UK Biobank^120^ and tested if concentration of GPNMB, CDKN1A, and LGALS3 in plasma is associated with various health outcomes after adjustment for chronological age and sex. Remarkably, all these proteins demonstrated a statistically significant positive association with human’s all-cause mortality and incidence of numerous diseases, including cardiac arrest, heart failure, liver cirrhosis, type I and type II diabetes, kidney failure, anaemia, atherosclerosis, emphysema, chronic ischemic heart disease, depression, and others (Fig. 5H; Supplementary Table 6). Interestingly, their concentration was also associated with established risk factors, such as obesity, hypertension, and sleep disorders. This suggests that the identified genes indeed reflect fundamental mechanisms of mammalian aging and mortality, shared across tissues, species, models of health degeneration, and levels of molecular organization.

### Molecular mechanisms of rejuvenation induced by heterochronic parabiosis

To identify molecular mechanisms associated with organismal rejuvenation, we utilized gene expression data corresponding to the models of heterochronic parabiosis^121^ and embryogenesis^122^. Heterochronic parabiosis (HPB), which involves connecting the circulatory systems of aged mice with younger counterparts, was shown to reverse multiple age-related impairments in old animals, including reduction of regenerative and repair capacity^123,124^, cardiac hypertrophy^125^, and cognitive function decline^126^. Recently, we observed a significant reduction of biological age measured with epigenetic clocks in the blood and liver of 20-month-old mice attached to 3-month-old counterparts for 3 months^121^. Interestingly, this effect was preserved 2 months after detachment from the young animal and resulted in lifespan extension of old mice compared to isochronic parabionts. At the same time, young mice attached to the old counterparts exhibited a transient increase of epigenetic age compared to the isochronic group, but this effect was diminished after 2 months of recovery^127^.

We analyzed liver gene expression data that was collected from young and old mice subjected to heterochronic and isochronic parabiosis for 3 months, sacrificed at the end of the intervention (Attached) or 2 months after detachment (Detached). In line with epigenetic data^121^, multi-tissue transcriptomic clocks of lifespan-adjusted age and mortality revealed a significant decrease of biological age induced by HPB in attached old mice (p.adjusted < 0.031), while the chronological clock showed marginal significance (p.adjusted = 0.075) (Fig. 6A; Extended Data Fig. 19A-B). Remarkably, the effect of HPB was even more prominent 2 months after detachment, being statistically significant for all the clocks. We also detected an elevated tAge in young heterochronic parabionts compared to isochronic controls right after the procedure (Fig. 6A; Extended Data Fig. 19A-B), whereas this difference was diminished after 2 months of recovery, in total agreement with the predictions of epigenetic clocks^127^.

**Fig. 6.**
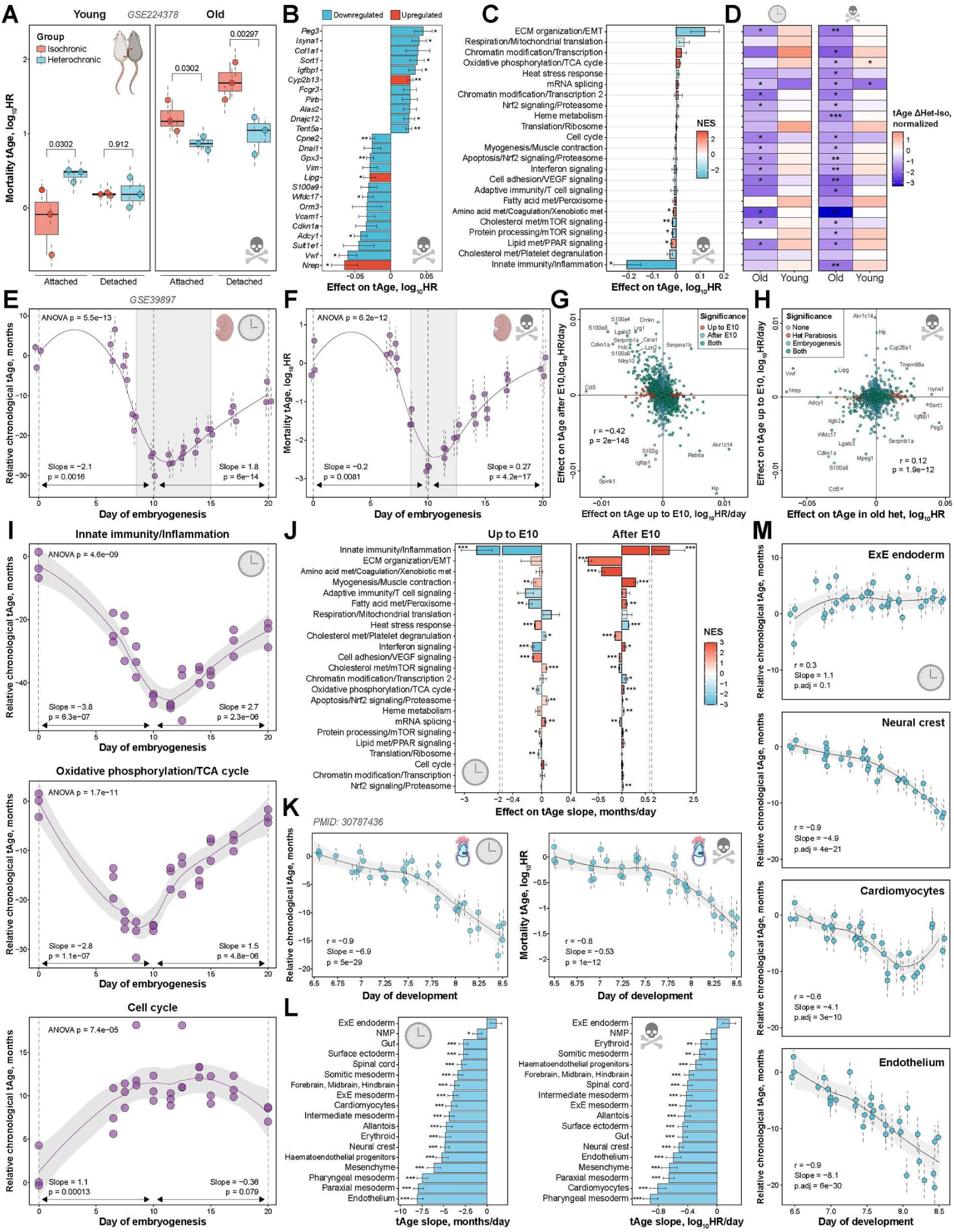
Heterochronic parabiosis and early embryogenesis exhibit rejuvenation at the level of gene expression. **A.** Mortality transcriptomic age (tAge) of livers from 3-month-old (left) and 20-month-old (right) mice subjected to isochronic or heterochronic parabiosis for 3 months, as assessed with the rodent multi-tissue Bayesian Ridge (BR) mortality clock. tAges between the groups were compared with a mixed effect model, and corresponding BH-adjusted p-values are shown in text. Data are tAges ± SE. **B.** Top genes driving pro- or anti-mortality transcriptomic changes in livers of old mice subjected to heterochronic parabiosis compared to the isochronic parabiosis group, according to the rodent multi-tissue EN mortality clock. Top 25 genes with the highest absolute effect on tAge difference (logFC clock coefficient) are shown. Attached and detached samples were pooled together for the logFC calculation, and attachment status was included in the model as a separate covariate. Genes up- and downregulated in old heterochronic parabiosis group are colored in red and blue, respectively. Statistical significance of logFC for each gene is indicated with asterisks. Data are mean tAge difference ± SE. **C.** Contributions of transcriptomic modules to mortality tAge difference in old mice subjected to heterochronic parabiosis, estimated with the rodent multi-tissue Elastic Net (EN) mortality clock. Individual modules are shown in rows and named after representative functions. Statistical significance for each module was assessed with ANOVA and indicated with asterisks. Bars are colored based on normalized enrichment scores (NES) from gene set enrichment analysis (GSEA), reflecting if genes associated with a particular module are generally up- (red) or downregulated (blue) in old mice subjected to heterochronic parabiosis compared to age-matched isochronic parabiosis group. Modules significantly enriched for up- or downregulated genes (BH-adjusted p-value < 0.05) are visualized with thick bars. Data are means ± SE. **D.** Normalized tAge difference between isochronic and heterochronic parabiosis groups assessed with all module-specific multi-tissue chronological (left) and mortality (right) clocks. Color and asterisks reflect size and statistical significance (BH-adjusted p-values) of tAge difference between isochronic and heterochronic parabiosis groups, assessed with the ANOVA. Increased and decreased tAges in heterochronic models are shown in red and blue, respectively. Module names are provided in (C). Het: Heterochronic; Iso: Isochronic. **E-F.** Chronological (E) and mortality (F) tAge of mouse embryos during development, assessed with mouse multi-tissue BR chronological and mortality clocks. The overall change of tAge was assessed with mixed effect ANOVA, whereas slopes of tAge change up to day 10 and after day 10 were assessed with a mixed effect linear model. The corresponding p-values and slope estimates are shown in the text. Loess regression curve is shown with a purple line. Dotted line shows time point with the minimum average tAge, and 95% confidence interval that contains putative minimum of embryo’s biological age (ground zero state) is shown with shaded grey area. Data are tAges ± SE. **G.** Association between weighted gene expression signatures of mortality (slope * clock coefficient) up to day 10 and after day 10 of mouse embryogenesis, according to the mouse multi-tissue EN mortality clock. The union of top 2,000 differentially expressed genes (with the lowest p-values) are shown on the plot. Statistical significance (BH-adjusted p-value < 0.05) is indicated with color. Pearson correlation coefficient and corresponding p-value are shown in text. **H.** Association between weighted gene expression signatures of mortality during early embryogenesis (slope * clock coefficient) and in old mice subjected to heterochronic parabiosis (logFC * clock coefficient). The union of top 2,000 differentially expressed genes (with the lowest p-values) are shown on the plot. **I.** Chronological tAge of mouse embryos during development estimated with representative module-specific multi-tissue chronological clocks. Slope of change up to day 10 and after day 10 was assessed with the linear regression. Corresponding p-values and slope estimates are shown in text. Purple line and shaded grey area around it reflect loess regression curve and its 95% confidence interval, respectively. Results for all module-specific chronological and mortality clocks are in Extended Data Fig. 21E. **J.** Contributions of transcriptomic modules to chronological tAge change in mouse embryos up to day 10 (left) and after day 10 (right) of development estimated with the multi-tissue Elastic Net (EN) chronological clock. Statistical significance for each module was assessed with the linear regression and indicated with asterisks. Bars are colored based on normalized enrichment scores (NES) from gene set enrichment analysis (GSEA), reflecting if genes associated with a particular module are generally up- (red) or downregulated (blue) during the corresponding stage of embryogenesis. Modules significantly enriched for up- or downregulated genes (BH-adjusted p-value < 0.05) are visualized with thick bars. Data are slopes ± SE. **K.** Chronological (left) and mortality (right) tAge of metacells representing mouse embryos between days 6.5 and 8.5 of embryogenesis, assessed with the mouse multi-tissue BR chronological clock. Slope of tAge change during this period was assessed with a mixed effect linear model. Pearson correlation coefficient, slope estimate and corresponding p-value are shown with text. Black line and shaded grey area around it reflect loess regression curve and its standard error, respectively. Data are tAges ± SE. **L.** Slopes of chronological (left) and mortality (right) tAges between days 6.5 and 8.5 for different cell lineages presented at the end of this interval. Statistical significance of the slope coefficient for each lineage indicated by asterisks was assessed with a mixed effect linear model. Data are slope estimates ± SE. **M.** Chronological tAge of metacells for representative cell lineages between days 6.5 and 8.5 of embryogenesis, assessed with the mouse multi-tissue BR chronological clock. Slope of tAge change was assessed with a mixed effect linear model. Pearson correlation coefficient, slope estimate and corresponding p-value are shown with text. Black line and shaded grey area around it reflect loess regression curve and its 95% confidence interval, respectively. Data are tAges ± SE. HR: Hazard Ratio; ECM: Extracellular matrix; EMT: Epithelial-Mesenchymal Transition; met: metabolism; ExE: Extraembryonic; NMP: Neuromesodermal Progenitors. ^ p.adj < 0.1; * p.adj < 0.05; ** p.adj < 0.01; *** p.adj < 0.001.

The top gene driving the rejuvenating effect of HPB in old mice across attached and detached groups was *Nrep*, upregulated in response to young blood exposure (Fig. 6B; Extended Data Fig. 19C), in line with the improved regenerative potential induced by parabiosis^123,124^. Besides, the anti-mortality molecular profile of old heterochronic parabionts was associated with downregulation of *Cdkn1a*, *Vwf* (Von Willebrand Factor), involved in regulation of platelet aggregation^128^, and *Vcam1* (vascular cell adhesion molecule 1), a modulator of vascular-immune cell interaction, whose ablation was shown to counteract neuroinflammation and improve learning and memory in age mice^129^.

Top modules contributing to the HPB-related decrease of biological age included those representing inflammation, cholesterol metabolism and platelet degranulation, lipid metabolism, and mTOR signaling (Fig. 6C; Extended Data Fig. 19D). Accordingly, a functional enrichment analysis revealed a statistically significant downregulation of genes involved in inflammatory response, complement cascade, p53 pathway, apoptosis, and mTOR signaling in old animals subjected to HPB, opposing signatures of mortality (Extended Data Fig. 19E; Supplementary Table 5C). Interestingly, most module-specific clocks predicted a statistically significant reduction of biological age in old HPB animals, especially at the level of mortality (Fig. 6D; Extended Data Fig. 19F), suggesting that heterochronic parabiosis induces systemic rejuvenation across different cellular components, ranging from immune response and apoptosis to mTOR signaling and lipid metabolism, in agreement with functional enrichment results. At the same, the effect of HPB in young animals was much less prominent and included a mixture of pro- and anti-mortality features, such as upregulation of complement cascade and downregulation of p53 pathway, respectively (Extended Data Fig. 19E).

### Molecular mechanisms of rejuvenation during early embryogenesis

To examine dynamics of transcriptomic age during mammalian embryonic development, we analyzed the gene expression dataset covering the whole period of mouse embryogenesis, from fertilized egg to newborn animal^122^. Previous studies demonstrated a rejuvenation event during the first 4.5-10.5 days of development at the level of DNA methylation, followed by a monotonous increase of epigenetic age^130^. This finding was consistent with the ground zero hypothesis, stating that early embryogenesis is accompanied by systemic repair and elimination of aging-associated damage accumulated in parental germ cells through their lifespan^131^. Interestingly, we also observed a similar U-shaped trajectory of biological age during embryogenesis at the level of gene expression, supported by multi-tissue transcriptomic clocks of chronological age, lifespan-adjusted age, and mortality (Fig. 6E-F; Extended Data Fig. 20A-B). Minimum tAge was detected on day 10 of embryonic development (E10), with the 95% confidence interval spanning between E8.5 and E12.5-E15. Therefore, according to our analysis, early development indeed appears to be associated with systemic rejuvenation of the organism at different levels of molecular organization.

Expression changes driving alteration of biological age up to E10 and after E10 were significantly correlated (Pearson’s r = −0.42) (Fig. 6G), suggesting that overall the same individual genes were responsible for rejuvenation during early embryogenesis and the increase of tAge afterwards. Top drivers of such dynamics included *Cdkn1a* and multiple genes associated with the regulation of inflammation, such as *S100a8*, *S100a9*, and *Lgals3*, all of which were downregulated up to E10 and upregulated afterwards (Fig. 6G; Extended Data Fig. 20C-D). Downregulation of the intracellular damage marker *Cdkn1a* during early embryonic development may reflect the global process of damage removal, in line with the ground zero model^131^. At the same time, mortality-associated signatures of early embryogenesis exhibited a weak positive statistically significant correlation with the signatures of heterochronic parabiosis (Pearson’s r = 0.12) (Fig. 6H), indicating that although generally these models share common molecular hallmarks of rejuvenation (such as downregulated *Cdkn1a* and *Lgals3*), their effects are also partially achieved through distinct biological mechanisms (e.g., *Nrep* is upregulated in old heterochronic parabionts but not during early embryonic development).

Overall, the first half of mouse embryogenesis was characterized by relative decrease of mRNA levels for most genes across the genome, while more genes were upregulated after E10 (Extended Data Fig. 20E). To test if the observed U-shaped trajectory of biological age may be driven by this global remodeling of the transcriptome, we examined components of tAge dynamics separately for genes up- and downregulated during each phase of embryonic development (Extended Data Fig. 20F-G). For both subsets of genes, we observed a statistically significant decrease of biological age up to E10 estimated with chronological and mortality clocks (p < 0.002), while tAge was increased during the later stages of embryogenesis according to both up- and downregulated gene sets (p < 6.4.10^-9^). At the same time, genes following the global expression pattern showed a higher amplitude of the U-shaped curve for biological age during embryonic development (Extended Data Fig. 20F-G), suggesting that it may indeed play a significant role in the observed rejuvenation signal.

Interestingly, genes downregulated up to E10 and upregulated afterwards mostly consisted of those associated with immune response and lipid metabolism, while genes with the opposite behavior were enriched for cell cycle and mRNA splicing (Extended Data Fig. 21A-B). In line with these findings, functional enrichment revealed strong downregulation of inflammatory response, interferon signaling, p53 pathway, and fatty acid metabolism up to E10, and their upregulation afterwards (Extended Data Fig. 21C; Supplementary Table 5D). Inflammation and interferon signaling modules were among top drivers of the rejuvenation signal during early embryogenesis according to both module-specific clocks (Fig. 6I; Extended Data Fig. 21E, 22) and the module contribution analysis (Fig. 6J; Extended Data Fig. 21D). Remarkably, most module-specific clocks, including those representing respiration/mitochondrial translation, lipid metabolism, and heat stress response, produced the U-shaped trajectory of tAge during embryogenesis (Extended Data Fig. 21E, 22), pointing to the systemic nature of the rejuvenation event during early embryonic development. At the same time, some modules, such as that associated with cell cycle, didn’t follow the common trajectory and demonstrated a significant elevation of tAge during the first phase of embryogenesis (Fig. 6I; Extended Data Fig. 21E), suggesting that development contains a mixture of pro- and anti-aging signals that may be distinguished at the level of co-regulated modules.

To examine if molecular rejuvenation during early embryogenesis is conserved across different cell lineages, we utilized scRNA-seq data collected from embryos between E6.5 and E8.5^132^. To validate a general anti-aging trend during this period of embryonic development, we pooled all cells within each sample into a single metacell and applied multi-tissue clocks of chronological age and mortality. Indeed, both clocks revealed a statistically significant decrease of tAge between E6.5 and E8.5 (Fig. 6K), consistent with the bulk data. To reconstruct ancestral lineage trajectories for each cell type presented at the last timepoint (E8.5), we then employed an optimal transport method (see Methods)^133^. Based on gene expression profiles of single cells, for each point of time it provided probability that a given cell is a part of lineage leading to the group of cells representing a particular cell type at E8.5. Overall, recovered trajectories resembled established lineages of embryonic tissues (Extended Data Fig. 23-24)^132,134,135^. Thus, extraembryonic endoderm was predicted to have an independent developmental trajectory, cardiomyocytes had mesodermal origin, endothelium was derived from haematoendothelial progenitors, while neural crest and spinal cord shared similar ancestral trajectories originating from neuroectoderm (Extended Data Fig. 23-24).

Utilizing reconstructed ancestral lineages, for every cell type presented at E8.5, we recovered the corresponding gene expression trajectory throughout the whole examined period of development. To do that, for each sample we aggregated gene expression profiles of individual cells into metacells, weighing them based on probabilities that a given cell is an ancestor of the respective cell type. Remarkably, multi-tissue transcriptomic clocks of chronological age and mortality revealed a statistically significant decrease of biological age from E6.5 to E8.5 for almost all examined cell lineages (Fig. 6L-M; Extended Data Fig. 25). The strongest rejuvenation effect was observed for mesodermal tissues, including endothelium, paraxial mesoderm and pharyngeal mesoderm. However, consistent effects were also observed for the representatives of other germ layers, including gut (endoderm), neural crest, and spinal cord (ectoderm).

The only cell type that showed a marginally significant increase of tAge during the examined period was extraembryonic endoderm (p.adjusted = 0.06-0.1), indicating that systemic rejuvenation observed during early embryogenesis may be restricted to cell layers with intraembryonic origin. This finding is in line with previous studies, showing that extraembryonic endoderm and ectoderm have a higher epigenetic age compared to other embryonic cell types from the same developmental stage^47^. Interestingly, although other cell lineages were characterized by decreased tAge at E8.5 compared to E6.5, some of them (e.g., cardiomyocytes) exhibited a U-shaped curve, demonstrating an increase of biological age starting from ∼E8 (Fig. 6M; Extended Data Fig. 25). This may indicate that although early development is accompanied with systemic rejuvenation, ground zero stage may vary across embryonic tissues and molecular modalities. Further studies focused on a detailed characterization of biological age dynamics during later stages of embryogenesis may shed light on mechanistic underpinnings of this rejuvenation process.

### Biological age deviation and rejuvenation in human models

To examine if the rodent multi-tissue transcriptomic clock model can be further expanded to cover aging-associated molecular changes in humans, we aggregated publicly available gene expression data from 12 sources, encompassing 2,296 brain, skeletal muscle, skin, and blood samples from healthy people of different ages as well as 10 brain samples from people with Hutchinson-Gilford progeria syndrome (Supplementary Table 1C). For every sample, we calculated chronological age divided by human’s maximum recorded lifespan (122 years^68,69^), as well as lifespan-adjusted age and expected all-cause mortality rate estimated from published survival data. We then trained unified relative multi-tissue clocks of chronological age, lifespan-adjusted age, and mortality using transcriptomic profiles from all 6,845 samples representing mice, rats, and humans. The developed EN and BR models accurately predicted differences in outcome variables in all 3 species (Fig. 7A; Extended Data Fig. 26A-B). Median Pearson’s r on test sets across 10 iterations was 0.94 to 0.95 for all the clocks, and median MAE for chronological clock was 5.5% to 5.7% of maximum lifespan, corresponding to ≈6.7-7 years and 2.6-2.8 months for humans and mice, respectively (Extended Data Fig. 26C-D).

**Fig. 7.**
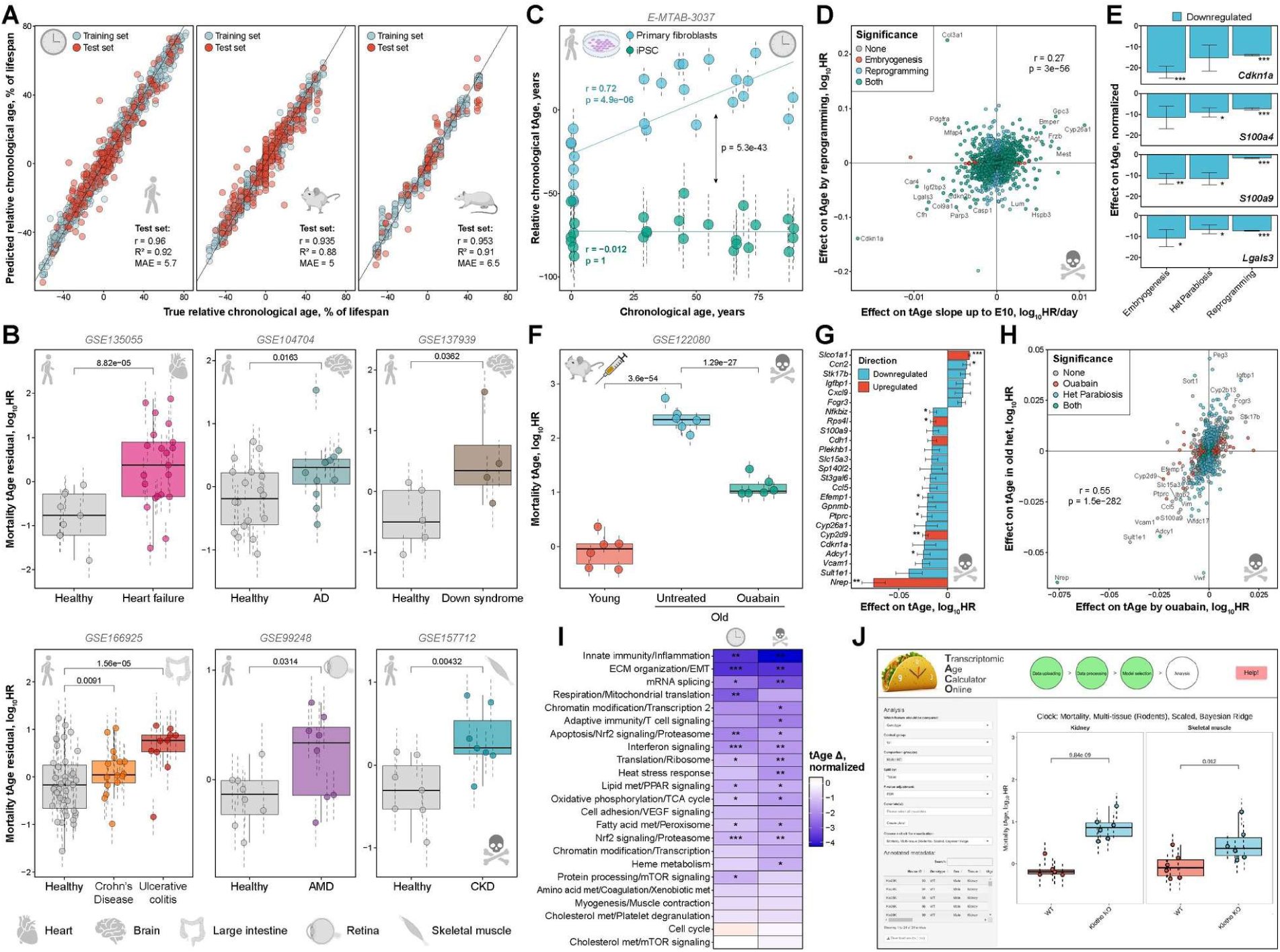
Conserved transcriptomic biomarkers of aging and mortality across mammals allow for the identification and characterization of health-regulating interventions. **A.** Accuracy of prediction of relative chronological age adjusted for species maximum lifespan with multi-tissue transcriptomic clocks trained with Elastic Net (EN) model on human, mouse and rat data (with scaling normalization). Accuracy of the trained model was assessed separately for humans (left), mice (middle) and rats (right). Training and test sets are denoted by color. Pearson correlation coefficient, R^2^ and mean absolute error (MAE) for test sets are shown in text. **B.** Mortality transcriptomic age (tAge) residual of tissues from healthy people (grey) and patients diagnosed with age-related and genetic diseases (indicated with color). tAges were calculated with the multi-species multi-tissue Bayesian Ridge (BR) mortality clock and adjusted for chronological age and sex. tAges between the groups were compared with mixed effect model, and corresponding BH-adjusted p-values are shown in text. Organs are depicted with icons. GEO IDs of corresponding datasets are provided in text. Data are tAges ± SE. AD: Alzheimer’s Disease; AMD: Age-Related Macular Degeneration; CKD: Chronic Kidney Disease. **C.** Chronological tAge of primary fibroblasts and reprogrammed induced pluripotent stem cells (iPSCs) from patients of different chronological ages, assessed with the multi-species multi-tissue BR chronological clock. Association between chronological age and tAge for each cell type was assessed with a mixed effect model, and corresponding p-value and Pearson correlation coefficient are shown with colored text. tAges of primary fibroblasts and corresponding iPSCs were compared with a mixed effect model, where patient ID was included as a factor covariate. The corresponding p-value is provided in black. Data are tAges ± SE. **D.** Association between weighted gene expression signatures of mortality up to day 10 of embryogenesis (slope * clock coefficient) and during iPSC reprogramming (logFC * clock coefficient), according to the multi-species multi-tissue EN mortality clock. Union of top 2,000 differentially expressed genes (with the lowest p-values) are shown on the plot. Statistical significance (BH-adjusted p-value < 0.05) is indicated with color. Pearson correlation coefficient and corresponding p-value are shown in text. **E.** Top genes contributing to decreased transcriptomic age during reprogramming, early embryogenesis and heterochronic parabiosis in old mice, according to the multi-species multi-tissue EN mortality clock. Statistical significance of expression change for each gene in the corresponding model is denoted with asterisks. Data are normalized tAge difference ± SE. **F.** Mortality transcriptomic age (tAge) of livers from 7-week-old (Young) and 24-month-old (Old) mice treated with saline (Untreated) or ouabain (Ouabain), examined with the rodent multi-tissue BR mortality clock. tAges between the groups were compared with a mixed effect model, and corresponding BH-adjusted p-values are shown in text. Data are tAges ± SE. **G.** Top genes driving pro- or anti-mortality transcriptomic changes in livers of old mice subjected to ouabain, according to the rodent multi-tissue EN mortality clock. Top 25 genes with the highest absolute effect on tAge difference (logFC * clock coefficient) are shown. Statistical significance of logFC for each gene is indicated with asterisks. Data are mean tAge difference ± SE. **H.** Association between weighted gene expression signatures of mortality (logFC * clock coefficient) of old mice subjected to ouabain and heterochronic parabiosis, according to the rodent multi-tissue EN mortality clock. Union of top 2,000 differentially expressed genes (with the lowest p-values) are shown on the plot. Statistical significance (BH-adjusted p-value < 0.05) is indicated with color. Pearson correlation coefficient and corresponding p-value are shown in text. **I.** Normalized tAge difference between control old mice and animals treated with oubain assessed with all module-specific multi-tissue chronological (left) and mortality (right) clocks. Color and asterisks reflect size and statistical significance (BH-adjusted p-values) of tAge difference between control and treated samples, assessed with the two-sample unpaired t-test. **J.** Screenshot of a TACO app that provides a platform and interactive interface to examine chronological, lifespan-adjusted and mortality tAges for the given gene expression data. HR: Hazard Ratio; ECM: Extracellular matrix; EMT: Epithelial-Mesenchymal Transition; met: metabolism. * p.adj < 0.05; ** p.adj < 0.01; *** p.adj < 0.001.

To test if the developed multi-species models capture biological age acceleration in humans caused by health impairment, we applied them to gene expression data collected from tissues of healthy people and patients diagnosed with various diseases, including heart failure (heart)^136^, Alzheimer’s disease (brain)^137^, Down syndrome (brain)^138^, Crohn’s disease (large intestine)^139^, ulcerative colitis (large intestine)^139^, age-related macular degeneration (AMD) (retina)^140^, and chronic kidney disease (CKD) (skeletal muscle)^141^. In agreement with health status, multi-species multi-tissue transcriptomic clocks of mortality predicted a statistically significant acceleration of biological age for patients diagnosed with all examined diseases compared to healthy people after adjustment for chronological age and sex (Fig. 7B), while chronological clocks predicted a tAge increase in all models except for Down syndrome where marginal significance was observed (p = 0.08) (Extended Data Fig. 27A). Therefore, transcriptomic hallmarks of aging and mortality appear to be generally conserved across species, in line with our previous findings^28^, allowing to develop universal models that are capable of distinguishing people and rodents with aging-associated diseases or genetically inherited impairments, such as *Klotho* knockout progeria (in mice) and Down syndrome (in humans).

To explore molecular hallmarks of rejuvenation in human cells and establish shared mechanisms of biological age reversal across species, we applied multi-species transcriptomic clocks to the model of cell reprogramming. For that, we utilized gene expression profiles of primary skin fibroblasts derived from patients of different ages along with induced pluripotent stem cells (iPSCs) reprogrammed from these cells through overexpression of Yamanaka factors (OSKM)^142^. In line with previous studies^39,143,144^, we observed a substantial decrease of biological age induced by reprogramming according to both chronological and mortality clocks (p < 2.2.10^-24^) (Fig. 7C; Extended Data Fig. 27D). Remarkably, while chronological age of patients was correlated with tAge of their skin fibroblasts, prefrontal cortex, and neurons transdifferentiated from the fibroblasts, there was no significant association between the age of patients and biological age of iPSCs (Pearson’s r = −0.012 and −0.017 for chronological and mortality clocks, respectively) (Fig. 7C; Extended Data Fig. 27B-D). This indicates that while aging-associated features of original cells may be weakly preserved in iPSCs at the level of DNA methylation^145^, aging- and mortality-associated hallmarks of gene expression are mostly reset during reprogramming regardless of the donor’s age.

Interestingly, mortality-associated gene expression changes contributing to alteration of biological age were positively correlated across all examined models of rejuvenation, including heterochronic parabiosis in old mice, early mouse embryogenesis (up to E10), and OSKM reprogramming of human fibroblasts (Extended Data Fig. 27E). At the same time, signatures of iPSCs exhibited higher correlation with the signatures of early embryonic development (Pearson’s r = 0.27, p = 3.10^−56^) (Fig. 7D), highlighting general similarity between iPSCs and embryonic stem cells^146^. Remarkably, some of the top hallmarks contributing to the decreased biological age were shared across all three models, including downregulation of *Cdkn1a* and several pro-inflammatory factors, such as *S100a4*, *S100a9*, and *Lgals3* (Fig. 7E). Therefore, *Cdkn1a* and *Lgals3* appear to be universal biomarkers of mammalian aging, rejuvenation, and mortality affected by multiple diseases and lifespan-regulating interventions, pointing to the presence of fundamental molecular hallmarks of aging and longevity conserved across experimental models and species.

### Identification and characterization of novel mortality-regulating interventions

The developed multi-tissue transcriptomic models allow one to screen for novel interventions with potential positive or negative effects on mortality-associated phenotypes. To identify candidate interventions that decrease biological age and improve healthspan, we utilized ClockBase^147^, a database with the collection of harmonized metadata and preprocessed publicly available datasets from Gene Expression Omnibus (GEO). We used 2,487 RNA-seq datasets corresponding to mouse tissues exposed to various pharmacological, dietary, and genetic interventions, and applied multi-tissue transcriptomic clocks of mortality to calculate tAge of every sample within this collection. Using ANOVA, we selected datasets with significant differences in biological age across the groups after adjustment for age and other factors, such as tissue and sex (p.adjusted < 0.05). Then, for every intervention within chosen datasets, we assessed the difference in tAge compared to age-matched control mice. This resulted in a number of interventions with a statistically significant effect on biological age in mouse tissues (p.adjusted < 0.05) (Extended Data Fig. 28A).

Overall, 3.6 times more interventions in mouse tissues induced a statistically significant increase of biological age than a decrease (Pearson’s chi square test p = 3.10^−73^), suggesting that it is generally easier to elevate tissue damage than to effectively reduce it. As expected, pro-mortality gene expression changes were observed in murine models of chronic diseases, including Alzheimer’s disease, myocardial infarction and spinal muscular atrophy; in short-lived animal models not used to train the clock (e.g., in rodents with conditional knockout of *Tsc2*^148^ and PLN-R14del mutation^149^); and in mice subjected to lipopolysaccharide (LPS) injection, kainic acid (brain seizure inducing compound) and unhealthy diets, such as methionine-choline deficient diet (MCD) and choline-deficient, L-amino acid-defined, high-fat diet (CDAHFD) associated with development of nonalcoholic steatohepatitis^150^. At the same time, several interventions resulted in the decrease of biological age in murine tissues. In addition to the established health-improving interventions, such as rapamycin and exercise, they included estrogen and progesterone applied to ovariectomized mice^151^ and the cardiac glycoside ouabain applied to aged healthy mice^152^ (Extended Data Fig. 28A).

Ovariectomy is an established pro-aging intervention that was shown to induce neuroinflammation and shorten lifespan of mouse females^153,154^, while sex hormones estrogen and progesterone were able to ameliorate some of the aging phenotypes in ovariectomized animals, improving their cognitive performance and working memory^155,156^. When applied to gene expression profiles of mammary glands of 3-month-old ovariectomized mice treated with estradiol and progesterone for 2 weeks^151^, multi-tissue transcriptomic clocks of chronological age and mortality revealed a strong statistically significant reduction of biological age induced by the treatment in luminal mature and luminal progenitor cells (p.adjusted < 10^-6^) (Extended Data Fig. 28D-E). Interestingly, mammary stem cells were not affected by this treatment, pointing to its cell-specific activity. Moreover, progesterone receptor antagonists telapristone and mifepristone partially mitigated the treatment effect, particularly in mammary progenitor cells. This finding confirms that the observed reduction in biological age was in part caused directly by the examined hormones.

Another identified intervention that produced the anti-aging effect on liver gene expression profile of healthy old mice was ouabain. Ouabain is a cardiac glycoside that was shown to exhibit senolytic activity and improve physical function of aged rodents^152^. According to both chronological and mortality multi-tissue transcriptomic clocks, 3-month treatment with ouabain was able to significantly decrease biological age of livers of 24-month-old female mice (p.adjusted < 4.10^−18^) (Fig. 7F; Extended Data Fig. 28B). Remarkably, this compound decreased predicted chronological tAge of mouse livers by ∼10 months, suggesting that it did not just slow down the accumulation of aging-associated changes but also activated mechanisms of molecular rejuvenation. Indeed, top drivers of the decrease in tAge induced by ouabain included many biomarkers of mortality shared by other models of rejuvenation, such as upregulation of *Nrep*, and downregulation of *Cdkn1a*, *Vcam1*, *Gpnmb*, and *Ccl5* (Fig. 7G). Overall, the mortality-associated gene expression signature of ouabain demonstrated a strong positive correlation with the rejuvenation signature of heterochronic parabiosis (Pearson’s r = 0.55) (Fig. 7H), indicating that this drug may be the first discovered pharmacological mimetic of HPB. As in the parabiosis model, the top module contributing to ouabain-induced molecular rejuvenation was the one representing innate immune response and inflammation (Extended Data Fig. 28C). At the same time, most module-specific clocks demonstrated a statistically significant decrease of chronological and mortality tAge induced by this compound, suggesting that ouabain may produce systemic rejuvenation of multiple cellular components, including immune response, respiration, ECM organization, mRNA splicing, and others (Fig. 7I). In agreement with our observations, experimental data demonstrated the reversal of multiple age-related phenotypes produced by ouabain, including the reduction of senescence and local inflammation in the liver revealed by immunohistochemistry, restoration of albumin and phosphate levels in blood, and improvement of motor coordination and strength^152^. Therefore, ouabain appears to be a promising candidate for future studies that may reveal its effect on other organs and molecular modalities and shed light on specific mechanisms of tissue rejuvenation induced by this compound.

Finally, to facilitate application of the developed transcriptomic clocks, we developed an interactive platform TACO (Transcriptomic Age Calculator Online). This tool simplifies implementation of the models by offering software and an interface for RNA-seq data preprocessing and application of liver and multi-tissue clocks of chronological age, lifespan-adjusted age, and expected mortality, developed using Elastic Net and Bayesian Ridge models. Additionally, TACO enables users to visualize clock results and conduct analyses to assess statistical significance of tAge difference between groups (Fig. 7J).

## DISCUSSION

While aging is accompanied by the systemic deterioration of organismal functions, and models that predict chronological age (i.e., chronological clocks) may be used to indirectly assess biological age, a unified transcriptomic model of mortality that encompasses both aging and various models of lifespan-shortening and longevity interventions (i.e., mortality clocks) has been lacking. The ability to capture the effects of these interventions is the key to targeting the aging process in order to extend lifespan/healthspan and to partially rejuvenate organismal systems, as opposed to merely assessing progression through aging. The robust biologically interpretable multi-species multi-tissue mortality biomarkers we report in the study provided critical insights into mechanisms of longevity, chronic disease, and rejuvenation.

To develop these tools, we sequenced the transcriptomes of a large cohort of ITP mice subjected to various neutral and longevity interventions, expanded the dataset with publicly available gene expression data representing organs of mice and rats across various strains and lifespan-regulating interventions, connected these models with survival data, and performed a meta-analysis of aggregated 4,539 rodent samples, which allowed us to identify multi-tissue transcriptomic signatures of aging, mortality rate, and maximum lifespan. Interestingly, they included well established biomarkers of longevity, such as *Igf1*^58^ and *Fmo3*^63^, but also other genes, such as *Lgals3*, a modulator of inflammation and fibrosis^117,119^, *Ddost*, involved in processing of AGEs^59^, *Nmrk1*, a member of NMN biosynthesis pathway^64^, and *Nrep*, involved in wound healing and scar formation^82^. The signatures of aging were negatively correlated with maximum lifespan and positively correlated with age-adjusted mortality, indicating that molecular hallmarks of aging are generally associated with impaired health and shortened lifespan. At the functional level, aging and mortality were characterized by upregulation of genes involved in inflammation, complement cascade, apoptosis, and p53 pathway, while oxidative phosphorylation, fatty acid metabolism, and mitochondrial translation were negatively associated with mortality, both before and after adjustment for age, in agreement with the established role of these functions in the regulation of aging and longevity^28,33,157–161^.

Utilizing the aggregated dataset, we developed rodent multi-tissue transcriptomic clocks of chronological age, lifespan-adjusted age, and mortality. The clocks showed high quality on test sets (Pearson’s r = 0.94 to 0.96) and were able to capture age dynamics in the tissues and datasets that were not used in training. While the chronological clock could distinguish the effect of detrimental genetic and dietary models, it did not show a decrease in biological age in response to longevity interventions. In contrast, clocks of lifespan-adjusted age and mortality both captured aging-associated dynamics and correctly predicted the effect of lifespan-shortening and extending interventions, indicating that they can be applied for the identification and characterization of diverse models of aging and longevity. By expanding our dataset with gene expression profiles of 2,306 human samples collected from brain, skin, skeletal muscle and blood, we also developed unified multi-species multi-tissue transcriptomic clocks, capable of predicting chronological age, lifespan-adjusted age, and mortality risk across mice, rats and humans, with Pearson’s r on test sets > 0.92 for each of the species. The multi-species clocks were validated on 7 models of human age-related and genetic diseases, including heart failure, Alzheimer’s disease, Down syndrome, Crohn’s disease, ulcerative colitis, AMD, and CKD, where they were able to capture the acceleration of biological age in tissues of diseased patients after adjustment for chronological age and sex. Therefore, mammals share common aging- and mortality-associated molecular mechanisms that may be combined in a single transcriptomic model. While mouse- and human-specific transcriptomic clocks of chronological age have been developed previously^42,44,45,48^, multi-species multi-tissue clocks trained to predict mortality across various models of healthspan regulation, including aging, genetic manipulations, diets, and pharmacological treatments, provide a direct assessment of universal mortality-associated biomarkers that reflect an aggregated level of molecular damage and are validated both in short-lived and long-lived models.

To characterize specific cellular components associated with age and mortality, we performed a network analysis. We identified 26 co-regulated transcriptomic modules enriched for specific cellular pathways and used them to develop interpretable multi-tissue module-specific clocks of chronological age and mortality. Remarkably, clocks trained on 23 modules were able to predict outcomes with Pearson’s r > 0.3, and the corresponding mortality clocks demonstrated both a positive association with chronological age and negative correlation with lifespan, indicating that the identified individual gene expression modules contain information about aging and health status of the tissue. To demonstrate application of the developed tools, we sequenced kidneys and skeletal muscles from *Klotho* KO mice and age-matched controls. Chronological and mortality transcriptomic clocks revealed an acceleration of biological age induced by this genetic manipulation, with more prominent effect in kidneys where *Klotho* is typically expressed. Among top contributors to this pro-mortality effect, we identified upregulation of cellular damage marker *Cdkn1a* along with downregulation of genes associated with mitochondrial function and ECM organization, while inflammation was not elevated in *Klotho* KO mice and didn’t induce biological age acceleration. These findings were in line with the results of functional enrichment, suggesting that modules provide interpretable information about the cellular functional components driving observed differences in biological age.

By applying transcriptomic clocks to the single-cell Tabula Muris data, we found that most cell types, including stem cells, exhibit an increase in biological age and share common pro-aging signatures, such as upregulation of pro-inflammatory chemokine *Ccl5*^89^ and downregulation of modulators of circadian rhythm *Dbp*^91^ and ECM organization *Sparc*^94^. To examine if *Klotho* KO induces systemic acceleration of biological age across different cell types, we performed snRNA-seq of kidneys and brains derived from an 8-week-old *Klotho* KO mouse and an age-matched control mouse. We observed a significant increase of tAge in most kidney and brain cell types, including proximal tubule epithelial cells, endothelial cells, astrocytes, and neurons. Interestingly, microglia demonstrated a marginally significant decrease of biological age, in agreement with downregulation of inflammatory module, observed across sequenced tissues. Thus, different cell types share similar molecular hallmarks of aging and mortality, and the same intervention can have opposite effects on different cell types and intracellular components revealing complex systemic nature of aging and regulation of longevity.

This was further highlighted by rodent models of aging-related diseases. While clocks captured accelerated biological age in most models, including ischemic stroke, Alzheimer’s disease, chronic kidney disease, diabetic neuropathy, and nonalcoholic steatohepatitis, the model of hepatocellular carcinoma showed a decreased transcriptomic age, driven by EMT/ECM organization and cell cycle modules. Remarkably, module-specific clocks revealed a pro-aging signal for multiple cellular components in HCC samples, while the clock trained on genes associated with EMT/ECM organization demonstrated marginally significant rejuvenation induced by HCC, in line with the dedifferentiation and remodeling of ECM associated with tumor progression^109,112^. Various age-related diseases shared common pro-aging and pro-mortality signatures, such as upregulation of the inflammatory module and several specific genes contributing to tAge increase, including *Gpnmb*, *Lgals3* and *Cdkn1a*. Interestingly, higher levels of the corresponding proteins in human plasma were associated with the increased mortality rate after adjustment for chronological age and sex, along with higher incidence of numerous aging-related diseases, including heart and kidney failure, type II diabetes, atherosclerosis, and others, and established risk factors, such as obesity and hypertension. Therefore, the identified factors represent universal biomarkers of biological age acceleration, shared across diseases, tissues, species, and levels of molecular organization.

Several established models of organismal rejuvenation, including heterochronic parabiosis in old mice, early embryogenesis, and cellular reprogramming, were also validated by the developed transcriptomic clocks that captured a decrease of biological age induced by these models. Interestingly, anti-mortality signatures of all 3 models were positively correlated with each other, with the most similar effect observed between the signatures of iPSCs and early embryonic development. Common hallmarks of rejuvenation models included downregulation of *Cdkn1a* and *Lgals3*, again highlighting the universal role of these genes in the regulation of damage affected by both aging-accelerating and rejuvenating models. Module-specific clocks revealed a systemic effect of early embryogenesis and HPB on biological age shared by multiple cellular components, with the strongest contribution of the inflammatory module. Remarkably, rejuvenation of the embryo during early organogenesis (between days E6.5 and E8.5) was also observed across almost all cell lineages, with the exception of extraembryonic endoderm, in line with the results of epigenetic clocks predicting higher biological age for extraembryonic tissues^47^. At the same time, several individual module-specific clocks, including those associated with cell cycle, didn’t show a decrease in biological age during early phases of embryogenesis, indicating that global rejuvenation may be still accompanied by pro-aging signals of individual cellular components.

Finally, by employing Clockbase database^147^, we searched for interventions with a significant anti-mortality effect according to multi-tissue transcriptomic clocks. We identified several such cases, including the therapy of ovariectomized mice with the combination of estrogen and progesterone^151^ as well as treatment of old healthy mice with the cardiac glycoside ouabain^152^. In particular, a 3-month treatment with ouabain was able to decrease the biological age of 24-month-old mice by ∼10 months suggesting a first example of a radical *in vivo* rejuvenation effect induced by a single-molecule pharmacological intervention. Remarkably, the effect of ouabain was systemic, supported by most module-specific clocks, biochemical assays, and physiological tests, while its anti-mortality signature was strongly correlated with that of HPB, suggesting that ouabain mimics many effects of heterochronic parabiosis. Indeed, ouabain induced multiple rejuvenation hallmarks shared by HPB, such as downregulation of *Cdkn1a* and *Vcam1*, a vascular cell adhesion molecule shown to modulate neuroinflammation and impair the cognitive function of old mice^129^. Therefore, transcriptomic biomarkers developed in this study provide an opportunity to identify interventions promoting or counteracting molecular mechanisms of mortality, and characterize specific targets associated with their effects at the level of cell types, intracellular functional components, and individual genes. These methods may facilitate the development of effective therapies capable of ameliorating age-related impairments and restoring a young functional phenotype in mammalian organisms.

Our study has several limitations. Molecular signatures and transcriptomic clocks developed in this study may reflect both causal and compensatory response to damage accumulation associated with aging and mortality. Methods of causal inference, such as Mendelian Randomization, may be utilized to partially distinguish between causal and adaptive molecular changes in humans highlighted by the clocks and signatures^41^. Human UK Biobank data used for the analysis of the association between concentration of GPNMB, CDKN1A, and LGALS3 and health outcomes is based on an observational study. Therefore, some of the identified associations with diseases and risk factors may be driven by co-morbidities or other confounders, such as socioeconomic status, smoking and drinking behavior. Finally, an association of the identified co-regulated transcriptional modules with corresponding cellular processes was performed based on the pre-existing annotation of genes and may be further improved and directly tested using functional assays and established perturbations in cell culture.

Overall, our study resolves the challenge of predicting biological age and mortality by developing a comprehensive transcriptomic model encompassing both aging and lifespan-modulating interventions. By sequencing the transcriptomes of a large cohort of mice subjected to various treatments and integrating the dataset with gene expression profiles from multiple strains, species, and intervention models with corresponding survival data, we identified multi-species multi-tissue signatures of lifespan and mortality. We further developed robust clocks that accurately predict chronological age, lifespan-adjusted age, and mortality, which in turn revealed common molecular mechanisms of aging across cell types, and deconstructed them into functional components, providing broad insights into the effects of genetic, dietary, and pharmacological interventions on biological age as well as critical insights into the mechanisms of diseases and established models of rejuvenation. Furthermore, our findings revealed universal biomarkers of mammalian mortality contributing to the observed effect on biological age induced by most chronic diseases and anti-aging models, and identified potential rejuvenation interventions, such as the cardiac glycoside ouabain. Our study underscores the complexity of aging and mortality mechanisms, the interplay between various processes involved, and the clear potential for developing therapies to extend healthspan and lifespan.

## ACKNOWLEDGMENTS

We would like to thank Dr. Richard Miller for spearheading the ITP part of this study, providing ITP samples and critical insights into the study, and Iaroslava Pavlova for valuable suggestions and assistance with data visualization. The study was supported by NIA grants to VNG, DEH, and RS, by the Japan Agency for Medical Research and Development (AMED) 24zf0127001h0004 to TA, and by the James Fickel and Michael Antonov Foundations. RS was supported by a Senior Research Career Scientist Award from the Department of Veterans Affairs Office of Research and Development. snRNA-seq was supported by JSPS KAKENHI Grant-in-Aid for Early-Career Scientists (JP22K15354), Takeda Science Foundation, The Uehara Memorial Foundation, The Naito Foundation, and Okinaka Memorial Institute for Medical Research.

## AUTHOR CONTRIBUTIONS

A.T. and V.N.G. conceived and designed this research; A.T., D.K., K.Y., M.D., A.M., L.M.K., A.V., Y.T., T.K., and U.K. performed research and data analysis; Y.T., B.Z., A.M., and T.K. carried out RNA sequencing; A.T., K.Y., T.K., H.L., M.M., J.M.V.R., T.A., S.E.D., and V.N.G contributed new reagents/sample/analytic tools; A.T. and V.N.G. supervised the study; A.T. and V.N.G. wrote the manuscript with contributions from all other authors.

## COMPETING INTERESTS

A.T. and V.N.G. are the inventors on a U.S. Patent Application related to this work.

## METHODS

### Animals from ITP cohorts

Liver samples of mice from the Interventions Testing Program (ITP) were acquired from the collections of University of Michigan Medical School (UM), University of Texas (UT) and The Jackson Laboratory (TJL) obtained from animals of 2015, 2016 and 2017 cohorts^16,21–23^. 22-23-month-old female and male mice were sacrificed following subjection to 20 interventions, including 17-dimethylaminoethylamino-17-demethoxygeldanamycin hydrochloride (17-DMAG) (30 ppm, as in ^16^), b-guanidinopropionic acid (bGPA) (3300 ppm, as in ^16^), minocycline (300 ppm, as in ^16^), mitoQ (100 ppm, as in ^16^), rapamycin applied from 20 months (42 ppm, as in ^16^), canagliflozin (180 ppm, as in ^21^), candesartan cilexetil (30 ppm, as in ^22^), geranylgeranyl acetone (600 ppm, as in ^22^), 17-α-estradiol applied to males from 20 months (14.4 ppm, as in ^22^), 17-α-estradiol applied to males from 16 months (14.4 ppm, as in ^22^), 3-(3-hydroxybenzyl)-5-methylbenzo[d]oxazol-2(3H)-one (MIF098) (240 ppm, as in ^22^), nicotinamide riboside (NR) (1000 ppm, as in ^22^), 1,3-butanediol (100,000 ppm, as in ^23^), captopril (180 ppm, as in ^23^), leucine (40,000 ppm, as in ^23^), PB125 (100 ppm, as in ^23^), sulindac (5 ppm, as in ^23^), syringaresinol (300 ppm, as in ^23^), combination of rapamycin and acarbose applied from 9 months (14.7 ppm and 1000 ppm, as in ^23^), and combination of rapamycin and acarbose applied from 16 months (14.7 ppm and 1000 ppm, as in ^23^). Besides, livers were taken from control untreated male and female mice sacrificed at 4-6 and 22 months of age. All mice were fed *ad libitum* with the same diet (Purina 5LG6) made in the same commercial diet kitchen (TestDiet, Richmond, IN, USA). Mice represented genetically heterogenous UM-HET3 strain, produced by crossing female CByB6F1/J and male C3D2F1/J animals. Therefore, each rodent in the cohort had a unique genetic background but shared the same set of inbred grandparents (C57BL/6J, BALB/cByJ, C3H/HeJ, and DBA/2J). All interventions continued until the organisms were sacrificed. Details of the mouse housing and methods used for health monitoring are provided in ^10^.

### *Klotho* knockout (KO) mice

8-week-old male WT mice (C57BL6/J) and *α-klotho*^-/-^ knockout mice were purchased from CLEA Japan. Mice were maintained under specific pathogen-free conditions, on a 12-h light–dark cycle and fed normal diet. The animal protocols were approved by the Tohoku University Institutional Animal Care and Use Committee. The animal procedures performed conform to the NIH guidelines (Guide for the Care and Use of Laboratory Animals).

### Bulk RNA-seq profiling of mouse tissues

For RNA-seq profiling of liver samples from ITP cohorts, 3 biological replicates per sex were utilized for each intervention, except for canagliflozin represented by 7 biological replicates per sex. In addition, samples from 24 old and 6 young control mice per sex across different cohorts were sequenced, resulting in the total of 182 samples (Supplementary Table 1A). For RNA-seq analysis of bulk kidney and gastrocnemius skeletal muscle samples from WT and *Klotho* KO male mice, 6 biological replicates per group were utilized for every tissue, resulting in the total of 24 samples. RNA was extracted with PureLink RNA Mini Kit as described in the protocol and passed to sequencing. Paired-end sequencing with 150 bp read length was performed on Illumina NovaSeq 6000 platform.

### Nuclei extraction from tissues of *Klotho* KO mice

Brains and kidneys were harvested following cervical dislocation and flash frozen in liquid nitrogen. The cortical region of brain and radially section of kidney were dissected. Their nuclei were isolated with the Minute™ single nucleus isolation kit for neuronal tissue/cells and the Minute™ single nucleus isolation kit for tissue/cells (Cat# BN-020, SN-047, Invent Biotechnologies, INC), following manufacturer’s protocol with minor modifications. Specifically, the isolated nuclei were resuspended in 5% BSA solution with RNase inhibitor (Cat# 1000494; 10x genomics).

### Single-nucleus RNA-seq profiling of *Klotho* knockout mouse tissues

SnRNA-seq was performed with Chromium Single Cell 5’ Reagent Kits (Cat# PN-1000265; 10x genomics) according to the manufacturer’s instructions. Briefly, single-cell suspensions from the tissues were loaded into a Chromium Next GEM Chip K Single Cell Kit (Cat# PN-1000287; 10x Genomics), and a Chromium Controller was used for barcoding and cDNA synthesis. A Chromium Next GEM Single Cell 5’ Library & Gel Bead Kit v2 (Cat# PN-1000265; 10x Genomics) was used to amplify the cDNA and construct the cDNA libraries. RNA sequencing was performed by DNAFORM (Yokohama, Kanagawa, Japan). The sequenced raw data were then processed using Cell Ranger v7.0.0 and the R package Seurat (v5.0.1). For quality control, nuclei that were detected with a mitochondrial content > 10%, < 200 genes, or > 4,000 genes were considered dying cells, empty droplets, and doublets, respectively, and were excluded. Cells were annotated using established sets of marker genes for brain and kidney cell types^162–164^.

### Estimates of maximum lifespan, expected mortality and lifespan-adjusted age

Estimates of maximum registered lifespan for mice, rats and humans were obtained from AnAge database (4, 3.8 and 122 years, respectively)^68,69^. Survival curves corresponding to control mice and rats from various strains and sexes, and to mice subjected to different lifespan-shortening, neutral or longevity interventions were obtained from published resources and digitalized, where necessary^2,6–8,10,12,14,16–21,23–25,53–57,67,165–204^. Survival data for overall human population and patients with Hutchinson-Gilford progeria syndrome (HGPS) were downloaded from US CDC report^205^ and from the corresponding clinical trial, respectively^4^. Survival data was fitted with Gompertz mortality rate function adjusted for left truncation with flexsurvreg function from flexsurv R package^206^:

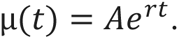

Mean and standard error estimates of parameters *ln*(*A*) and *r* from individual mortality curves corresponding to the same experimental model (strain, sex, intervention) were used to calculate aggregated meta-parameters. For that, mixed-effect model with restricted maximum likelihood (REML) criterion was applied via rma.uni function from metafor R package^207^.

For every sample presented in the transcriptomic dataset, logarithm of its expected hazard rate *log*_10_(µ) was calculated based on chronological age and annotation (strain, sex, intervention) of the sample as an average of *log*_10_(µ) from the corresponding aggregated model and individual model, fitted from survival data that was derived from the same study as the gene expression data. For transcriptomic data downloaded from studies without survival data, estimates of respective aggregated mortality functions were utilized. Quality of Gompertz model fitting was assessed with correlation coefficient between 90^th^ percentile lifespan estimates from real survival data and from survival data simulated with corresponding fitted Gompertz models. Maximum lifespan estimates (99.9^th^ percentile) for every experimental group were derived from survival data simulated with the corresponding fitted Gompertz functions. Lifespan-adjusted age was calculated as a chronological age divided by the expected maximum lifespan (99.9^th^ percentile) estimated for the corresponding experimental model.

### Preprocessing and analysis of ITP cohort RNA-seq data

Reads were mapped to mouse genome (GRCm39) with STAR (version 2.7.11b)^208^ and counted via featureCounts (version 2.0.6)^209^. To filter out non-expressed genes, we left only genes with at least 10 reads in at least 20% of samples, resulting in 13,422 detected genes according to Entrez annotation. Filtered data was then passed to RLE normalization^210^. Samples were considered outliers based on Principal Component Analysis (PCA) performed separately for each sex, if their 1^st^ or 2^nd^ principal component (PC) values were more than 1.5 interquartile ranges (IQR) above the third quartile or below the first quartile. Following this rule, 21 samples were identified as outliers and filtered out from the data. All interventions were still present in the dataset after the filtering step.

Transcriptomic signatures of chronological age, lifespan-adjusted age, expected hazard rate (*log*_10_(µ)) and expected maximum lifespan (90^th^ percentile), estimated directly from survival data (“estimate”) or corresponding Gompertz mortality models (“model”), were identified with linear models fitted with edgeR package^211^ separately for each sex and together for both sexes. For the pooled signature, sex was included in the model as a separate covariate. Besides, associations with age-adjusted expected mortality rate (in log scale), lifespan-adjusted age and maximum lifespan were examined by including chronological age as a separate covariate in corresponding models. Genes were considered statistically significant if their p-value, adjusted with Benjamini-Hochberg (BH) method^212^, was lower than 0.05.

### Aggregation and preprocessing of transcriptomic data across sources

Gene expression data for meta-analysis were obtained from the current study (ITP cohorts), published studies^201,213,214^ and the repositories NCBI GEO^215^, ArrayExpress^216^, and SRA^217^ with the following identifiers: GSE3150^218^, GSE6591^219^, GSE11291^220^, GSE34378^175^, GSE27625^221^, GSE53960^222^, GSE66715^223^, GSE132040^224^, GSE146977^225^, GSE145480^226^, GSE3129^218^, GSE55272^183^, GSE131754^33^, GSE81959^8^, GSE36838^173^, GSE46895^177^, GSE51108^178^, GSE26267^67^, GSE92486^227^, GSE124772^193^, GSE101657^187^, GSE117188^191^, GSE32609^2^, GSE1093^228^, GSE40977^229^, GSE48333^179^, GSE48331^179^, GSE49000^182^, GSE97074^188^, GSE108379^189^, GSE122116^230^, GSE121395^231^, GSE122080^152^, GSE122085^230^, GSE122243^232^, GSE122367^233^, GSE126814^234^, GSE127758^129^, GSE128724^233^, GSE129083^235^, GSE137504^236^, GSE139204^237^, GSE140286^238^, GSE143304^239^, GSE145972^240^, GSE146796^241^, GSE147666^242^, GSE148647^243^, GSE149569^244^, GSE155407^245^, GSE156762^246^, GSE158980^247^, GSE165409^248^, GSE154832^249^, GSE166615^250^, GSE166778^251^, GSE168211^252^, GSE168610^253^, GSE175571^254^, GSE178770^255^, GSE69952^256^, GSE78130^257^, GSE83931^258^, GSE93833, GSE124483^160^, GSE127475^194^, GSE190939^200^, GSE201207^259^, GSE141252^260^, GSE134781^204^, GSE134780^204^, GSE219203^57^, GSE149029^197^, GSE39699^176^, GSE75574^261^, GSE89272^262^, GSE54853^181^, GSE155064^263^, GSE86882^53^, GSE234563^202^, GSE51202^180^, GSE84390^264^, GSE123293^265^, GSE234667^266^, GSE52794^267^, GSE241904^203^, GSE75192^257^, PRJNA281127^268^, PRJNA516151^269^, E-MTAB-3374^270^, E-MEXP-2320^56^, and E-MEXP-153^271^. For training of multi-species transcriptomic clock, human gene expression data was derived from GTEx^272^ (blood samples) and the following GEO datasets: GSE103232^257^, GSE113957^45^, GSE134080^273^, GSE17612^274^, GSE21935^275^, GSE226189^276^, GSE36192^277^, GSE40645^278^, GSE5086^279^, GSE53890^280^, GSE674^281^, and GSE75337^257^.

Preprocessing pipelines for RNA-seq and microarray datasets were as described in ^28^. Briefly, for each RNA-seq dataset, raw counts were downloaded, and soft filtering was performed to remove unexpressed genes. Afterwards, genes were mapped to Entrez IDs, followed by RLE normalization, log-transformation and scaling of gene expression profiles. For microarray datasets, data was log-transformed, mapped to Entrez IDs and scaled. After data aggregation, we filtered out genes presented in less than 20% of datasets, keeping 18,592 genes that passed this criterion. As a quality control, for every sample in the meta-dataset we calculated correlation between its expression profile and median expression profiles across all samples corresponding to the same tissues. Samples with Spearman ρ < 0.5 were removed from the data as outliers. This resulted in 4,539 mouse and rat gene expression samples with known chronological age (Supplementary Table 1B), 4,487 of which also had associated estimates of lifespan-adjusted age, expected mortality and maximum lifespan. The same pipeline was used to preprocess 2,306 human tissue samples (Supplementary Table 1C), followed by mapping of human genes to mouse 1-to-1 (mutually exclusive) orthologs. Integration of human data resulted in a meta-dataset consisting of 6,845 tissue samples from mice, rats and humans, utilized for training the multi-species transcriptomic clock.

For clock development, we used either scaled gene expression profiles or utilized YuGene normalization^70^, shown to be effective for integration of transcriptomic data across multiple platforms^71^. To adjust for potential batch effects and differences in baseline expression across tissues, within every dataset and tissue we calculated relative gene expression of each sample using the following algorithm. (1) For each dataset and tissue, we randomly selected a subset of control samples of the same chronological age and sex; (2) for a selected reference group, we estimated median expression profile separately for each gene and (3) subtracted it from the expression profile of each sample in the examined dataset and tissue. Similarly, we subtracted chronological age, lifespan-adjusted age, expected mortality rate (in log scale) and maximum lifespan of the reference group from corresponding estimates of every sample. Therefore, for each sample we obtained changes in log-expression and changes in chronological age, lifespan-adjusted age, mortality rate (in log scale) and expected lifespan compared to a randomly chosen control group from the same dataset and tissue, thus correcting for batch effects and organ-specific differences in gene expression. These features and outcome variables were then employed to train the corresponding relative transcriptomic clocks.

For the identification of rodent gene expression signatures of aging, mortality and lifespan, and for a weighted gene co-expression network analysis (WGCNA), a similar algorithm of relative expression estimation was applied within every dataset, tissue and sex, to keep only transcriptomic changes associated with aging and the effect of various lifespan-regulating interventions. Outliers were identified during PCA analysis as samples with values of the 1^st^ or 2^nd^ PCs falling outside the boundaries of the quartiles ±1.5 IQR. After filtering out such samples, the dataset for meta-analysis included relative gene expression data for 4,476 mouse and rat samples. Besides, we filtered out genes detected in less than 50% of samples. This resulted in the expression data for 12,894 genes.

### Transcriptomic signatures of aging, mortality and lifespan on the aggregated meta-dataset

To identify transcriptomic signatures of chronological age, lifespan-adjusted age, expected mortality and maximum lifespan based on the aggregated meta-dataset of rodent relative gene expression, we utilized linear mixed-effect model with REML criterion via lmer function from lme4 and lmerTest R packages^282^. We considered change in log-expression as an outcome variable and difference in trait of interest as an independent variable, while tissue and source of data were implemented in the model as random terms, and sex was included as a fixed term. To examine genes associated with age-adjusted differences in expected mortality (in log scale), lifespan-adjusted age and maximum lifespan, we introduced difference in chronological age into a mixed-effect model as a separate fixed term covariate. Genes were considered statistically significant if their p-value, adjusted with Benjamini-Hochberg (BH) method^212^, was lower than 0.05. Pairwise overlaps between transcriptomic signatures associated with different traits were assessed separately for up- and downregulated genes with Fisher’s exact test, and together with Pearson’s chi-square test.

### Development of transcriptomic clocks

To develop rodent liver-specific, mouse multi-tissue, rodent multi-tissue and multi-species (based on mice, rats and humans) multi-tissue transcriptomic clocks of chronological age, lifespan-adjusted age and expected mortality, we utilized Elastic Net (EN) and Bayesian Ridge (BR) models from scikit-learn library in Python (functions ElasticNet and BayesianRidge, respectively). Models were applied to relative expression datasets normalized with scaling and YuGene methods. For the chronological clock, only control samples not subjected to interventions were utilized, while for the lifespan-adjusted and mortality clocks all samples with estimated outcome values from both control organisms and those subjected to interventions were employed. To account for differences in lifespan across species, chronological age was divided by maximum recorded lifespan (4, 3.8 and 122 years for mice, rats and humans), and the resulting value was provided as a response value to train the chronological clock.

90% and 10% of samples were selected as training and test sets, respectively, with stratification based on species and type of intervention (controls, lifespan-shortening, neutral and lifespan-extending). Genes detected in less than 20% of samples in training set were filtered out. Missing values for every gene were imputed with median expression calculated on training set. Hyperparameters of the model were selected based on R^2^ accuracy metric estimated with grid search (GridSearchCV) through 5-fold cross-validation method (KFold). For EN model, optimized hyperparameters included *alpha* (between 10^-5^ and 10^2^), *l1_ratio* (between 0 and 1) and those defining whether scaling of features should be performed (*with_mean* and *with_std* parameters of StandardScaler function). For BR model, trained hyperparameters included initial values for alpha *alpha_init* (between 10^-3^ and 10^-1^) and lambda *lambda_init* (between 10^-3^ and 1) as well as *with_mean* and *with_std* parameters of StandardScaler function. Interestingly, for all trained models *with_std* parameter was selected to be 0, suggesting that standardization of features doesn’t improve the quality of the model. Pearson correlation coefficient, R^2^ and mean absolute error (MAE) were examined as metrics of accuracy on test set.

For robust quality assessment, nested cross-validation with 10 iterations was performed. In particular, we randomly divided data into training and test sets 10 times, each time training the model with 5-fold cross-validation and applying it to the test set. Median accuracy (Pearson’s r, R^2^ and MAE) across 10 test sets was considered as a final quality of the model. To have a reference for MAE assessment, we also calculated MAE based on random guessing – i.e., using mean outcome value on the training set as a prediction for every sample in the test set. Mean and standard error of gene coefficients from trained EN models across 10 test sets were calculated to identify top predictors of rodent chronological age and mortality.

Similarly, for rodent multi-tissue data we trained transcriptomic clocks of absolute chronological age, lifespan-adjusted age and expected mortality with linear or log-log transformed outcomes, and clock of relative expected maximum lifespan with chronological age included or excluded from the set of features. Besides, we developed tissue-specific transcriptomic clocks of chronological age utilizing data only from liver, kidney, brain or skeletal muscle. For rodent multi-tissue chronological clock, in addition to EN and BR, we also tested other machine learning models, including support vector machines (SVM), random forest (RF), k-nearest neighbors (KNN) and light gradient-boosting machine (LightGBM) (functions SVR, RandomForestRegressor, KNeighborsRegressor and LGBMRegressor from lightgbm package, respectively). For all models, in addition to model hyperparameters, StandartScaler *with_mean* and *with_std* parameters were optimized. Besides, for SVM model optimized hyperparameters included *kernel* (linear or rbf), regularization parameter *C* (from 10^-4^ to 1) and *epsilon* (from 10^-3^ to 1); for KNN they included *n_neighbors* (from 2 to 10), *weights* (uniform or distance) and power parameter *p* (from 1 to 5); for RF they included *n_estimators* (from 100 to 2000), *max_depth* (from 10 to 110), *min_samples_split* (from 2 to 10) and *min_samples_leaf* (from 1 to 4); and for LightGBM they included *boosting_type* (dart or gbdt), *num_leaves* (from 100 to 250), *max_depth* (from 1 to 3), *n_estimators* (from 100 to 5000), *subsample* (from 0.5 to 1), *min_child_samples* (from 50 to 200), *min_child_weight* (from 10^-5^ to 0.1), *reg_alpha* (from 10^-7^ to 1), *reg_lambda* (from 10^-8^ to 1) and *colsample_bytree* (from 0.2 to 1). To identify an optimal set of hyperparameters for RF and LightGBM, we employed RandomizedSearchCV with 60 iterations.

To improve applicability of the clocks on new unseen data, we trained final models with GroupKFold cross-validation, ensuring that optimization of hyperparameters occurs on independent datasets not used to train the model. Gene coefficients of final EN models trained on the whole datasets are in Supplementary Table 3. To test the quality of the trained clocks on novel tissues and datasets, we utilized Leave-One-Tissue-Out (LOTO) and Leave-One-Dataset-Out (LODO) approaches. Iteratively, we trained models on all tissues or datasets but one and estimated the quality of the model on the remaining tissue or dataset. Besides, we tested the quality of clocks to separately predict chronological age and effect of interventions on maximum lifespan in new independent data. Correlation with chronological age was examined using tAge predictions for all control samples estimated during LODO test. The effect of a given intervention on maximum lifespan was calculated within each dataset and experimental group by dividing expected maximum lifespan (99.9^th^ percentile) of cohort subjected to the intervention by expected maximum lifespan of corresponding control cohort. Then we estimated the correlation of log-ratio of maximum lifespans 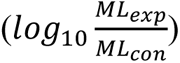 with average difference in tAge between experimental and control group after adjustment for chronological age and tissue estimated with ANOVA (*tAge*_exp_ − *tAge*_con_) across all available interventions and experimental groups (datasets, strains, sexes). In addition, we divided all interventions into 3 categories based on the size of their effect on maximum lifespan 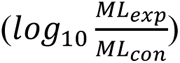: (1) lifespan-extending interventions with log-ratio of lifespans > IQR (more than +15.7% to the original maximum lifespan); (2) lifespan-shortening interventions with log-ratio of lifespans < -IQR (less than 86.4% of the original maximum lifespan); and (3) neutral interventions with the intermediate effect on lifespan. For lifespan-extending and lifespan-shortening interventions, we assessed statistical significance of non-zero mean tAge difference across interventions within given category using mixed-effect model with rma.mv function from metafor package (with REML criterion)^207^. Mean and standard error of (*tAge*_exp_ − *tAge*_con_) from ANOVA were considered as an outcome variable, while source of expression data was included as a random term. Pairwise comparison of predicted effect on tAge between categories of interventions was assessed using similar mixed-effect model, with intervention category included as a fixed term. Resulted p-values were adjusted for multiple comparisons with BH apptoach^212^.

### Co-regulated transcriptomic modules of aging and longevity

Preprocessed rodent relative expression data were filtered using the goodSamplesGenes function from the WGCNA package^83^ to remove samples and genes with a high number of missing values. The remaining missing values were then imputed with the median values for each gene. To identify co-regulated transcriptomic modules associated with aging and longevity, Pearson correlation metric was utilized to determine the similarity of expression profiles across all pairs of genes and construct adjacency matrix with the SoftThreshold = 4. Hierarchical clustering was performed based on the topological overlap matrix (TOM) with the settings TOMType = “unsigned”, deepSplit = 2, and minClusterSize = 20, using the dynamic tree cut method. The same procedure was applied separately to male and female data to ensure consistency of the results. Pairwise overlaps of modules identified across sexes were assessed with Fisher’s exact test (Extended Data Fig. 7B). Obtained 28 clusters of co-regulated genes were further iteratively filtered, ensuring that all genes within each module have Spearman ρ > 0.475 with the median gene expression profile estimated based on all genes included in the given module. The threshold of 0.475 reflects maximum Spearman ρ with median expression profile for 2,500 randomly chosen genes not attributed to any module according to WGCNA, multiplied by 1.1 to account for potential variation caused by random subsetting of this gene set.

Functional and upstream regulator enrichment of gene lists associated with each identified module was performed using Fisher’s exact test via the Python package GSEApy^283^. KEGG^284^, MSigDB HALLMARK^285^, REACTOME^286^ and ChEA transcription factor^287^ ontologies were utilized for the analysis (Supplementary table 4A). Adjustment for multiple comparisons was performed with Benjamini-Hochberg method, and terms with adjusted p-value < 0.05 were considered significantly enriched by the given module. Based on the results of functional enrichment, 26 modules were annotated based on their top representative enriched pathways (Supplementary table 4B). The resulting annotated and filtered 26 modules were utilized for subsequent analysis and development of module-specific clocks.

Gene correlation network was visualized via spectral embedding^288,289^. First, negative correlations were replaced by zeros to ensure non-negative edge weights, providing an adjacency matrix A(i,j)=max(cor(i,j), 0). Subsequently, the Laplacian D-A and the random walk normalized Laplacian *L*_*norm* = *D*^-1^*L* were computed, where D is a diagonal degree matrix, with entries equal to the weighted degrees of nodes. Eigenvalues and eigenvectors of the normalized Laplacian were then computed using eig function from numpy.linalg module, and the coefficients of the 1^st^ and the 3^rd^ non-trivial eigenvectors were used as vertical and horizontal coordinates of a node. To visualize WGCNA modules, a sample of 3000 genes was generated and displayed, including all genes associated with the corresponding module and randomly sampled background genes.

For each module, its eigengenes reflecting the 1^st^ PC were calculated for every sample in the rodent meta-dataset of relative expression. Partial Spearman correlation between module eigengenes after adjustment for chronological age was estimated. Besides, for eigengenes of every module, we calculated their Spearman correlation with chronological age and partial Spearman correlation with maximum lifespan adjusted for chronological age. Sparse partial correlation network of identified modules, chronological age and expected maximum lifespan was computed using graphical lasso regularization and model selection based on extended Bayesian information criterion^290^ with EBICglasso function from bootnet package^291^. The resulting Gaussian graphical model was visualized with qgraph R package using the spring layout algorithm. The same method was used to visualize partial correlation network of tAge predictions of module-specific clocks estimated on test sets.

Rodent multi-tissue module-specific EN and BR clocks of relative chronological age and expected mortality were trained similarly to the conventional clocks, with stratification and 5-fold cross-validation, using only genes associated with the given module. To obtain robust estimates of accuracy, we also performed nested cross-validation, dividing whole dataset into 10 equal folds, and iteratively training each module-specific clock on 9 subsets and testing its quality on the remaining fold. To examine the overall power of module genes to predict age and mortality, we also trained composite clocks using all genes associated with at least one module. We then assessed the accuracy of the module-specific clocks by calculating average Pearson’s r with response variable across 10 test sets. Besides, for mortality clocks we separately calculated correlation of their tAge predictions with chronological age on control test samples and partial correlation of their predictions with expected maximum lifespan after adjustment for chronological age.

### Application of Transcriptomic Clocks

Preprocessing of gene expression data for clock application was similar to that performed for aggregated meta-data. Particularly, for RNA-seq data non-expressed genes were removed based on soft filtering, genes were mapped to Entrez IDs, and the resulting data was normalized with RLE method, log-transformed and subjected to scaling or YuGene transformation. For preprocessed microarray data, expression matrix was log-transformed (if required), mapped to Entrez IDs and subjected to scaling or YuGene transformation. Afterwards, one of control groups was chosen as a reference group and used to calculate relative log-expression for each sample in the dataset. Relative gene expression data were then passed to transcriptomic clocks. Missing values were imputed with median values precalculated during the training of the clock. For convenience of tAge interpretation, the outputs of the chronological clocks were multiplied by the maximum registered lifespan of the corresponding species (4, 3.8, and 122 years for mice, rats and humans^68,69^), while the outputs of lifespan-adjusted clocks were multiplied by 100 to represent percentage of the passed expected lifespan. For EN conventional and module-specific clocks, comparison of predicted biological age across groups was assessed with t-test, ANOVA or linear model. The outcomes of the BR clocks were compared with a mixed-effect model, fitted with the point estimates of tAge and their corresponding standard errors provided by the BR model.

Top genes associated with the effect on tAge were estimated using the following procedure. First, differential expression between groups of interest was estimated with limma^292^. Then, logFC or slopes of log-expression together with their standard errors were multiplied by the corresponding gene coefficient from the utilized clock. The resulting clock-weighted log-expression changes were ranked and visualized on a barplot. Besides, they were employed to estimate correlation between aging- or mortality-associated signatures of different models.

The impact of various transcriptomic modules on the difference in tAge predicted by a conventional EN clock after adjustment for the effect of other modules was assessed as follows. First, standard preprocessing of the dataset was performed. Then, calculated relative expression values were multiplied by gene coefficients from the conventional EN clock trained only on genes associated with any module. Since EN transcriptomic clocks represent linear models without standardization of features, the composite relative tAge for each sample can be represented as a sum of tAge contributions from individual modules plus some fixed intercept equal for all samples in the data:

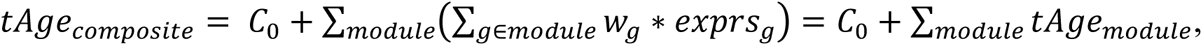

where *exprs*_g_ and *w*_g_ are relative log-expression and clock weight for a given gene *g*, while tAge_module_ reflects contribution of the corresponding module to the composite tAge value. Therefore, for every module we then calculated its impact on sample tAge by adding contributions of all genes associated with this module. Finally, t-test, ANOVA or linear model were employed to assess the average contribution of each module to tAge difference between control and experimental groups, together with its statistical significance. Resulting p-values were adjusted for multiple comparisons with BH procedure^212^. Besides, for each module we estimated if gene expression signature of the examined intervention is enriched for the list of genes associated with this module using gene set enrichment analysis (GSEA)^293^, as described below.

### Functional Enrichment Analysis of Signatures

For identification of functions enriched by transcriptomic signatures of examined traits (chronological age, lifespan-adjusted age, expected mortality and expected maximum lifespan), along with signatures of individual interventions (*Klotho* KO, age-related diseases, heterochronic parabiosis, embryogenesis), we performed gene set enrichment analysis (GSEA)^293^ on a pre-ranked list of genes based on log_10_(p-value) corrected by the sign of regulation, calculated as:

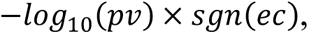

where *pv* and *ec* are p-value and expression change for certain gene, respectively, estimated with ANOVA, linear model or mixed-effect model, and *sgn* is signum function (is equal to 1, −1 and 0 if value is positive, negative and equal to 0, respectively). REACTOME, KEGG and HALLMARKS ontologies from Molecular Signature Database (MSigDB) have been used as gene sets for GSEA^293^. False discovery rate (FDR) cutoff of 0.05 was used to select statistically significant associations.

Hierarchical clustering of enriched functions for a heatmap was performed based on normalized enrichment scores (NES), using complete linkage and euclidean distance. Enriched functions that (i) were significantly enriched (adjusted p-value <0.05) by multiple signatures of aging, mortality and lifespan, and (ii) represented different aspects of cellular biology, were selected for visualization. The whole lists of statistically significant enriched functions are available in Supplementary Tables 2 and 5.

### Analysis of Tabula Muris Senis single-cell RNA-seq data

To validate the applicability of the clock to single-cell RNA-seq (scRNA-seq) data, we utilized droplet Tabula Muris Senis dataset^88^ that covers gene expression profiles of single cells from multiple organs extracted from mice of different ages, ranging from 1 to 30 months. For the subsequent analysis we filtered out organs that covered less than 15 months of mouse lifespan and were presented in less than 7 samples. Employing the data on remaining 9 tissues, we assessed if mouse chronological multi-tissue transcriptomic clock can predict dynamics of chronological age across organs based on expression profiles of individual cells. To do that, we utilized metacell procedure, randomly selecting a subset of cells corresponding to the same tissue and organism, and pooling reads of these cells together, combining their gene expression profiles in a single meta-profile (Fig. 4A). The resulting meta-profiles were treated as bulk samples, and the standard preprocessing and clock application pipelines were applied. First, we utilized this algorithm, combining all available cells for a given tissue and organism in a single metacell. We assessed Pearson’s correlation coefficient between chronological age and tAge of a metacell predicted with mouse chronological multi-tissue EN clock and obtained accuracy similar to that achieved on the bulk data (Extended Data Fig. 12A-B). We then repeated the same procedure, utilizing different number of randomly chosen cells for a metacell aggregation (from 1 to 750 cells per metacell). In each case, for every tissue and cell number threshold we obtained a set of metacells, each of which represented a single mouse. We then calculated Pearson’s correlation between chronological age and metacell tAge within every tissue for each cell number (Fig. 4B). Besides, we calculated average coverage across metacells for every cell number threshold to establish the dependence between total coverage of the sample and accuracy of the transcriptomic clock (Fig. 4C).

We then utilized metacell algorithm separately for each cell type, aggregating all cells annotated as the same cell type into a single metacell within each organism. For each cell type, association between chronological age and tAge predicted with the mouse chronological or mortality multi-tissue EN clock was assessed with linear model. Corresponding p-values were adjusted for multiple comparisons with BH procedure^212^. To compare aging-associated signatures across cell types, we selected genes expressed in at least 75% of cell types and estimated their weighted age-associated expression slopes within each cell type, calculated as *Slope* ∗ *Coefficient*, where *Slope* is a slope estimate representing association of log-expression with chronological age estimated with linear model from limma package^292^, and *Coefficient* is a coefficient of the corresponding gene in the mouse chronological multi-tissue EN clock. We then calculated pairwise Pearson correlation between obtained weighted signatures using a union of top 1,000 genes associated with age (with the lowest p-value from linear model) for each pair of cell types. The resulted correlation matrix was used to calculate a distance matrix and perform complete hierarchical clustering of cell types according to their aging-associated weighted signatures. To identify top shared pro-aging and pro-mortality signatures across cell types, we normalized weighted slopes by standard deviation estimated across all examined genes for the given cell type and ranked genes by average normalized weighted slopes. Top 5 genes driving pro-aging and pro-mortality phenotype according to mouse chronological and mortality multi-tissue EN clocks were visualized on barplots (Fig. 4F; Extended Data Fig. 13B).

### Preprocessing and analysis of *Klotho* KO bulk and single-nucleus RNA-seq data

Bulk RNA-seq reads were mapped to mouse genome (GRCm39) with STAR (version 2.7.11b)^208^ and counted via featureCounts (version 2.0.6)^209^. To filter out non-expressed genes, we left only genes with at least 10 reads in at least 25% of samples, resulting in 15,744 detected genes according to Entrez annotation. Filtered data was then passed to RLE normalization^210^ and preprocessed for the clock separately for each tissue. Control samples were selected as a reference group for calculation of relative expression. To have unbiased estimation of tAge for *Klotho* KO mice, we retrained transcriptomic clocks excluding *Klotho* gene from training set. tAges estimated with resulting rodent chronological, lifespan-adjusted and mortality multi-tissue BR clocks were compared between control and *Klotho* KO mice separately for skeletal muscle and kidney using mixed-effect model. Difference in *Klotho* expression between control and progeroid mice was assessed with edgeR^211^. To identify top genes driving the difference in tAge, we performed differential expression analysis between control and *Klotho* KO groups separately for each organ with limma^292^ and multiplied logFC by clock coefficients, as described above. Contribution of each module to the difference in tAge according to conventional clock was examined separately for each tissue as described above. Difference in tAge between control and progeroid mice predicted by module-specific clocks in each organ was assessed with ANOVA. P-values across modules were adjusted with BH approach. Pearson correlation between weighted mortality-associated signatures of *Klotho* KO in kidney and skeletal muscle was assessed as described above based on the union of top 2,000 differentially expressed genes (with the lowest p-value) in each organ.

snRNA-seq data from brain and kidney of control and *Klotho* KO mice were normalized, subjected to PCA and visualized with Uniform Manifold Approximation and Projection (UMAP) separately for each organ using Seurat package^294^ (Fig. 4I; Extended Data Fig. 14A-C). To establish homogenous clusters of cell types for kidney, we filtered out cells with dimensionally reduced coordinates falling outside the boundaries of the quartiles ±1.5 IQR calculated for the corresponding cell type (Fig. 4H). Following filtering, we aggregated individual cells within each cell type and experimental group into metacells using coverage threshold of 200K reads per metacell. Difference in expression of *Klotho* between metacells representing control and progeroid mice was assessed for each cell type with edgeR. Afterwards, standard preprocessing pipeline was applied separately for each cell type with control metacells selected as a reference group, followed by application of the clocks trained on all genes excluding *Klotho*. Predictions of tAges estimated with rodent chronological and mortality multi-tissue BR clocks were compared between control and progeroid mice with mixed-effect model separately for each cell type. P-values across cell types were adjusted for multiple comparison with BH procedure.

For most represented cell types, i.e., neurons and proximal tubule, we performed differential expression analysis, calculating logFC of genes between metacells of control and *Klotho* KO mice. Resulting logFC profiles were multiplied by clock coefficients to establish top genes driving aging- and mortality-associated phenotype in neurons and proximal tubules of progeroid animals. Contribution of modules into tAge difference between control and *Klotho* KO mice was examined separately for each cell type. Difference in tAge estimated with module-specific clocks in neurons and proximal tubules of control and *Klotho* KO mice was tested with ANOVA. P-values across modules were adjusted for multiple comparisons using BH approach. Pairwise Pearson correlation between weighted mortality-associated signatures of different tissues and cell types was assessed based on the union of top 2,000 genes differentially expressed (with the lowest p-value) in at least one cell type or tissue from the given pair.

### Mortality-associated mechanisms of aging-related diseases

Gene expression data representing tissues of mice and rats with chronic disease models and corresponding age-matched controls were obtained from the NCBI GEO repository with the identifiers GSE142633^101^, GSE137482^104^, GSE148350^106^, GSE129586^101^, GSE102558^105^, GSE189377^102^, GSE197699^103^, GSE120977^107^, GSE26538^108^. Each dataset was preprocessed separately as described above. Control samples from the youngest age group were selected as a reference group for calculation of relative expression. Following preprocessing, we applied rodent chronological and mortality multi-tissue transcriptomic clocks. Difference in tAge between healthy and diseased animals estimated with BR clocks was assessed with mixed-effect model separately for each age group and time point within the dataset. P-values in datasets with multiple comparisons were adjusted with BH approach.

To identify top genes driving the observed age acceleration, for every dataset we performed differential expression analysis between control and disease groups. For datasets with several age groups (GSE142633, GSE137482, GSE148350), age group was included in the limma model as a separate factorial covariate. For datasets with multiple time points (GSE102558, GSE148350), time after stroke induction was included in the limma model as a numeric covariate, and slope of its association with log-expression was evaluated. For dataset with expression profiles of two hemispheres examined after stroke induction (GSE148350), slope of association between log-expression and time after disease was evaluated separately for ipsilateral and contralateral hemispheres. The resulting differential expression profiles were multiplied by mortality EN clock coefficients, and top 25 genes with pro- or anti-mortality effects were visualized with barplots and volcano plots. Pearson correlation between weighted aging- and mortality-associated signatures of various disease models was estimated based on the union of top 1,000 differentially expressed genes (with the lowest p-value) for each pair of disease signatures. To identify top pro-mortality signatures shared across models, we normalized each weighted mortality signature by standard deviation across all genes, filtered out genes with missing values in at least one signature and ranked genes based on their average normalized mortality association. Using a similar methodology, we identified top modules contributing to the pro- or anti-mortality effect on tAge across examined disease models. Contribution of each module to tAge difference for every disease was assessed with the same statistical models utilized for identification of differentially expressed genes (ANOVA or linear regression depending on the dataset). Same models were also used to examine change in biological age induced by diseases according to individual module-specific clocks. P-values across modules were adjusted for multiple comparison using BH procedure.

Gene expression profiles of human organs representing healthy cohort and patients with diagnosed age-related and genetic diseases were uploaded from GEO using the following identifiers: GSE135055^136^, GSE104704^137^, GSE137939^138^, GSE166925^139^, GSE99248^140^ and GSE157712^141^. Following preprocessing of each dataset, we estimated biological ages of samples using multi-species chronological and mortality multi-tissue BR transcriptomic clocks. Difference in tAge between healthy and diseased patients was assessed with a mixed-effect model with chronological age and sex included as fixed term covariates. P-values in datasets with multiple comparisons were adjusted with BH approach.

### Association between plasma protein concentration and health outcomes in humans

To investigate the association between concentration of GPNMB, CDKN1A and LGALS3 in human plasma and the incidence of mortality and various diseases in the UK Biobank cohort^120^, we utilized Cox proportional hazards regression analysis using the coxph function from the survival package in R. The UK Biobank provides normalized protein expression data for selected patients, measured by Olink using Proximity Extension Assay (PEA). UK Biobank includes information on the first disease occurrence for patients. Covariates in our model included chronological age, sex, and the interaction between these two factors. To standardize protein levels, we divided concentrations by their standard deviation. Since the patient data is continuously updated, for every disease, including death, the right censoring date was the date of the latest recorded occurrence of that disease in the UK Biobank. The data was left truncated, meaning that patients with disease occurrences reported before the day of sample collection were excluded. Because the samples were collected at different time points, the time to disease was calculated as the time of disease occurrence minus the time of sample collection. For censored samples, the time to the latest recorded occurrence among patients in the dataset was used, subtracted by the time of sample collection. Several diseases were combined into groups, and the earliest occurrence within each group was used for the model. The following diseases were grouped by ICD-10 codes: dementia (F00-F03), kidney failure (N17-N19), hypertension (I10, I15), and myocardial infarction (I21, I22). To control for multiple testing, p-values were adjusted using BH procedure.

### Rejuvenation-associated mechanisms of heterochronic parabiosis

Liver gene expression profiles of 3-month-old and 20-month-old mice subjected to isochronic or heterochronic parabiosis (HPB) for 3 months following sacrifice or 2-month recovery period after detachment were downloaded from GSE224378^121^. Data was preprocessed as discussed above, and young isochronic attached mice were selected as a reference group for calculation of relative expression. tAge of samples was estimated with rodent chronological, lifespan-adjusted and mortality multi-tissue transcriptomic clocks. Difference in tAge between isochronic and heterochronic animals estimated with BR models was assessed with mixed-effect model separately for each age and attachment status group. P-values were adjusted for multiple comparisons using BH procedure.

To identify genes associated with pro- or anti-mortality effect of HPB in old mice, we identified differentially expressed genes between old mice subjected to isochronic and heterochronic parabiosis, including attachment status as a separate factorial covariate in limma model. logFC between isochronic and heterochronic parabionts were multiplied by rodent mortality multi-tissue EN clock coefficients. Genes were ranked based on absolute clock-weighted logFC. Module contributions to chronological and mortality tAge difference between old isochronic and heterochronic animals were assessed using ANOVA model with attachment status included as a separate factorial covariate. The same statistical model was utilized to examine differences in predictions of individual module-specific clocks of chronological age and mortality between isochronic and heterochronic mice separately for young and old animals. P-values across modules were adjusted using BH procedure.

### Transcriptomic age trajectory during mouse embryogenesis

Microarray gene expression data representing mouse embryos at different stages of development, from fertilized eggs to newborn mice, were downloaded from GSE39897^122^. Samples corresponding to day 0 of embryogenesis (fertilized eggs) were selected as a reference group used to calculate relative expression. tAges of embryo samples were estimated with mouse chronological and mortality multi-tissue EN and BR clocks. Overall variation of biological age during development was assessed with ANOVA. Decrease of tAge up to day 10 of embryogenesis (E10) and its increase afterwards was tested with linear model and mixed effect models for predictions of EN and BR models, respectively. To estimate 95% confidence interval of development stage that contains minimum tAge (ground zero state), we took samples representing stage with the minimum average tAge (E10) and iteratively expanded this subset with samples corresponding to earlier time points until statistically significant slope of biological age with time was detected (p-value < 0.05). Same procedure was utilized for the right side of the confidence interval (later stages of development). Identified interval was shown on tAge trajectory plot with a grey shadow rectangular (Fig. 6E-F; Extended Data Fig. 20A-B).

Top genes associated with changes in chronological and mortality tAge during early and late embryogenesis were identified with the linear regression limma model based on samples representing stages up to E10 and stages after E10, respectively. Gene log-expression slopes were multiplied by corresponding coefficients from mouse chronological and mortality multi-tissue EN clocks, and top genes with the average absolute weighted slope across phases of embryogenesis were visualized on barplots and volcano plots. Module contributions to tAge dynamics during early (up to E10) and late (after E10) embryogenesis were calculated based on coefficients of mouse chronological and mortality EN models and assessed with linear models. The same statistical model was utilized to examine the dynamics of tAges predicted with individual module-specific clocks of chronological age and mortality. P-values across modules were adjusted for multiple comparisons using BH procedure.

To assess if early and late embryogenesis are characterized by global remodeling of gene expression profiles, we examined differences in absolute slopes between statistically significant up- and downregulated genes (BH-adjusted p-value < 0.05) separately for each phase of embryonic development (up to E10 and after E10) using Wilcoxon rank sum test (Extended Data Fig. 20E). To assess dynamics of biological age separately for genes with increased and decreased expression, for each phase of embryogenesis we selected statistically significant up- or downregulated genes (BH-adjusted p-value < 0.05) and calculated tAge components formed by these genes for every sample using coefficients of mouse chronological or mortality multi-tissue EN clocks. We then examined if estimated tAge components change during early and late phases of embryonic development using linear regression model separately for up- and downregulated genes (Extended Data Fig. 20F-G). Functional enrichment of gene sets upregulated during early embryogenesis (up to E10) and downregulated afterwards or vice versa was performed with Fisher’s exact test based on GO BP ontology, utilizing genes expressed in this dataset as a background and using enrichGO function from clusterProfiler package in R^295^. Terms with BH-adjusted p-value < 0.05 were considered statistically significant. Redundant functional terms were removed with the simplify function. Network of enriched maintained functional terms was visualized with emapplot function from enrichplot package in R (Extended Data Fig. 21A-B).

Pearson correlation of mortality-associated weighted slopes of log-expression during early and late embryonic development was estimated based on the union of the top 2,000 differentially expressed genes (with the lowest p-values from the limma model) representing each phase of embryogenesis. Similarly, mortality signature of early embryogenesis was compared with the signature of HPB in old animals.

#### Dynamics of biological age across individual cell lineages during early organogenesis

To assess dynamics of transcriptomic age across individual lineages during early organogenesis (from E6.5 to E8.5), we utilized scRNA-seq data from ^132^. To validate global decrease of biological age for the whole embryo during this interval of development, we aggregated all cells corresponding to each sample in a single metacell. We then filtered out metacells with total coverage less than 200K and preprocessed remaining meta-profiles using the standard pipeline for bulk RNA-seq data described above. Samples representing the earliest stage of development (E6.5) were selected as a reference group for relative expression calculation. Mouse chronological and mortality multi-tissue BR clocks were applied to the preprocessed metacell data, and the slope of tAge change with time was assessed with a linear mixed effect model.

To reconstruct ancestral lineages for all cell types presented at day 8.5, we utilized Waddington optimal transport method (wot package in python)^133^. Before inferring the trajectories, we preprocessed raw data following methodology described in ^133^. First, we conducted depth normalization of cell counts so that each cell had an equal total number of reads (denoted as CP10k), using normalize_total function in Scanpy^296^ with parameter target_sum=1e4. Next, we log-transformed the values using log(x+1) (log1p function in Scanpy). This combination of transformations is referred to as log1pCP10k and is necessary to control for dispersion and read depth in the data. Then we identified 3000 most highly variable genes, using highly_variable_genes function with parameters n_top_genes=3000, flavor=“seurat_v3” in Scanpy.

To infer trajectories, we first computed the average z-score for each cell based on the expression of genes involved in cell cycle and apoptosis. For each gene, we calculated z-score, truncated it at −5 or 5, and then computed the average z-score across all genes in the set for each cell, using score_gene_sets function with parameters permutations=0, method=“mean_z_score” in the wot library. Using the resulting values, representing signatures of apoptosis and proliferation, we assessed cell growth rate g(x) using a Birth-Death Process model. We then calculated transport matrices linking each pair of adjacent time points. We used OTModel function from the wot library with parameters epsilon = 0.05, lambda1 = 1, lambda2 = 50, growth_iters=3. The calculated transport matrices reflect ancestors and descendants of each cell. To determine the probabilistic distribution of ancestors of the cell populations present at E8.5, we first calculated probability vectors of these cell types using population_from_cell_sets function from the wot library with parameter at_time=8.5. The probability vectors of ancestor cell types were calculated by multiplying the cell type probability vector with the transport matrix using the wot trajectories function. Thus, we obtained ancestral trajectories of all cell types at E8.5, representing a sequence of probability distributions across individual cells at earlier time points.

To validate ancestral lineages reconstructed from the data, for every embryonic tissue at E8.5 we examined top cell types at each stage of development that together covered more than 90% of its total ancestral probability (Extended Data Fig. 23). Besides, we estimated pairwise Spearman correlation between ancestral lineages for each pair of cell types presented at E8.5 (Extended Data Fig. 24). Utilized trajectories represented probability distributions of cells that cumulatively explained more than 99% of total ancestral probability for the given cell type.

Reconstructed trajectories were utilized to estimate tAge dynamics for individual cell lineages between E6.5 and E8.5. To ensure sufficient gene coverage at E8.5, we selected cell types represented by at least 400 cells at this time point. Additionally, for each cell type, we filtered out cells that had zero probability for the given trajectory. To account for the probabilistic distribution of ancestors derived from our analysis, for each cell at earlier time points we multiplied its estimated probability by the gene expression profile. Thus, the counts of each gene in a specific cell were weighted according to its probability of being a part of the trajectory, i.e., being an ancestor of the corresponding cell type from E8.5. Therefore, low probability of being a member of ancestral lineage resulted in a reduced impact of cell’s expression profile on the meta-profile of the corresponding metacell and, therefore, lower effect on estimated tAge. We then aggregated weighted gene expression profiles of all cells within each sample into a single metacell, similar to the pipeline utilized for evaluation of tAge for the whole embryos, and multiplied the resulting aggregated weighted counts by the number of cells at the corresponding time point to get total expression values on the similar scale to the original ones. We filtered out meta-cells with total coverage less than 200K reads and preprocessed the data for clock analysis, utilizing samples at E6.5 as a reference group for calculation of relative expression. We then assessed tAge of each metacell using mouse chronological and mortality multi-tissue BR clocks and examined change of tAge between E6.5 and E8.5 for every cell lineage using a linear mixed-effect model. In addition, we computed Pearson correlation between the predicted point estimate of tAge and the day of development. P-values representing slope difference from zero estimated with mixed effect models were adjusted for multiple comparisons using BH method.

### Rejuvenation-associated signature of human iPSCs

To examine the change of tAge during cellular reprogramming of human fibroblasts and association between biological age of induced pluripotent stem cells (iPSCs) and chronological age of their donors, we utilized E-MTAB-3037 dataset^142^. Following standard preprocessing and mapping of human genes to corresponding mouse orthologs, we calculated relative expression using all primary fibroblasts samples as a reference group and applied multi-species multi-tissue BR clocks of chronological age and mortality. For every cell type presented in the data, including prefrontal cortex, primary fibroblasts, iPSCs and directly transdifferentiated neurons, we estimated association between tAge of samples and chronological age of donors, from whom the samples were derived, using linear mixed effect model. Besides, we calculated paired difference in biological age between original primary fibroblasts and iPSCs derived from these cells, using mixed effect model with patient ID incorporated there as a separate factorial covariate. Similar model was used to identify genes differentially expressed between primary fibroblasts and corresponding reprogrammed iPSCs. logFC were multiplied by the corresponding coefficients of multi-species multi-tissue EN clocks, and Pearson correlation between signatures of cellular reprogramming of human fibroblasts, HPB in old mice and early embryogenesis (up to E10) weighted by coefficients of multi-species mortality clock was assessed based on the union of top 2,000 differentially expressed genes (with the lowest p-value) for each pair of models. Top genes with the highest rejuvenation effect across models were estimated based on average gene association with mortality normalized by standard deviation (Fig. 7E).

### Screening of anti-aging compounds with Clockbase

To identify interventions with significant effect on molecular mechanisms of aging and mortality, we utilized Clockbase collection^147^ of 2,487 mouse tissue RNA-seq datasets from GEO repository. To focus on systemic effects of interventions on animal’s organs, we filtered out datasets representing cell culture experiments, keeping only gene expression profiles of mouse tissues. Following standard preprocessing of transcriptomic data, we calculated relative expression considering all samples within the dataset as a reference group. Afterwards, we utilized rodent multi-tissue EN clocks of chronological age, lifespan-adjusted age and expected mortality. For each dataset, we aggregated ANOVA model by introducing age, sex and tissue (if available) as well as all columns that included at least 2 non-unique and 2 different factorial or numerical values (‘experimental columns’). This threshold was applied to remove columns with unique sample IDs as well as columns specifying technical information shared across all the samples within the dataset. Using ANOVA test, for every experimental column within each dataset we calculated p-value reflecting statistical significance of tAge variation across the groups covered by the corresponding column. Following p-value adjustment with BH method, we filtered out datasets with not statistically significant variation according to at least one experimental column (BH adjusted p-value > 0.05). For the remaining datasets, we repeated ANOVA, examining pairwise comparisons between control and experimental groups after adjustment for age, sex, tissue and other available experimental columns. As control groups, we considered strings containing the following keywords: ‘wt’, ‘wild type’, ‘wild-type’, ‘control’, ‘ctrl’, ‘naïve’, ‘none’, ‘sham’, ‘dmso’, ‘pbs’, ‘ad libitum’, ‘untreat’, ‘no treat’, ‘unstimulated’, ‘intact’, ‘healthy’, ‘vehicle’, ‘veh’ (whole string), ‘empty’, ‘mock’, ‘saline’, ‘negative’, ‘neg’ (whole string), ‘no’ (whole string), ‘flox’, ‘standard’, ‘normal’, ‘chow’, ‘scramble’, ‘uninfected’. Resulting p-values representing pairwise comparisons between control and treated groups across datasets were adjusted for multiple comparisons with BH approach. Experimental groups with BH-adjusted p-value < 0.05 were considered significant. Top hits identified during this screening were verified by manual preprocessing and analysis of the corresponding datasets.

Gene expression profiles of luminal mature, luminal progenitor and mammary stem cells from 3-month-old ovariectomized mice treated with Estrogen and Progesterone (EP), EP and telapristone (TPA), EP and mifepristone (MFP), or sham for 14 days were downloaded from GSE127197^151^. Data were preprocessed with a standard pipeline, with sham samples specified as a reference group for calculation of relative expression. Resulting preprocessed profiles were passed to rodent chronological and mortality multi-tissue BR clocks. Pairwise differences between sham and EP, EP and TPA, and EP and MFP groups were assessed with mixed effect model separately for each cell type. Resulting p-values were adjusted for multiple comparisons using BH procedure.

Transcriptomic profiles of livers from young 7-week-old untreated C57BL/6J female mice, and old 24-month-old mice subjected to saline injections or 3-month treatment with ouabain were obtained from GSE122080^152^. Data were preprocessed with a standard pipeline, and young control animals were selected as a reference group for calculation of relative expression profiles. tAges of tissues were estimated with conventional and module-specific rodent multi-tissue transcriptomic clocks of chronological age and mortality trained on independent data, not intersecting with the examined dataset. Pairwise differences in conventional tAge estimated with BR models between young and old control groups, and between old animals subjected to saline and ouabain, were assessed with mixed-effect models. P-values were adjusted for multiple comparisons using BH procedure. Differences in tAge between old control and ouabain-treated animals revealed by individual module-specific clocks were assessed with ANOVA model. Same statistical model was used to assess module contributions to observed changes in biological age induced by ouabain treatment. To identify top genes driving rejuvenation effect of ouabain in old animals, differential expression analysis between old control and ouabain groups was performed, and the resulting logFC were multiplied by corresponding coefficients of the conventional rodent mortality multi-tissue EN clock. Genes were ranked by absolute value of weighted mortality-associated changes, and top 25 genes were visualized on a barplot. Pearson correlation between mortality-associated signatures of ouabain and HPB in old mice was assessed based on the union of top 2,000 differentially expressed genes (with the lowest p-value) across examined models.

