## Supplementary figures for "Transcriptomic Hallmarks of Mortality Reveal Universal and Specific Mechanisms of Aging, Chronic Disease, and Rejuvenation"

### Extended Data Figures

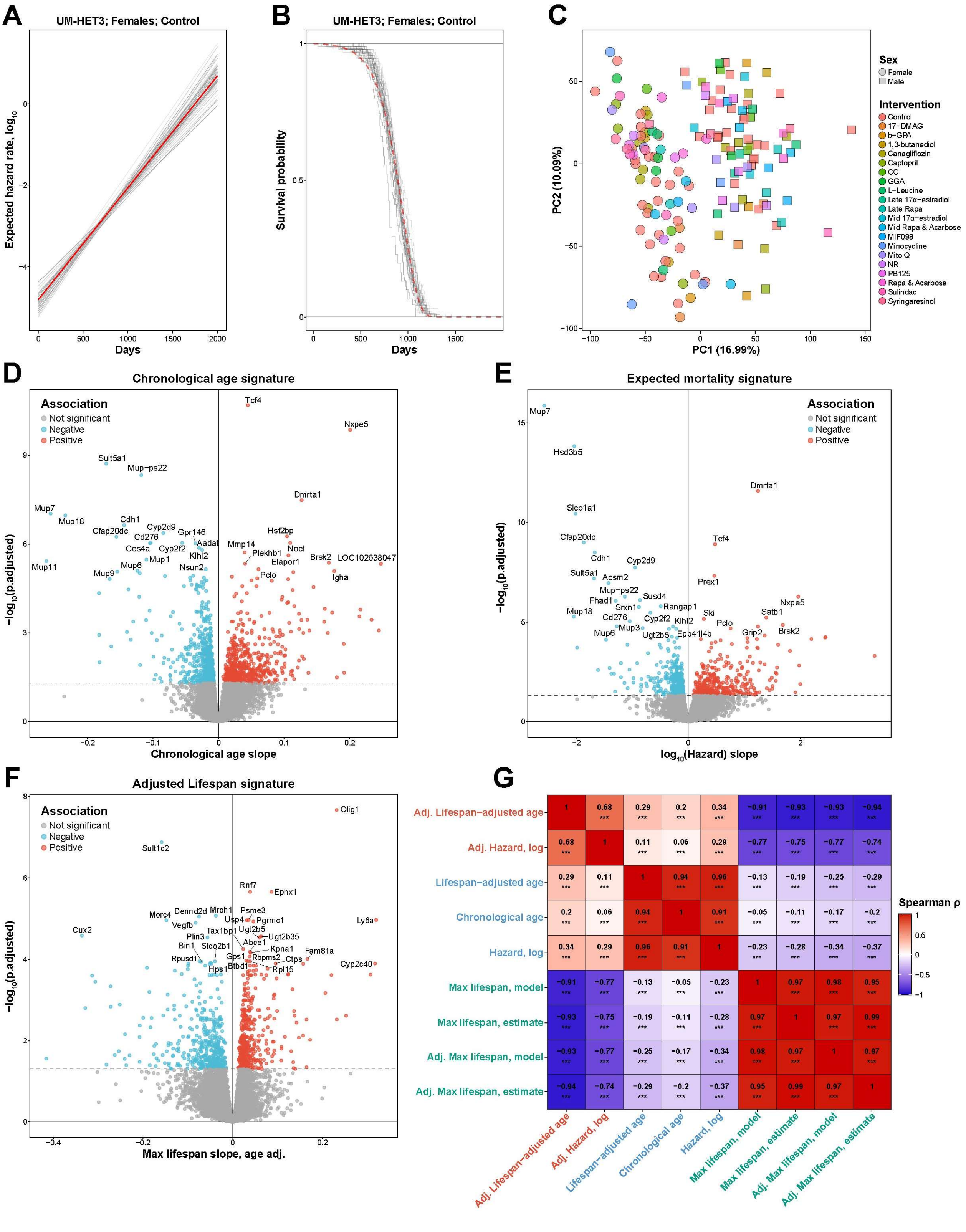

**Extended Data Fig. 1. Gene expression signatures of aging, mortality, and maximum lifespan in the Interventions Testing Program (ITP) cohorts.**

A. Meta-mortality model (red) based on individual Gompertz models fitted on different control female UM-HET3 cohorts (grey). Meta-estimates of Gompertz model parameters were derived with mixed-effect model.

B. Meta-survival curve (red) of control female UM-HET3 mice and corresponding individual survival curves (grey).

C. Principal component analysis (PCA) of samples from ITP data. Sexes and interventions are shown with shapes and colors, respectively. Rapa: Rapamycin; CC: Candesartan Cilexetil; GGA: Geranylgeranyl Acetone; NR: Nicotinamide Riboside; Late: Late-life; Mid: Mid-life.

D. Gene expression signatures of chronological age in control animals from ITP dataset (n=55). Slope of association and BH-adjusted p-value (in log scale) are shown on x and y axis, respectively. Top genes associated with age are shown in text.

E. Gene expression signatures of expected mortality of animals from ITP dataset (n=161). Expected mortality was estimated with Gompertz models fitted for corresponding sex and intervention group.

F. Gene expression signatures of maximum lifespan adjusted for chronological age of animals from ITP dataset (n=161). Expected maximum lifespan ($90^{th}$ percentile) was estimated from survival data for corresponding sex and intervention group.

G. Correlation between gene expression signatures of maximum lifespan (green), chronological age, lifespan-adjusted age and mortality unadjusted (blue) and adjusted for chronological age (red) identified for animals from ITP dataset. Correlation coefficient and BH-adjusted p-values are shown in text and asterisks, respectively. * p.adj < 0.05; ** p.adj < 0.01; *** p.adj < 0.001.

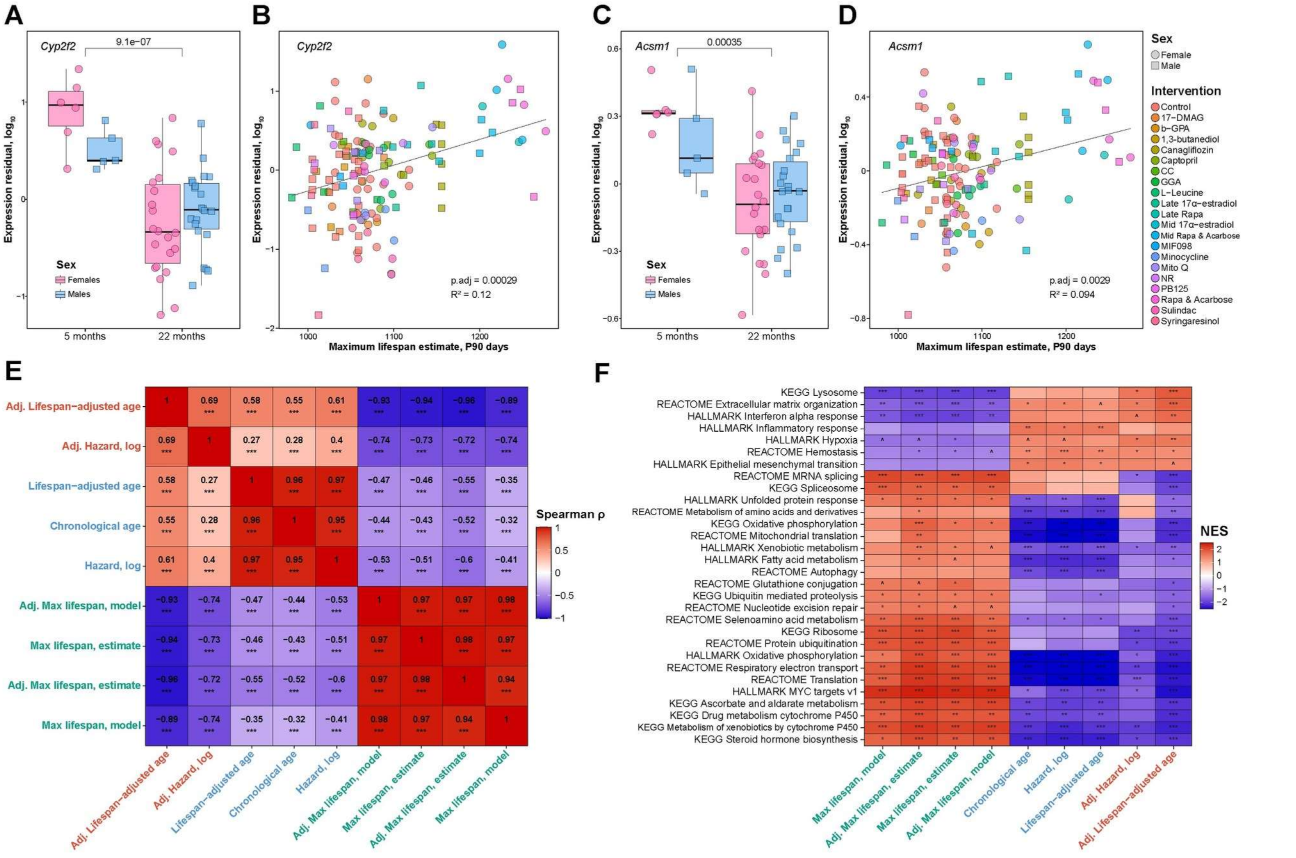


**Extended Data Fig. 2. Representative genes and pathways associated with aging, mortality, and maximum lifespan in the Interventions Testing Program (ITP) cohorts.**

A. Expression of *Cyp2f2* adjusted for sex in young (left) and old (right) control UM-HET3 mice. BH adjusted p-value corresponding to age difference in expression is shown in text.

B. Association between expected maximum lifespan of ITP cohorts estimated from survival data (x axis) and expression of *Cyp2f2* adjusted for sex (y axis).

C. Expression of *Acsm1* adjusted for sex in young (left) and old (right) control UM-HET3 mice.

D. Association between expected maximum lifespan of ITP cohorts estimated from survival data (x axis) and expression of *Acsm1* adjusted for sex (y axis).

E. Correlation matrix of normalized enrichment scores (NES) of pathways associated with signatures of maximum lifespan (green), chronological age, lifespan-adjusted age and mortality unadjusted (blue) and adjusted for chronological age (red), identified from ITP dataset. NES were estimated with GSEA. Correlation coefficient and BH-adjusted p-values are shown in text and asterisks, respectively.

F. Functional enrichment (GSEA) of transcriptomic signatures of maximum lifespan (green), chronological age, lifespan-adjusted age and mortality unadjusted (blue) and adjusted for chronological age (red), identified from ITP dataset. Statistical significance is shown with asterisks. Only functions significantly enriched by at least one signature are shown (BH-adjusted p-value < 0.05). The whole list of enriched functions is in Supplementary Table 2A. NES: Normalized Enrichment Score.

Rapa: Rapamycin; CC: Candesartan Cilexetil; GGA: Geranylgeranyl Acetone; NR: Nicotinamide Riboside; Late: Late-life; Mid: Mid-life. ^ p.adj < 0.1; * p.adj < 0.05; ** p.adj < 0.01; *** p.adj < 0.001.

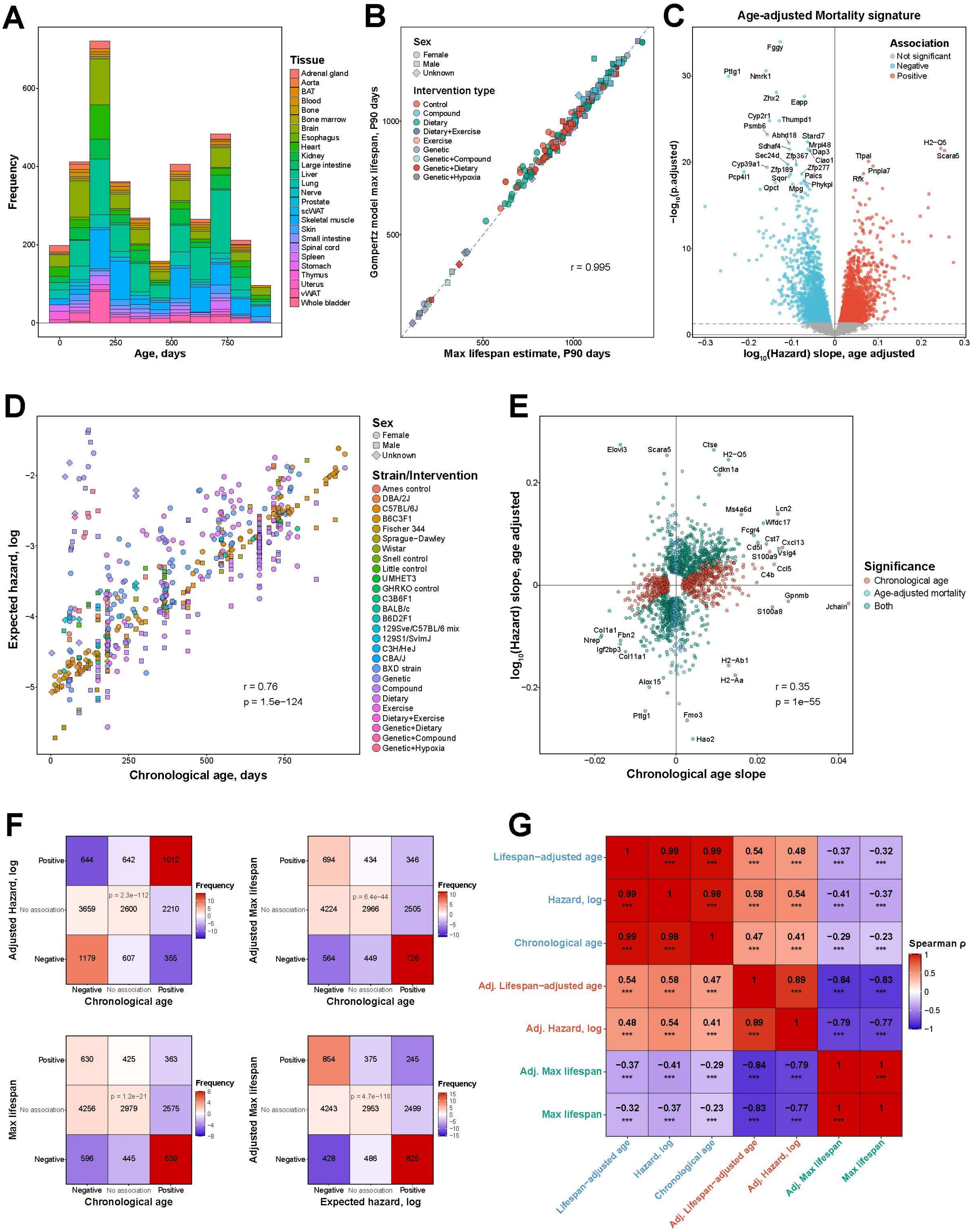


**Extended Data Fig. 3. Transcriptomic signatures of aging, mortality, and maximum lifespan in meta-dataset.**

A. Distribution of chronological age (x axis) and tissues (shown by color) of control mice and rats covered in the aggregated meta-dataset (n=3,575). BAT: Brown Adipose Tissue; scWAT: Subcutaneous White Adipose Tissue; vWAT: Visceral White Adipose Tissue.

B. Estimates of expected maximum lifespan (90$^{th}$ percentile) for various interventions, strains and sexes presented in aggregated meta-dataset derived from survival data (x axis) and from fitted

Gompertz models (y axis). Pearson correlation coefficient and p-value are shown in text. Type of intervention is denoted with color.

C. Gene expression signatures of mortality adjusted for chronological age identified from the whole meta-dataset (n=4,539). Expected hazard was estimated with Gompertz model based on survival data for the corresponding sex, strain and intervention. Slope of association and BH-adjusted p-value (in log scale) are shown on x and y axis, respectively. Top genes associated with mortality are shown in text.

D. Association between chronological age and expected mortality (in log scale) for all intervention, strain and sex models of mice and rats presented in the aggregated meta-dataset. Strains and intervention are denoted with color, sex is denoted with shapes. Pearson correlation coefficient and corresponding p-value are shown in text.

E. Association between gene expression signatures of chronological age and age-adjusted mortality identified in meta-dataset. The union of top 1,000 genes associated with age and age-adjusted mortality (with the lowest p-value) are shown on the plot. Genes significantly associated (BH-adjusted p-value $< 0.05$) with chronological age, age-adjusted mortality or both of these traits are shown in red, blue and green, respectively. Pearson correlation coefficient and corresponding p-value are shown in text.

F. Overlap of age-, mortality- and lifespan -associated genes. Number of genes and deviation from the expected random overlap are indicated with text and color, respectively. p value was estimated with Pearson's chi-square tests.

G. Correlation between gene expression signatures of maximum lifespan (green), chronological age, lifespan-adjusted age and mortality unadjusted (blue) and adjusted for chronological age (red) identified for animals from meta-dataset. The union of top 1,000 genes with the lowest p-value for each pair of signatures was used to calculate Spearman correlation coefficient. Correlation coefficient and BH-adjusted p-values are shown in text and asterisks, respectively. * p.adj < 0.05; ** p.adj < 0.01; *** p.adj < 0.001.

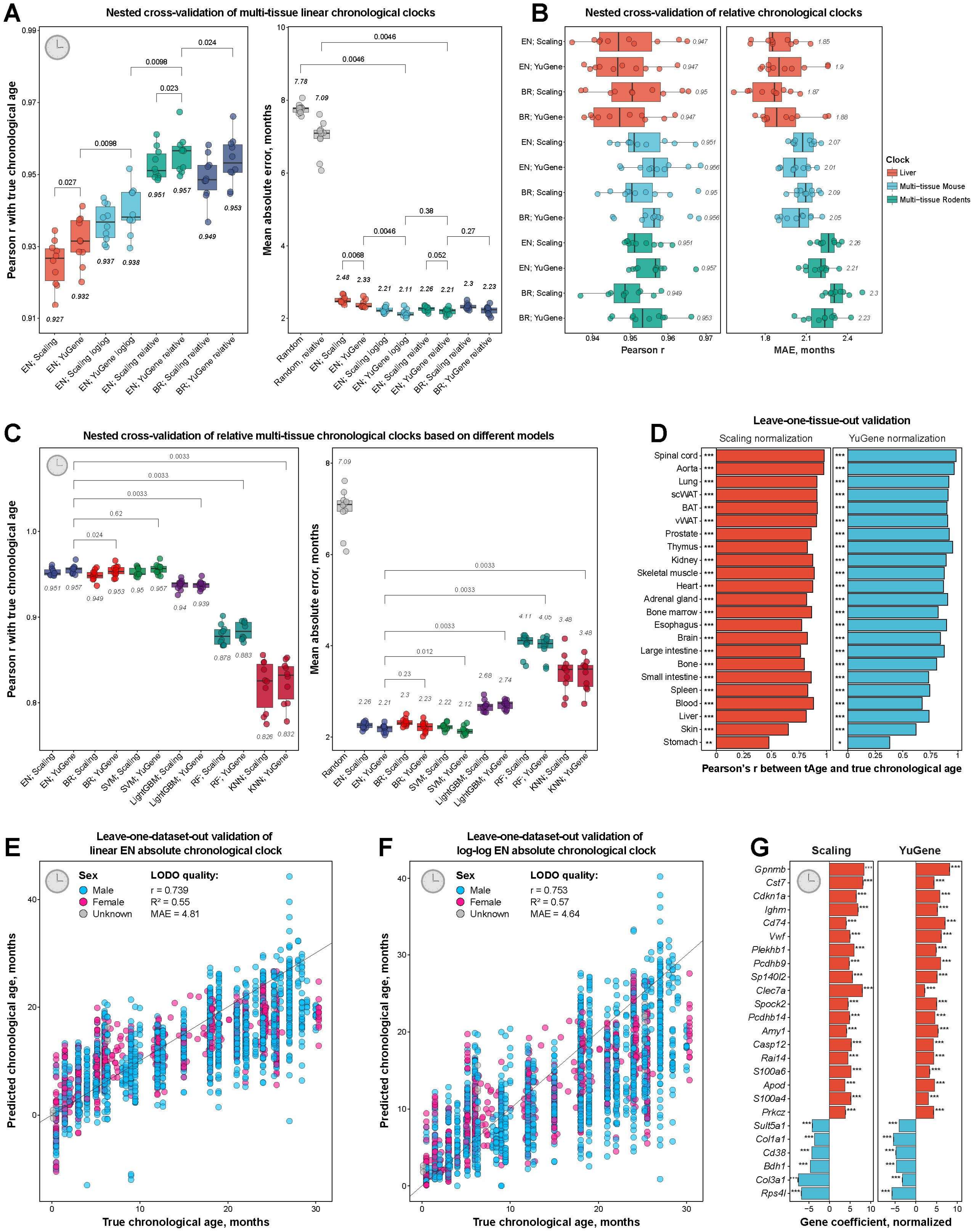


**Extended Data Fig. 4. Quality of chronological transcriptomic clocks.**

A. Accuracy of predictions of absolute age, log-log age (loglog) and relative chronological age (relative) on 10 randomly chosen test sets for rodent multi-tissue transcriptomic clocks with Elastic Net (EN) or Bayesian Ridge (BR) model with scaling or YuGene normalization. Pearson correlation coefficient and MAE are shown on left and right panels, respectively. Median estimate of quality across 10 runs is provided in text. MAE of random age prediction is shown in grey.

Pairwise comparison of accuracy for various models was performed with Wilcoxon signed-rank test, corresponding BH-adjusted p-values are shown in text.

B. Accuracy of predictions of relative chronological age on 10 randomly chosen test sets for liver (red) and multi-tissue mouse (blue) or rodent (green) transcriptomic clocks with EN or BR model with scaling or YuGene normalization. Median Pearson correlation coefficient is shown in text.

C. Accuracy of predictions of relative chronological age on 10 randomly chosen test sets for rodent multi-tissue transcriptomic clocks trained with various machine learning models. Pairwise comparison of accuracy for various models was performed with Wilcoxon signed-rank test, corresponding BH-adjusted p-values are shown in text. EN: Elastic Net; BR: Bayesian Ridge; SVM: Support Vector Machines; LightGBM: Light Gradient-Boosting Machine; RF: Random Forest; KNN: K-Nearest Neighbors.

D. Quality of relative chronological age prediction for novel tissues not included in the training set with rodent multi-tissue EN transcriptomic clock. Pearson correlation between real and predicted age was estimated for each organ after training on remaining tissues (leave-one-tissue-out procedure) with scaling (left) and YuGene (right) normalization methods. Statistical significance of correlation is denoted with asterisks.

E-F. Leave-one-dataset-out (LODO) accuracy of absolute linear chronological age (E) and -log(-log(Age)) (F) prediction with rodent multi-tissue EN transcriptomic clocks (with YuGene normalization). Sex of samples is depicted with color. Pearson correlation coefficient, $R^2$ and mean absolute error (MAE) are shown in text.

G. Top 25 genes with the highest average absolute coefficients in rodent multi-tissue EN chronological clocks trained on data subjected to scaling (left) or YuGene (right) normalization across 10 randomly chosen training sets. Data are mean normalized coefficients ± SE.

* p.adj < 0.05; ** p.adj < 0.01; *** p.adj < 0.001.

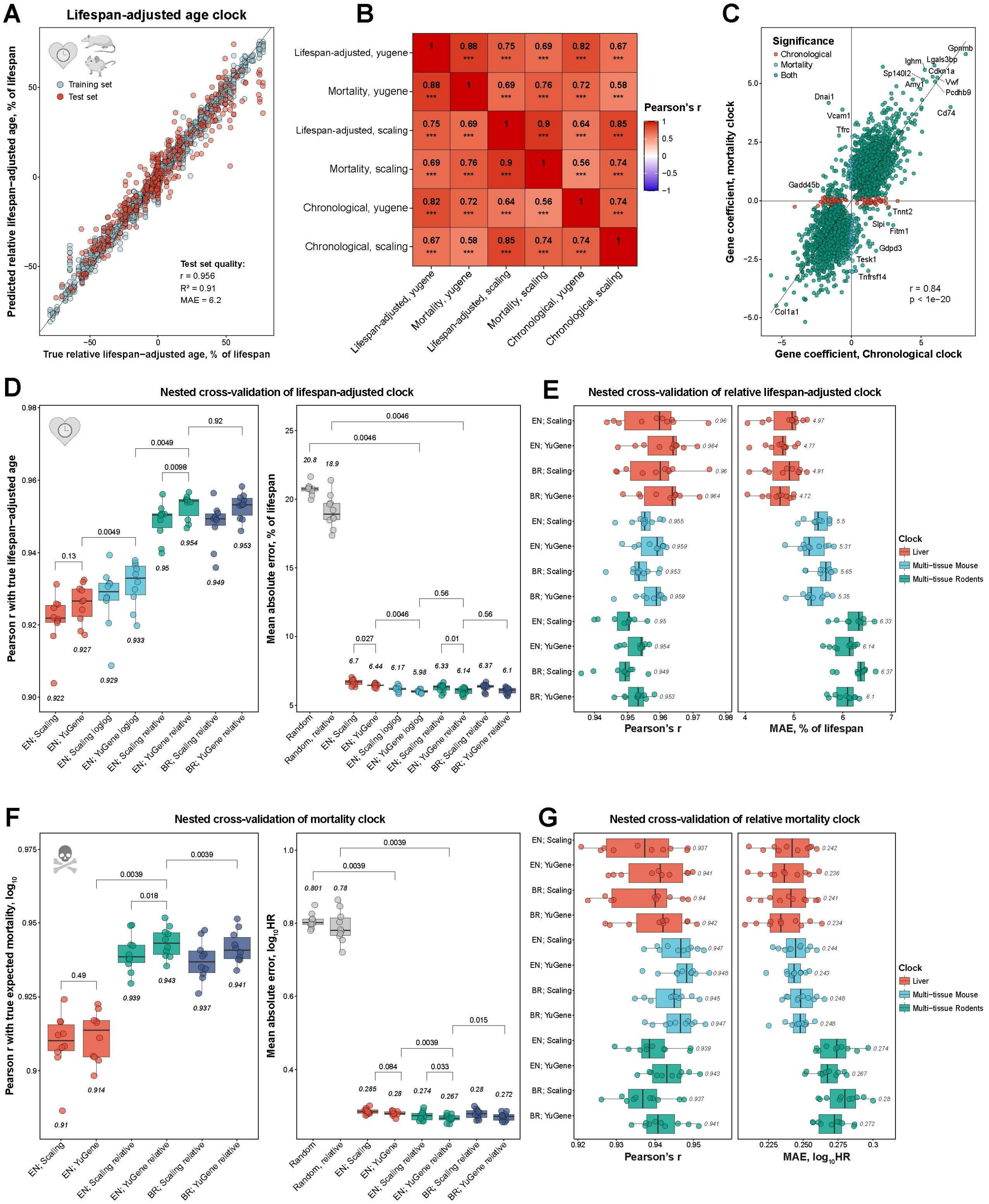


**Extended Data Fig. 5. Quality of transcriptomic clocks of lifespan-adjusted age and expected mortality.**

A. Accuracy of prediction of relative lifespan-adjusted age with multi-tissue Elastic Net (EN) transcriptomic clocks trained on mice and rats (with YuGene normalization). Training and test sets are denoted by color. Pearson correlation coefficient, $R^2$ and mean absolute error (MAE) for test set are shown in text.

B. Pearson correlation between average gene coefficients of rodent multi-tissue EN chronological, lifespan-adjusted and mortality clocks with scaling and YuGene normalization trained on 10 randomly chosen training sets. Correlation coefficient and BH-adjusted p-values are shown in text and asterisks, respectively. * p.adj < 0.05; ** p.adj < 0.01; *** p.adj < 0.001.

C. Association between gene coefficients of rodent multi-tissue EN chronological and mortality clocks (with YuGene normalization) trained on 10 randomly chosen training sets. The union of top 2,000 genes selected by chronological and mortality clocks (with the lowest p-values) are shown on the plot. Genes with coefficient significantly different from zero (BH-adjusted p-value < 0.05) in chronological, mortality or both clocks are shown in red, blue and green, respectively. Pearson correlation coefficient and corresponding p-value are shown in text.

D. Accuracy of predictions of absolute, log-log (loglog) and relative lifespan-adjusted age (relative) on 10 randomly chosen test sets for rodent multi-tissue transcriptomic clocks with EN or Bayesian Ridge (BR) model with scaling or YuGene normalization. Pearson correlation coefficient and MAE are shown on left and right panels, respectively. Median estimate of quality across 10 runs is provided in text. MAE of random prediction is shown in grey. Pairwise comparison of accuracy for various models was performed with Wilcoxon signed-rank test, corresponding BH-adjusted p-values are shown in text.

E. Accuracy of predictions of relative lifespan-adjusted age on 10 randomly chosen test sets for liver (red) and multi-tissue mouse (blue) or rodent (green) transcriptomic clocks with EN or BR model with scaling or YuGene normalization. Median Pearson correlation coefficient is shown in text.

F. Accuracy of predictions of absolute and relative expected hazard in log scaled on 10 randomly chosen test sets for rodent multi-tissue transcriptomic clocks with Elastic Net (EN) or Bayesian Ridge (BR) model with scaling or YuGene normalization. HR: Hazard Ratio.

G. Accuracy of predictions of relative mortality on 10 randomly chosen test sets for liver (red) and multi-tissue mouse (blue) or rodent (green) transcriptomic clocks with EN or BR model with scaling or YuGene normalization.

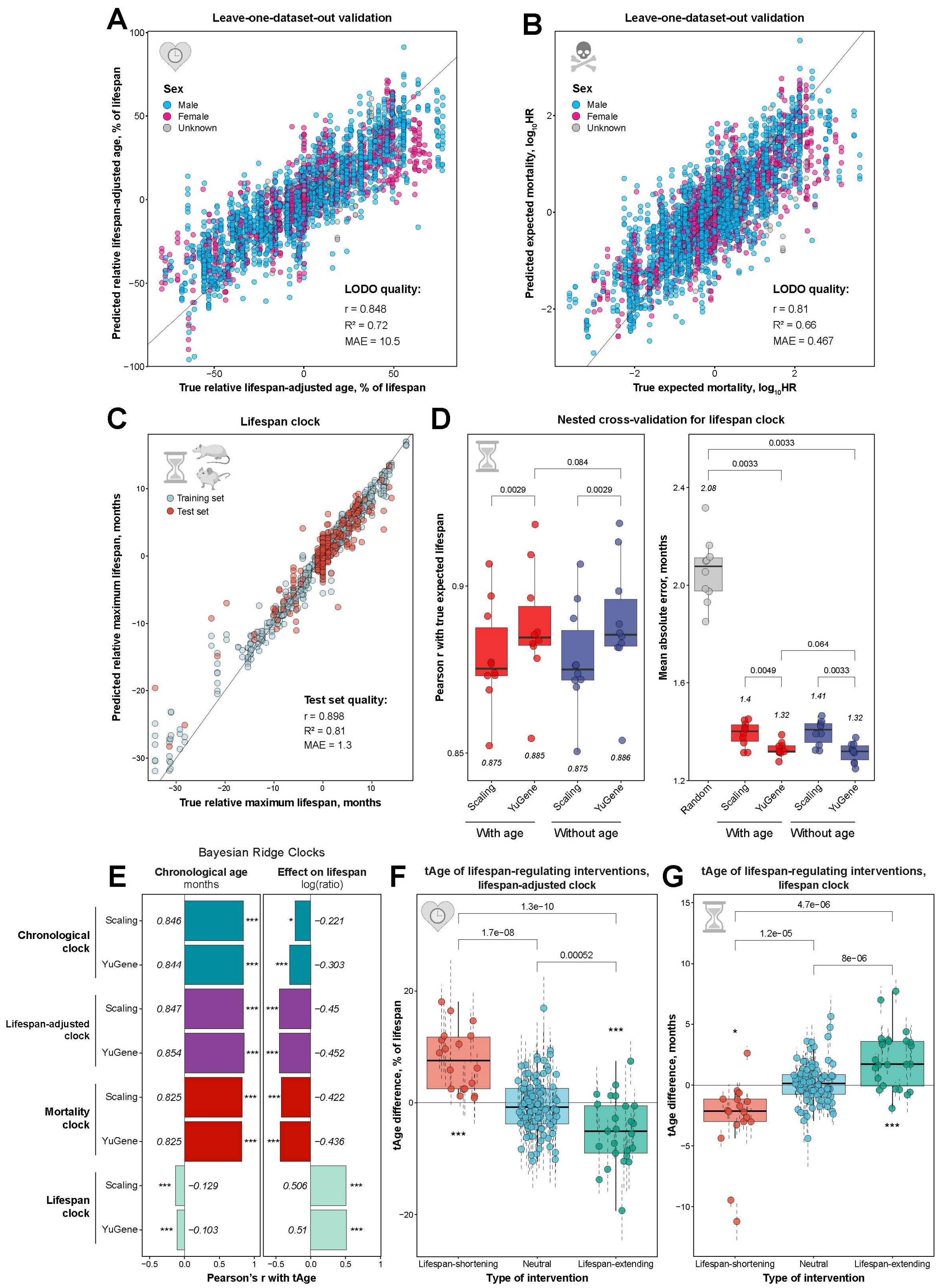


**Extended Data Fig. 6. Prediction of chronological age and effect on maximum lifespan with transcriptomic clocks.**

A-B. Leave-one-dataset-out (LODO) accuracy of prediction of relative lifespan-adjusted age (A) and mortality (B) with rodent multi-tissue Elastic Net (EN) transcriptomic clocks (with YuGene normalization). Sex of samples is depicted with color. Pearson correlation coefficient, $R^2$ and mean absolute error (MAE) are shown in text.

C. Accuracy of prediction of relative expected maximum lifespan with multi-tissue EN transcriptomic clocks trained on mice and rats (with YuGene normalization). Training and test sets are denoted by color. Pearson correlation coefficient, $R^2$ and mean absolute error (MAE) for test set are shown in text.

D. Accuracy of predictions of relative expected maximum lifespan on 10 randomly chosen test sets for rodent multi-tissue EN transcriptomic clocks with scaling or YuGene normalization with chronological age included (With age) or excluded (Without age) from the feature set. Pearson correlation coefficient and MAE are shown on left and right panels, respectively. Median estimate of quality across 10 runs is provided in text. MAE of random prediction is shown in grey. Pairwise comparison of accuracy for various models was performed with Wilcoxon signed-rank test, corresponding BH-adjusted p-values are shown in text.

E. Quality of prediction of chronological age (left) and effect on lifespan induced by various interventions (right) for rodent multi-tissue Bayesian Ridge transcriptomic clocks of chronological age, lifespan-adjusted age, mortality and expected maximum lifespan. Pearson correlation coefficient was calculated between predicted transcriptomic age and real chronological age (months) or effect on lifespan (log-ratio between treated and control animals) in independent datasets. Median Pearson's r and statistical significance are shown with text and asterisks, respectively.

F-G. Accuracy of prediction of intervention effect on expected lifespan for rodent multi-tissue EN transcriptomic clocks of lifespan-adjusted age (F) and maximum lifespan (G) (with YuGene normalization). For every type of interventions (lifespan-shortening, neutral or lifespan-extending), dots reflect mean tAge differences between treated and control samples for a given intervention, dataset and sex. Note that in case of the lifespan clock (G) tAge is positively correlated with expected lifespan and, therefore, has opposite association with predictions of mortality and lifespan-adjusted age clocks. Statistical significance of deviation from zero for lifespan-shortening and lifespan-extending interventions and pairwise comparison of tAges across the groups were assessed with mixed effect models. Corresponding BH-adjusted p-values are shown with asterisks and text, respectively. Data are mean tAge differences ± SE.

* p.adj < 0.05; ** p.adj < 0.01; *** p.adj < 0.001.

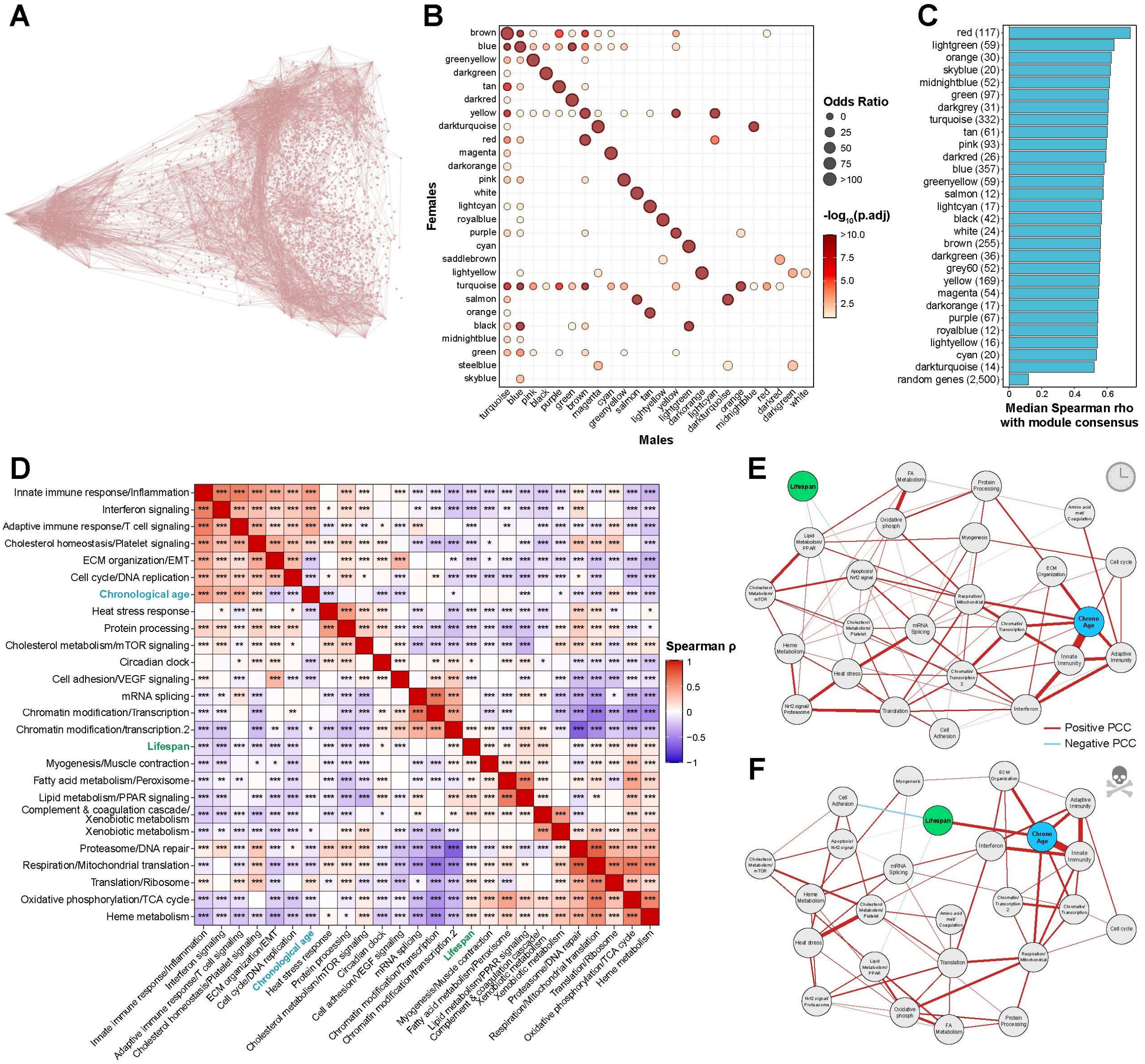


**Extended Data Fig. 7. Characterization of co-regulated gene expression modules of aging and longevity.**

A. Gene co-expression network of randomly chosen 3,000 genes based on spectral embedding and pairwise Pearson correlation estimates between the genes.

B. Enrichment of co-regulated gene modules identified separately for males (in rows) and females (in columns) with weighted gene co-expression network analysis (WGCNA). Significance of overlap for every pair of modules was assessed with Fisher's exact test. Size of dots and their color reflect odds ratio and BH-adjusted p-value (in log scale), respectively. Only dots for overlaps with adjusted p-value < 0.05 are shown.

C. Median Spearman correlation between expression of genes in every module and consensus expression profile for the respective module after filtering. Consensus expression is defined as a median expression of all genes within the given module. Correlation for the randomly chosen 2,500 genes not included in any module is shown at the bottom.

D. Partial correlation (adjusted for chronological age) between 1$^{st}$ Principal Components (PCs) of identified modules, expected maximum lifespan (green) and chronological age (blue). Color and asterisks reflect Spearman correlation coefficient and corresponding statistical significance, respectively. Modules are named after representative enriched functions. The dictionary between modules and functions is in Supplementary Table 4B. * p.adj < 0.05; ** p.adj < 0.01; *** p.adj < 0.001.

E-F. Partial correlation networks of chronological age (blue), expected maximum lifespan (green) and transcriptomic ages predicted on test sets by rodent multi-tissue chronological (E) and mortality (F) clocks trained on genes associated with individual modules (grey). Sign and scale of partial correlation coefficient (PCC) are indicated by color and line width.

AA: Amino acid; Chrono: Chronological; FA: Fatty acid; met: metabolism; ECM: Extracellular matrix; EMT: Epithelial-Mesenchymal Transition.

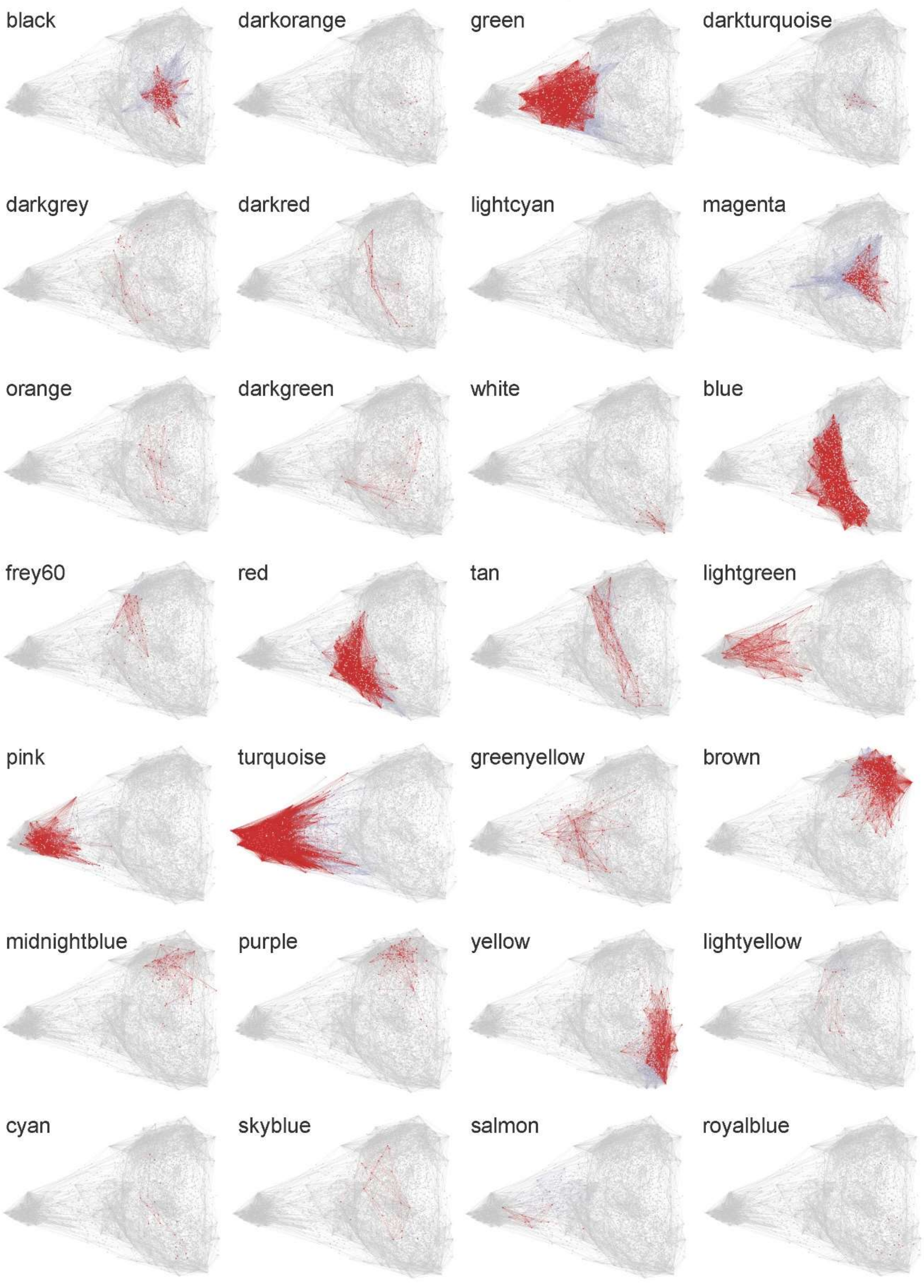


**Extended Data Fig. 8. Gene co-expression network based on spectral embedding colored by co-regulated modules identified with weighted gene co-expression network analysis (WGCNA).** Unfiltered and filtered modules are colored in blue and red, respectively.

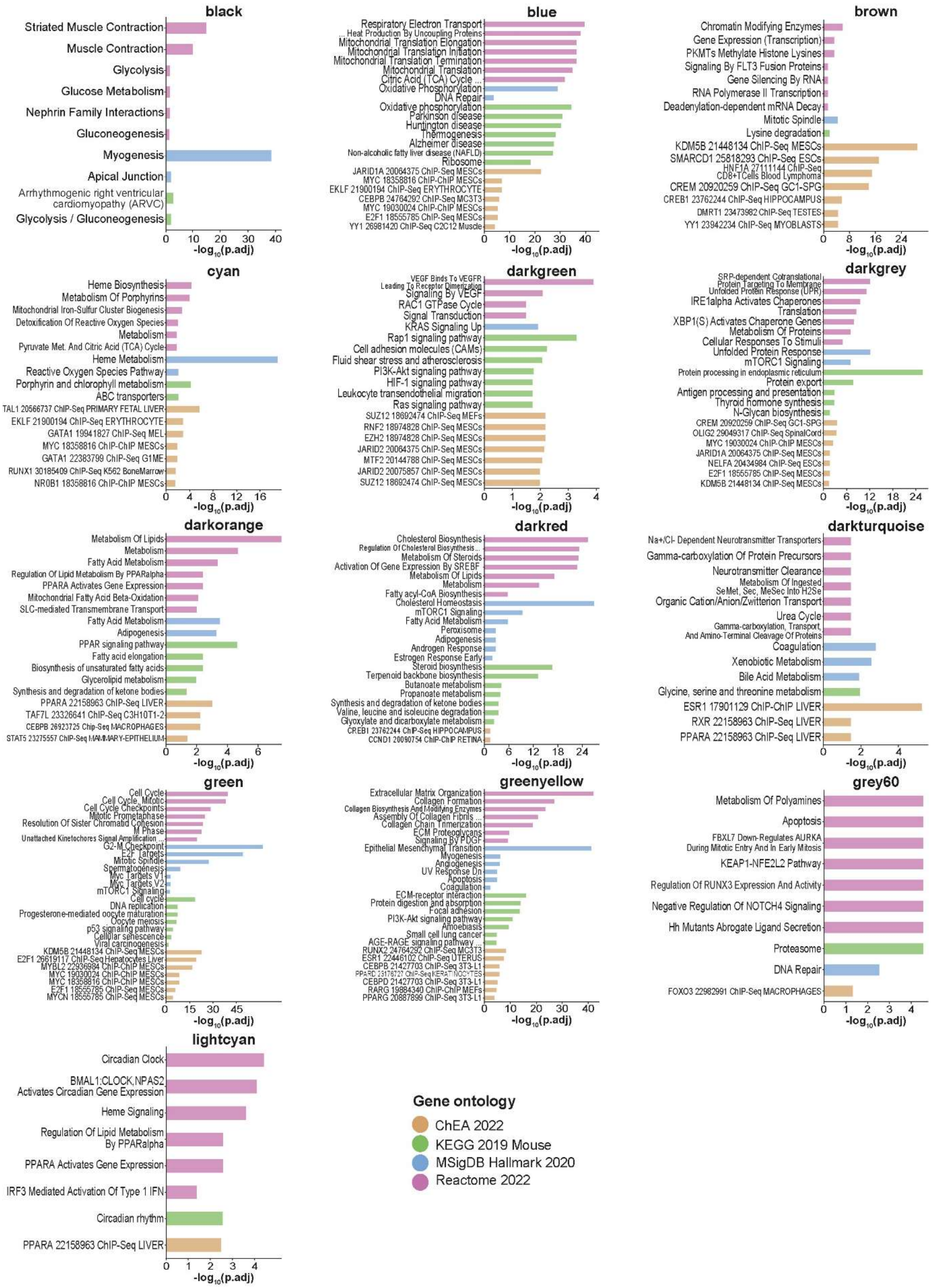


**Extended Data Fig. 9. Functional and upstream regulator enrichment of filtered modules identified with weighted gene co-expression network analysis (WGCNA).** Enrichment was performed with Fisher's exact test on KEGG, HALLMARK, REACTOME and ChEA ontologies. BH-adjusted p-values in log scale are shown on x axis. Module names are shown in bold, and colors reflect used ontology of gene sets. Only terms with adjusted p-value < 0.05 are shown.

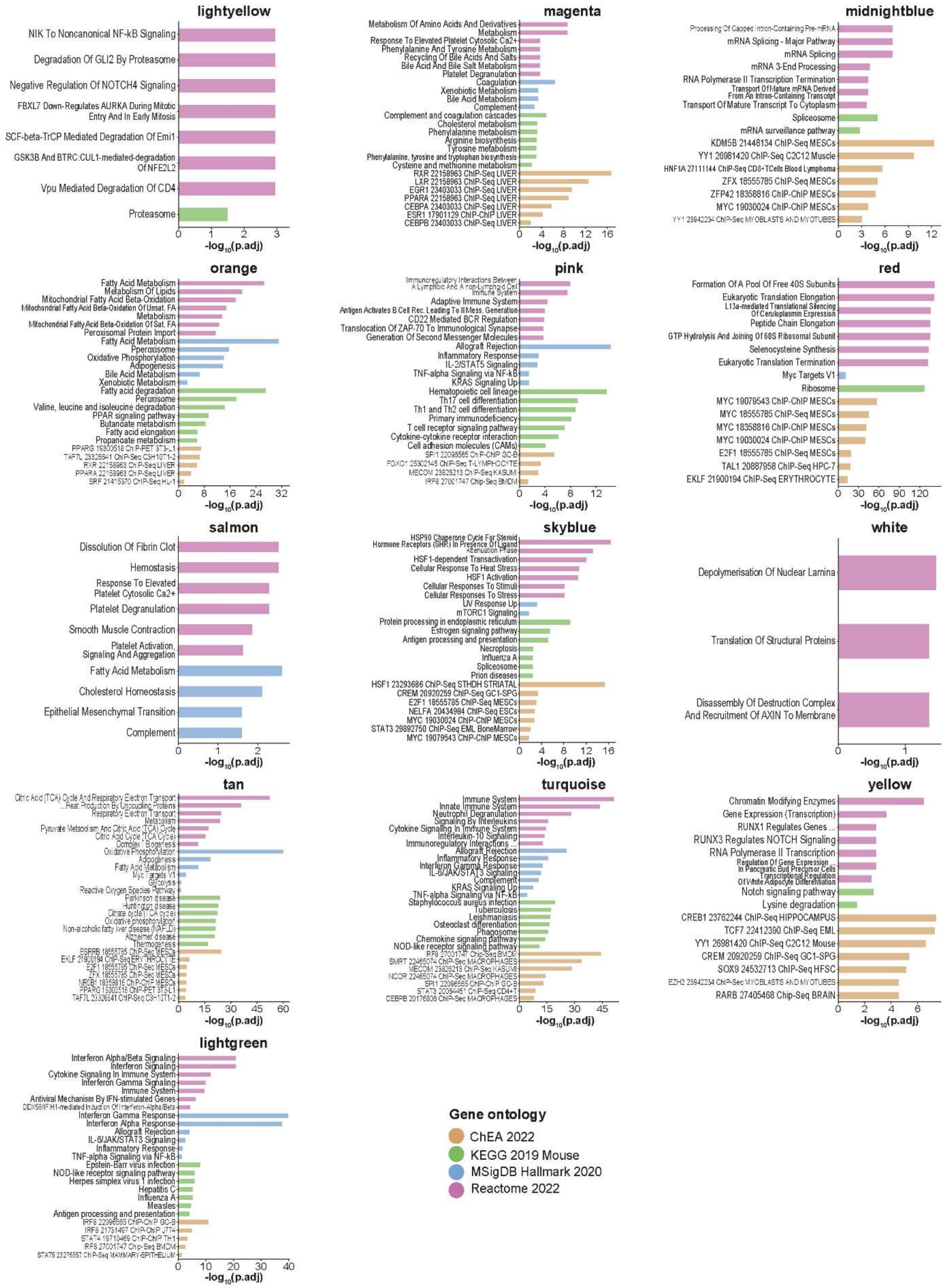


**Extended Data Fig. 10. Functional and upstream regulator enrichment of filtered modules identified with weighted gene co-expression network analysis (WGCNA) (continued).** Enrichment was performed with Fisher's exact test on KEGG, HALLMARK, REACTOME and ChEA ontologies. BH-adjusted p-values in log scale are shown on x axis. Module names are shown in bold, and colors reflect used ontology of gene sets. Only terms with adjusted p-value < 0.05 are shown.

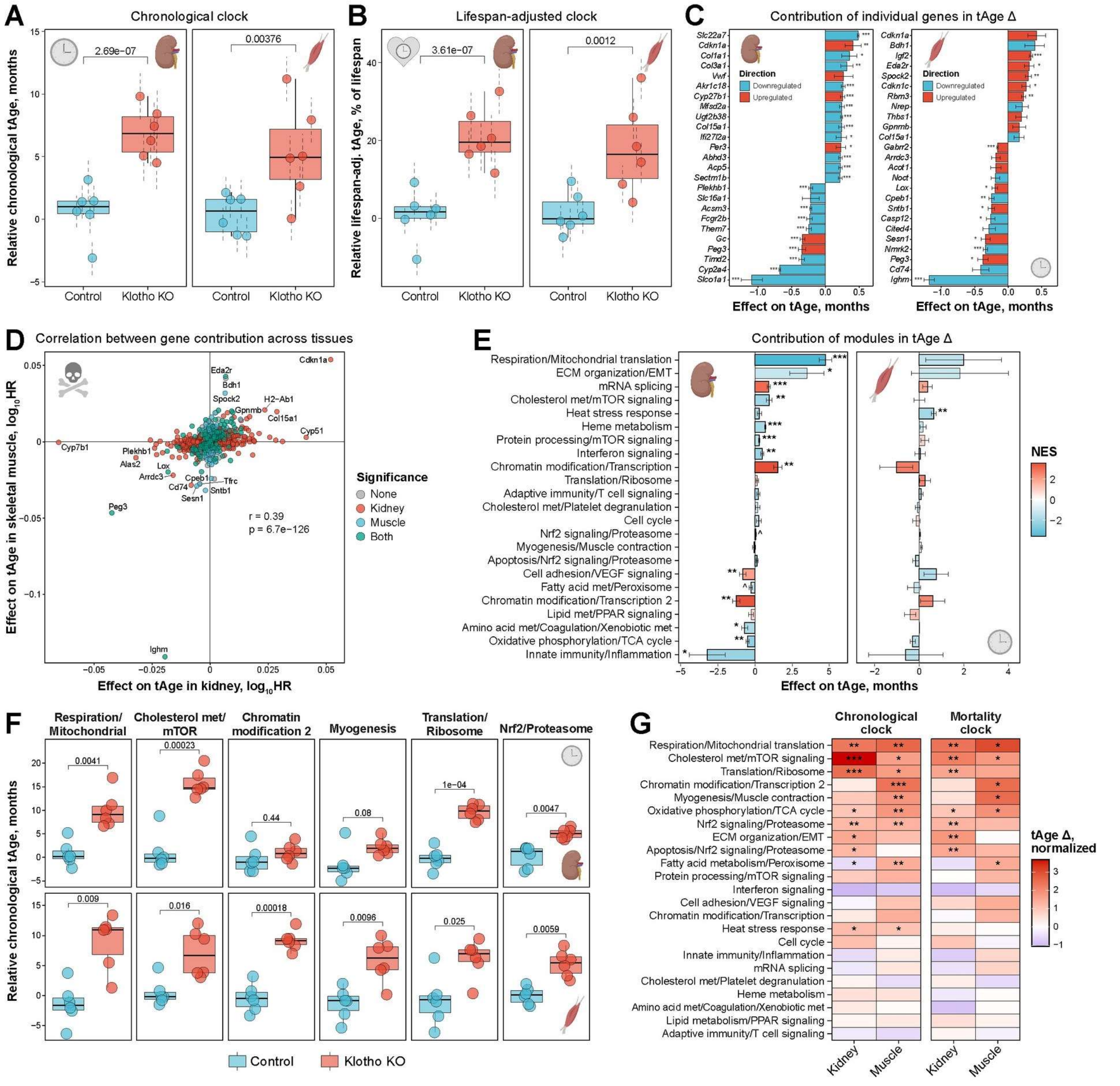


**Extended Data Fig. 11. Biological age and its molecular mechanisms in *Klotho* KO mice.**

A-B. Transcriptomic age (tAge) for control and *Klotho* KO mice in kidney (left) and muscle (right) estimated with rodent multi-tissue Bayesian Ridge (BR) chronological (A) and lifespan-adjusted (B) clocks trained on all genes excluding *Klotho*. tAges between the groups were compared with mixed effect model, corresponding BH-adjusted p-values are shown in text. Data are tAges ± SE.

C. Top genes driving pro- (positive) or anti-aging (negative) transcriptomic changes in kidneys (left) and skeletal muscles (right) of *Klotho* KO mice compared to age-matched controls, assessed with the rodent multi-tissue Elastic Net (EN) chronological clock. Top 25 genes with the highest absolute effect on tAge difference (logFC * clock coefficient) are shown. Genes up- and downregulated in *Klotho* KO mice are colored in red and blue, respectively. Statistical significance of logFC for each gene is indicated with asterisks. Data are mean tAge difference ± SE.

D. Association between weighted gene expression signatures of mortality (logFC * clock coefficient) in kidney and skeletal muscle of *Klotho* KO mice, according to the rodent multi-tissue EN mortality clock. The union of top 2,000 genes differentially expressed in *Klotho* KO (with the lowest p-values) in kidney and skeletal muscle are shown on the plot. Genes with statistically significant logFC (BH-adjusted p-value) in kidney, muscle or both tissues are colored in red, blue and green, respectively. Pearson correlation coefficient and corresponding p-value are shown in text. HR: Hazard Ratio.

E. Contributions of gene expression modules into tAge difference between control and *Klotho* KO mice estimated with the rodent multi-tissue EN chronological clock. Individual modules are shown in rows and names after representative functions. Positive and negative values reflect pro- and anti-aging changes in *Klotho* KO mice compared to age-matched controls, respectively. Statistical significance for each module was assessed with the two-sample unpaired t-test and denoted with asterisks. Bars are colored based on normalized enrichment scores (NES) from gene set enrichment analysis (GSEA), reflecting if genes associated with a particular module are generally up- (red) or downregulated (blue) in *Klotho* KO mice. Modules significantly enriched for up- or downregulated genes (BH-adjusted p-value $< 0.05$) are visualized with thick bars. Data are means ± SE.

F. Chronological tAge in control and *Klotho* KO mice estimated with representative module-specific chronological clocks. y axis reflects relative hazard ratios (in log scale) predicted with the respective multi-tissue module clocks. Difference between the groups was assessed with the two-sample unpaired t-test, and corresponding BH-adjusted p-values are shown in text.

G. Normalized chronological (left) and mortality (right) tAge difference between control and *Klotho* KO mice assessed with all module-specific multi-tissue chronological (left) and mortality (right) clocks. Color and asterisks reflect size and statistical significance of tAge difference between control and *Klotho* KO mice, assessed with the two-sample unpaired t-test. Increased and decreased tAge in *Klotho* KO mice is shown in red and blue, respectively.

ECM: Extracellular matrix; EMT: Epithelial-Mesenchymal Transition. ^ p.adj $< 0.1$; * p.adj $< 0.05$; ** p.adj $< 0.01$; *** p.adj $< 0.001$.

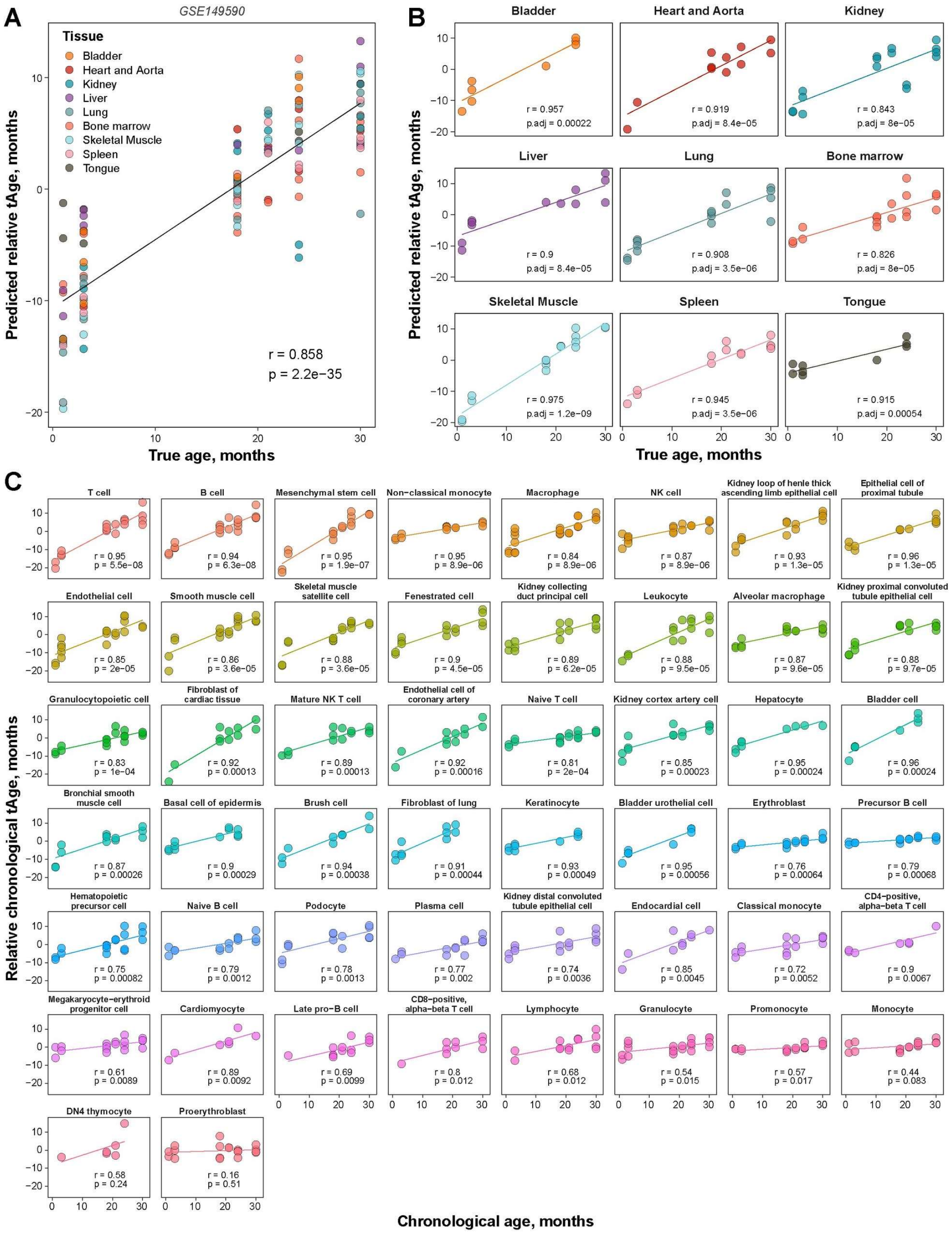


**Extended Data Fig. 12. Transcriptomic age dynamics in Tabula Muris Senis scRNA-seq data.**

A. Association between chronological age and transcriptomic age (tAge) of metacells across tissues estimated with mouse multi-tissue Elastic Net (EN) chronological clock. All cells corresponding to a specific sample and tissue were pooled into a metacell. Tissues are depicted with color. Pearson's correlation coefficient and corresponding p-value are shown in text.

B. Association between chronological age and tAge of metacells for individual tissues estimated with mouse multi-tissue EN chronological clock. Pearson's correlation coefficients and corresponding BH-adjusted p-values are shown in text.

C. Association between chronological age and tAge of metacells for individual cell types estimated with mouse multi-tissue EN chronological clock. All cells corresponding to a specific sample and tissue were pooled into a metacell. Pearson's correlation coefficients and corresponding BH-adjusted p-values are shown in text.

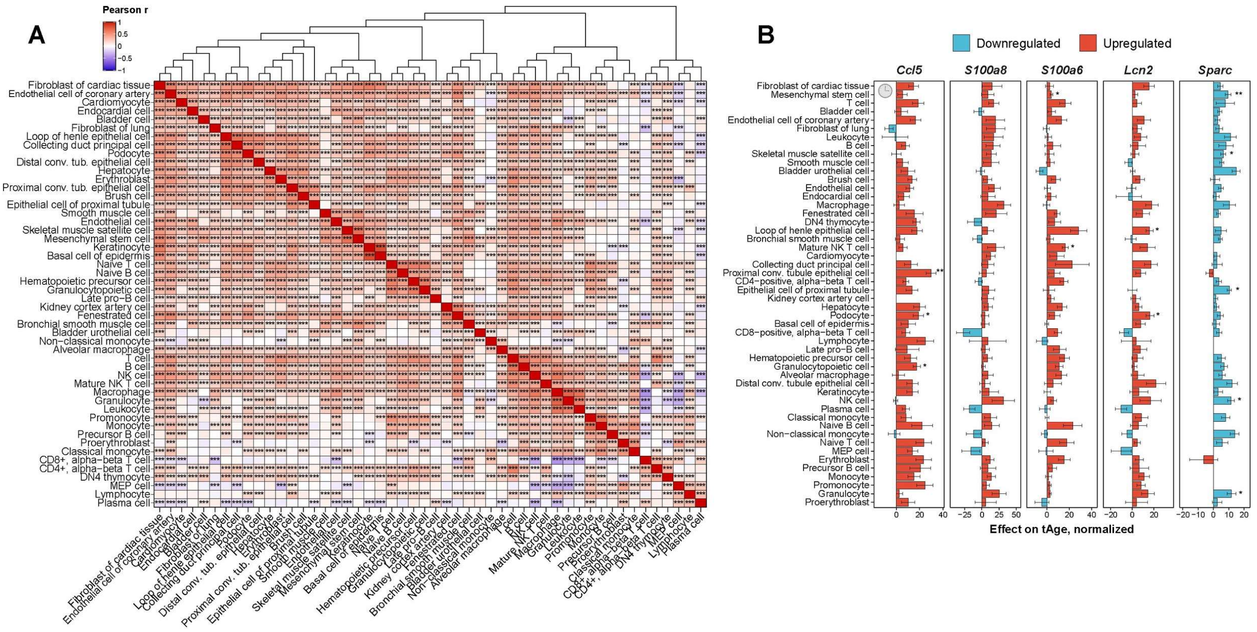


**Extended Data Fig. 13. Shared gene expression aging-associated biomarkers across cell types.**

A. Pearson correlation of weighted aging-associated gene expression signatures across individual cell types. The union of top 1000 genes associated with age (with the lowest p-value) were used to estimate Pearson's correlation coefficient for each pair of cell types. Weighted signatures were calculated as (age slope * chronological clock coefficient). Complete hierarchical method based on correlation distance was used for clustering. Statistical significance is depicted with asterisks.

B. Top genes contributing into increased molecular age across cell types according to mouse multi-tissue EN chronological clock. Top 5 genes with the highest average effect on normalized tAge (normalized slope * clock coefficient) across cell types are shown. Genes up- and downregulated with age are colored in red and blue, respectively. Statistical significance of slope for each gene is denoted with asterisks. Data are normalized tAge slope ± SE. MEP: Megakaryocytic-erythroid progenitors.

* p.adj < 0.05; ** p.adj < 0.01; *** p.adj < 0.001.

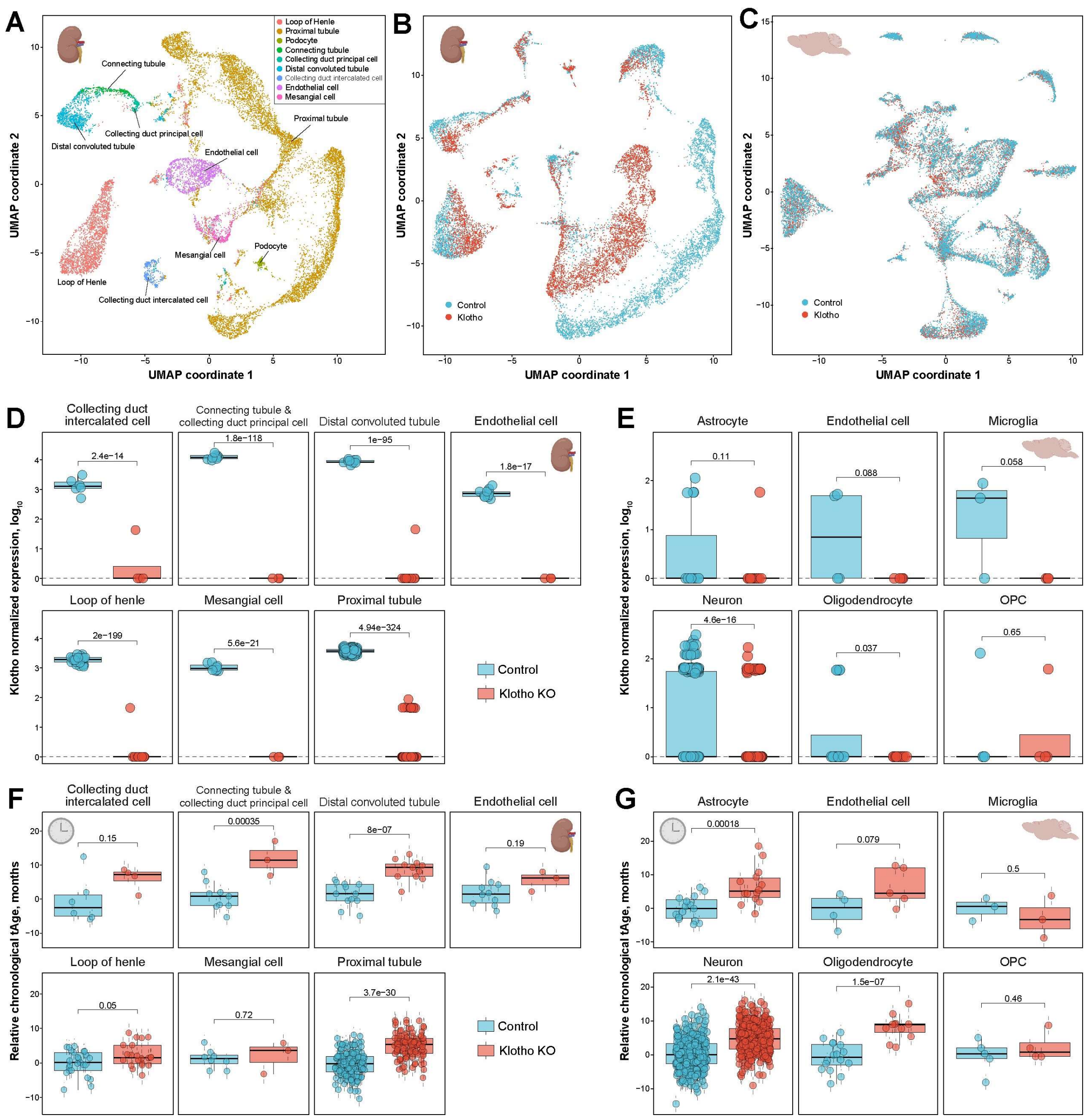


**Extended Data Fig. 14. Aging-associated dynamics of individual cell types in kidneys and brains of *Klotho* knockout mice.**

A. UMAP of unfiltered kidney cells from 8-week-old control and *Klotho* knockout (KO) mice. Cell type annotation is denoted with color.

B-C. UMAP of filtered kidney (B) and brain (C) cells from 8-week-old control and *Klotho* KO mice colored by experimental group.

D-E. Normalized expression of Klotho (in log scale) in metacells representing various kidney (D) and brain (E) cell types. Experimental group is denoted with color. Difference between groups was assessed with limma, and corresponding BH-adjusted p-values are shown in text.

F-G. tAge difference between metacells of control and age-matched *Klotho* KO mice representing various kidney (F) and brain (G) cell types estimated with the rodent multi-species Bayesian Ridge (BR) chronological clock trained on all genes excluding *Klotho*. tAges between the groups were compared with mixed effect model, and corresponding BH-adjusted p-values are shown in text. Data are tAges ± SE.

OPC: Oligodendrocyte Precursor Cells.

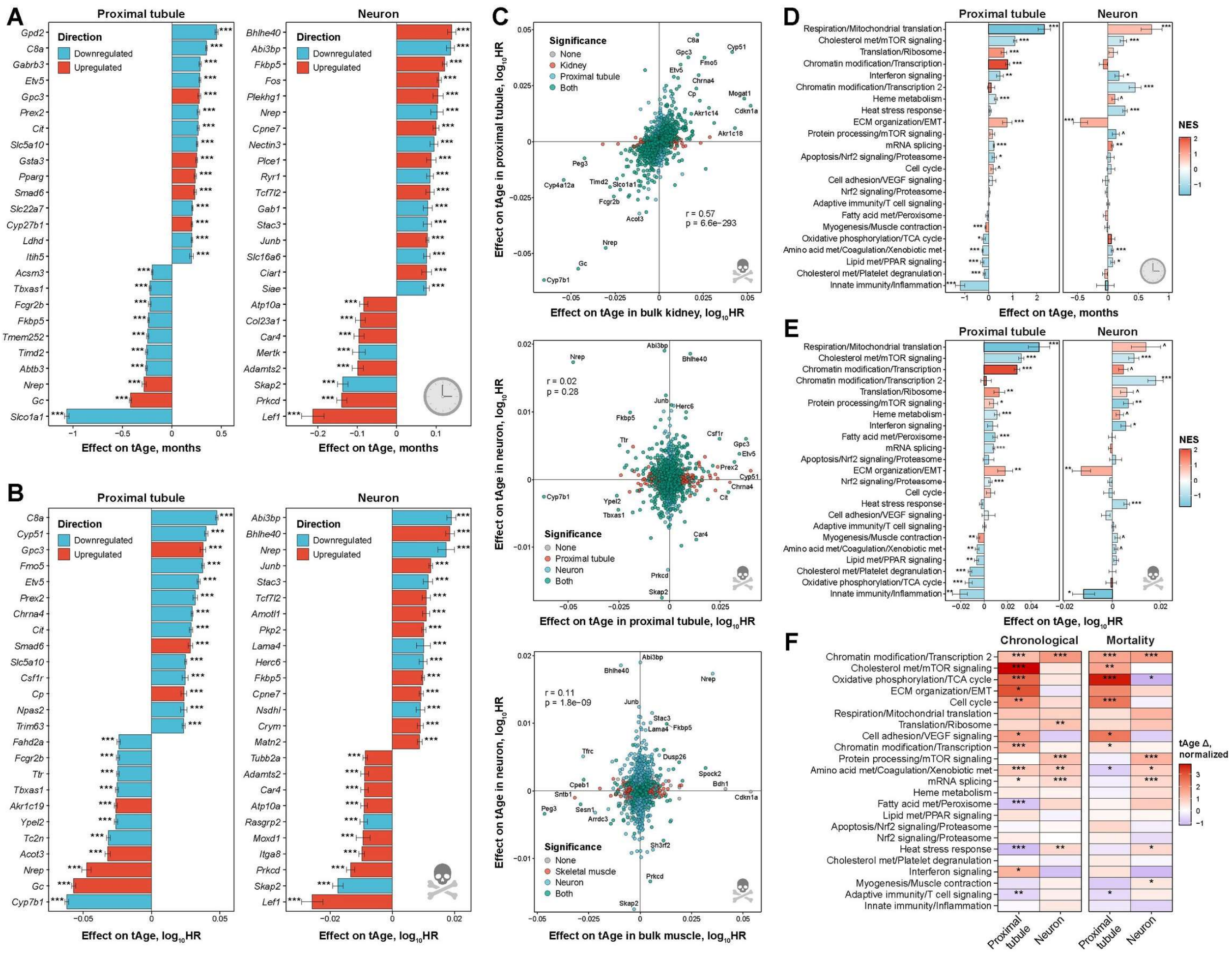


**Extended Data Fig. 15. Molecular mechanisms of pro-aging phenotype in individual cell types of *Klotho* knockout mice.**

A-B. Top genes driving pro/anti-aging (A) and pro-/anti-mortality (B) transcriptomic changes in proximal tubules (left) and neurons (right) of *Klotho* knockout (KO) mice compared to age-matched controls, assessed with rodent multi-tissue EN chronological (A) and mortality (B) clocks. Top 25 genes with the highest absolute effect on tAge difference (logFC * clock coefficient) are shown. Genes up- and downregulated in *Klotho* KO mice are colored in red and blue, respectively. Statistical significance of logFC for each gene is indicated with asterisks. Data are mean tAge difference ± SE.

C. Association between weighted gene expression signatures of mortality (logFC * clock coefficient) of *Klotho* KO mice in proximal tubule and bulk kidney (up), in proximal tubule and neurons (middle), and in bulk skeletal muscle and neurons (bottom), according to the rodent multi-tissue EN mortality clock. The union of top 2,000 genes differentially expressed in *Klotho* KO (with the lowest p-values) are shown on the plot for each pair of signatures. Statistical significance (BH-adjusted p-value < 0.05) is indicated with color. Pearson correlation coefficient and corresponding p-value are shown in text.

D-E. Contributions of gene expression modules into chronological (D) and mortality (E) tAge difference between control and *Klotho* KO mice in proximal tubule (left) and neurons (right), estimated with the rodent multi-tissue EN clocks. Individual modules are shown in rows and names after representative functions. Positive and negative values reflect pro- and anti-aging changes in *Klotho* KO mice compared to age-matched controls, respectively. Statistical significance for each module was assessed with the two-sample unpaired t-test and denoted with asterisks. Bars are colored based on normalized enrichment scores (NES) from gene set enrichment analysis (GSEA), reflecting if genes associated with a particular module are generally up- (red) or downregulated (blue) in *Klotho* KO mice. Modules significantly enriched for up- or downregulated genes (BH-adjusted p-value < 0.05) are visualized with thick bars. Data are means ± SE.

F. Normalized chronological (left) and mortality (right) tAge difference between control and *Klotho* KO mice for proximal tubules and neurons assessed with all module-specific multi-tissue chronological (left) and mortality (right) clocks. Color and asterisks reflect size and statistical significance of tAge difference between control and *Klotho* KO mice, assessed with the two-sample unpaired t-test. Higher and lower tAges in *Klotho* KO mice are shown in red and blue, respectively.

HR: Hazard Ratio; met: metabolism; ECM: Extracellular matrix; EMT: Epithelial-Mesenchymal Transition. ^ p.adj < 0.1; * p.adj < 0.05; ** p.adj < 0.01; *** p.adj < 0.001.

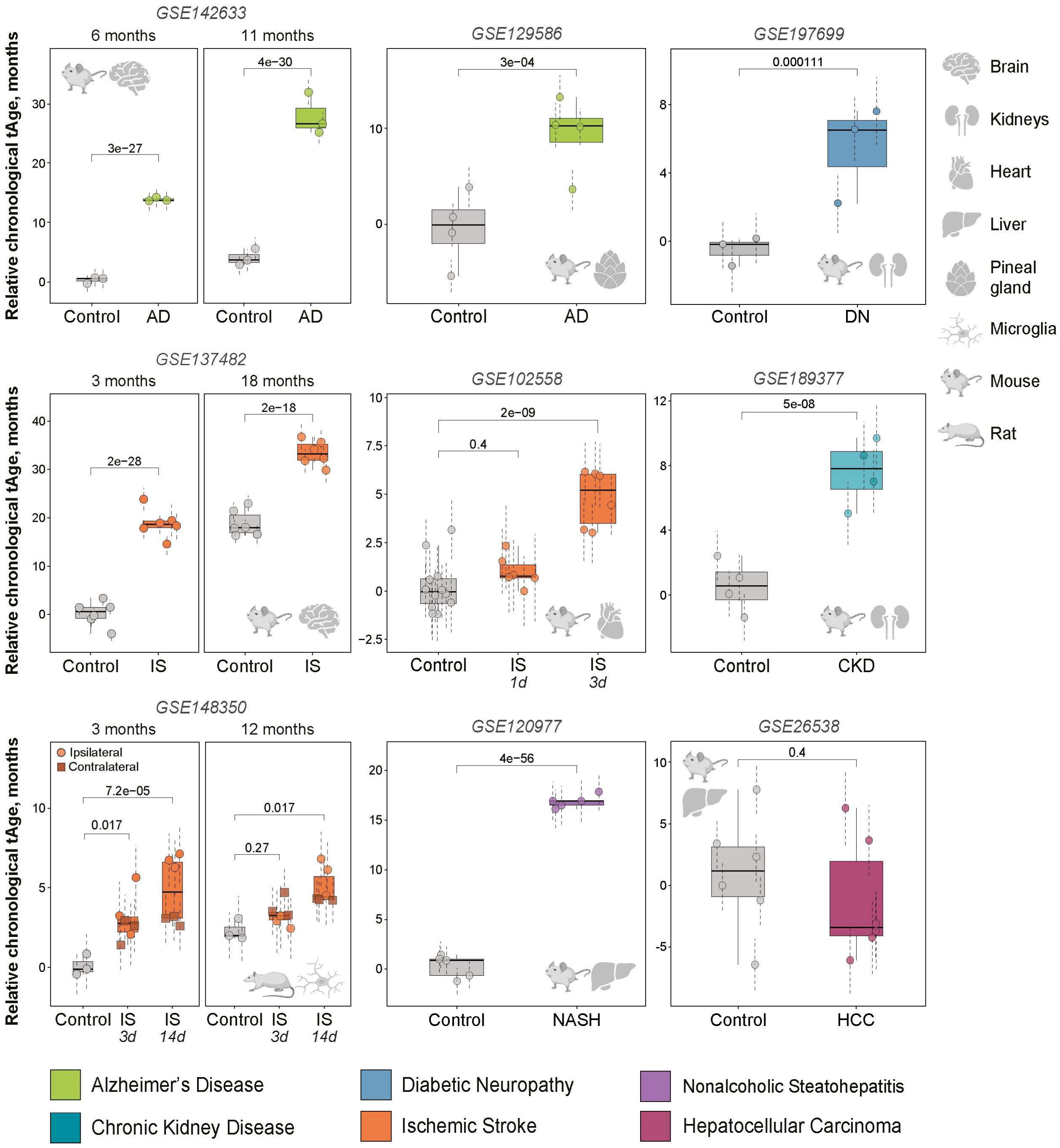


**Extended Data Fig. 16. Chronological transcriptomic age (tAge) of healthy rodents and animals with various models of age-related diseases.** tAges of tissues from control (grey) and age-matched diseased samples (indicated with color) were assessed with the rodent multi-tissue Bayesian Ridge (BR) chronological clock. tAges between the groups were compared with mixed effect model, and corresponding BH-adjusted p-values are shown in text. Species and organs are depicted with icons. GEO IDs of the datasets are provided in text. Data are tAges ± SE.

AD: Alzheimer's disease; DN: Diabetic Neuropathy; IS: Ischemic Stroke; CKD: Chronic Kidney Disease; NASH: Nonalcoholic Steatohepatitis; HCC: Hepatocellular Carcinoma.

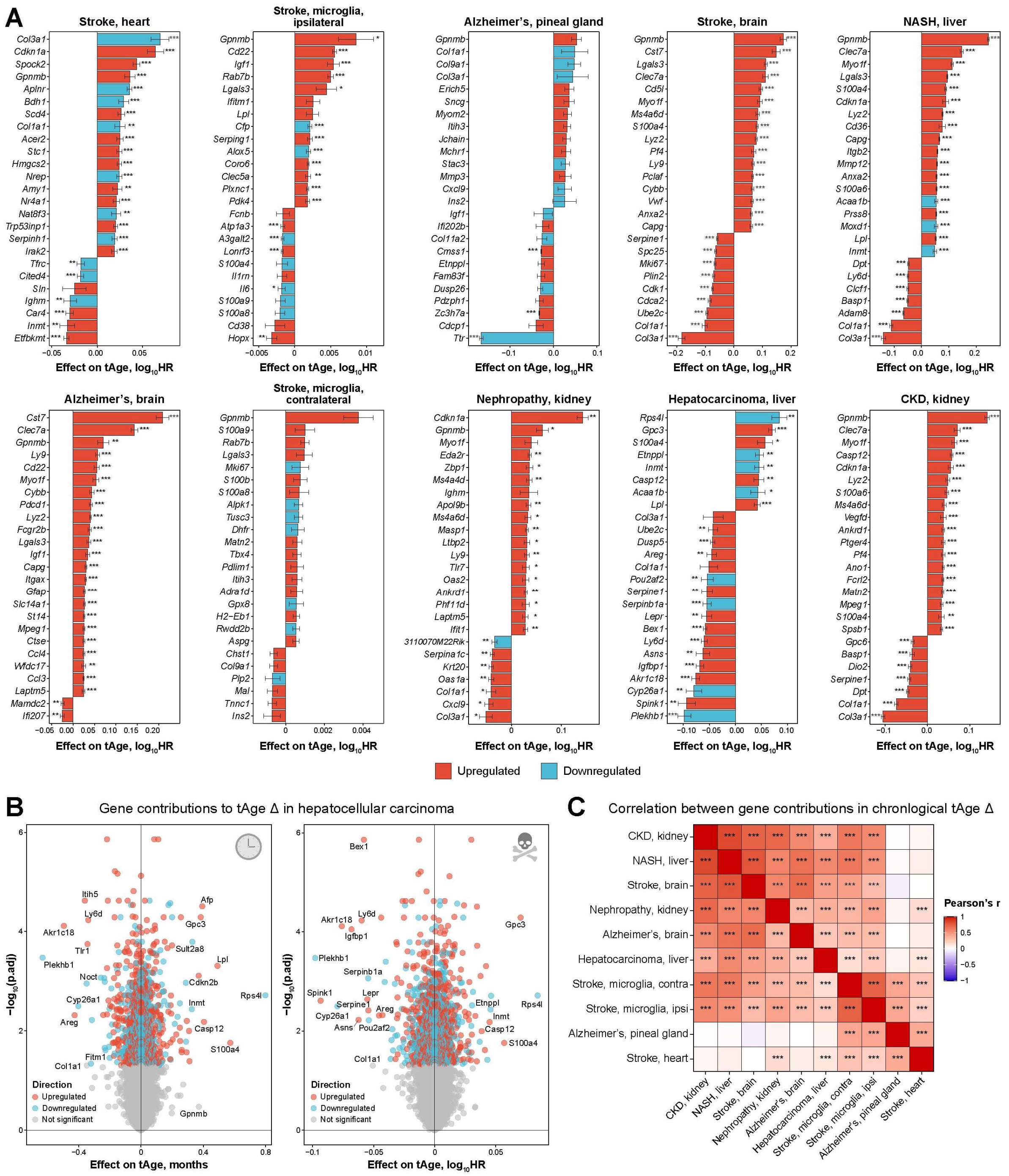


**Extended Data Fig. 17. Genes associated with the pro-aging and pro-mortality molecular phenotype of age-related diseases.**

A. Top genes driving pro- or anti-mortality transcriptomic changes in various models of age-related diseases in mice and rats. Top 25 genes with the highest absolute effect on tAge difference or slope (for microglia stroke models) according to the rodent multi-tissue Elastic Net (EN) mortality clock are shown. Genes up- and downregulated in diseased animals are colored in red and blue, respectively. Statistical significance of logFC for each gene is indicated with asterisks. Data are mean tAge difference ± SE.

B. Volcano plots of gene expression contributions to chronological and mortality tAge difference in hepatocellular carcinoma (HCC) model, according to the rodent multi-tissue EN clocks. The effect of the corresponding gene on tAge (logFC * clock coefficient) and BH-adjusted p-value of its change in HCC compared to age-matched controls are shown on x and y axes, respectively.

Statistically significant differentially expressed genes (BH-adjusted p-value < 0.05) are colored in blue or red if they are downregulated or upregulated in HCC samples, respectively.

C. Pearson correlation between weighted gene expression signatures of aging (logFC * clock coefficient) across various models of age-related diseases, according to the rodent multi-tissue EN chronological clock. The union of top 500 differentially expressed genes (with the lowest p-value) were used to estimate Pearson's correlation coefficient for each pair of signatures. Statistical significance is indicated with asterisks.

HR: Hazard Ratio; CKD: Chronic Kidney Disease; NASH: Nonalcoholic Steatohepatitis; ipsi: ipsilateral; contra: contralateral; ECM: Extracellular matrix; EMT: Epithelial-Mesenchymal Transition; met: metabolism. * p.adj < 0.05; ** p.adj < 0.01; *** p.adj < 0.001.

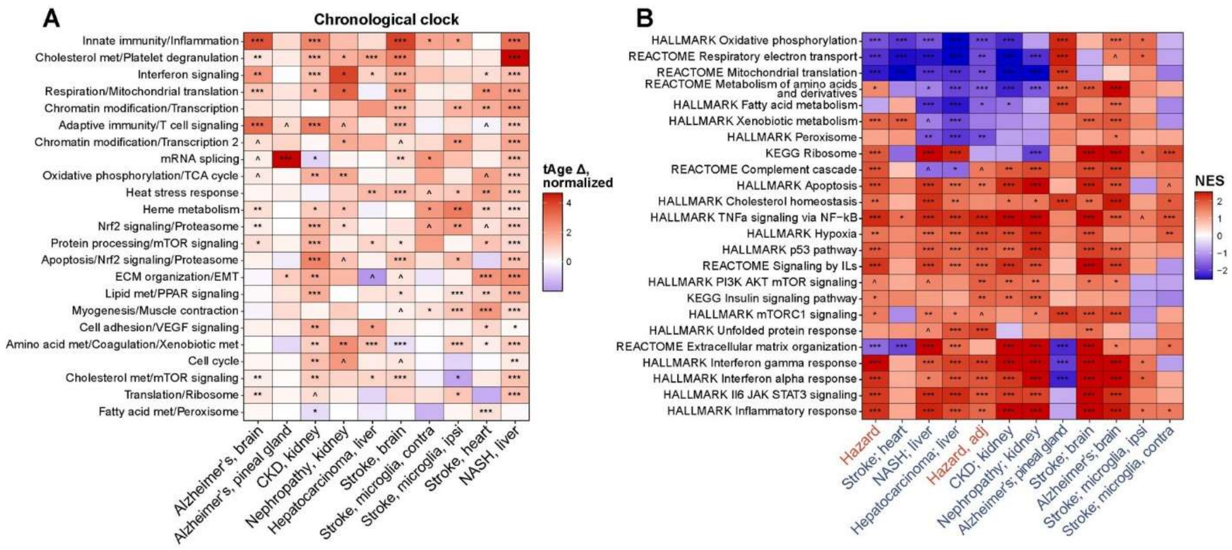


**Extended Data Fig. 18. Functional characterization of pro-aging molecular changes in the models of age-related disease.**

A. Normalized chronological tAge difference between control and age-matched diseased animals assessed with all module-specific multi-tissue chronological clocks. Color and asterisks reflect size and statistical significance (BH-adjusted p-values) of tAge difference between control and disease groups, assessed with the ANOVA or linear regression (for stroke microglia data). Increased and decreased tAges in disease models are shown in red and blue, respectively.

B. Functional enrichment (GSEA) of gene expression changes induced in the models of age-related diseases and signatures of mortality. Only functions significantly enriched by at least one signature are shown (BH-adjusted p-value < 0.05). The whole list of enriched functions is in Supplementary Table 5B. NES: Normalized Enrichment Score.

CKD: Chronic Kidney Disease; NASH: Nonalcoholic Steatohepatitis; ipsi: ipsilateral; contra: contralateral; ECM: Extracellular matrix; EMT: Epithelial-Mesenchymal Transition; met: metabolism. ^ p.adj < 0.1; * p.adj < 0.05; ** p.adj < 0.01; *** p.adj < 0.001.

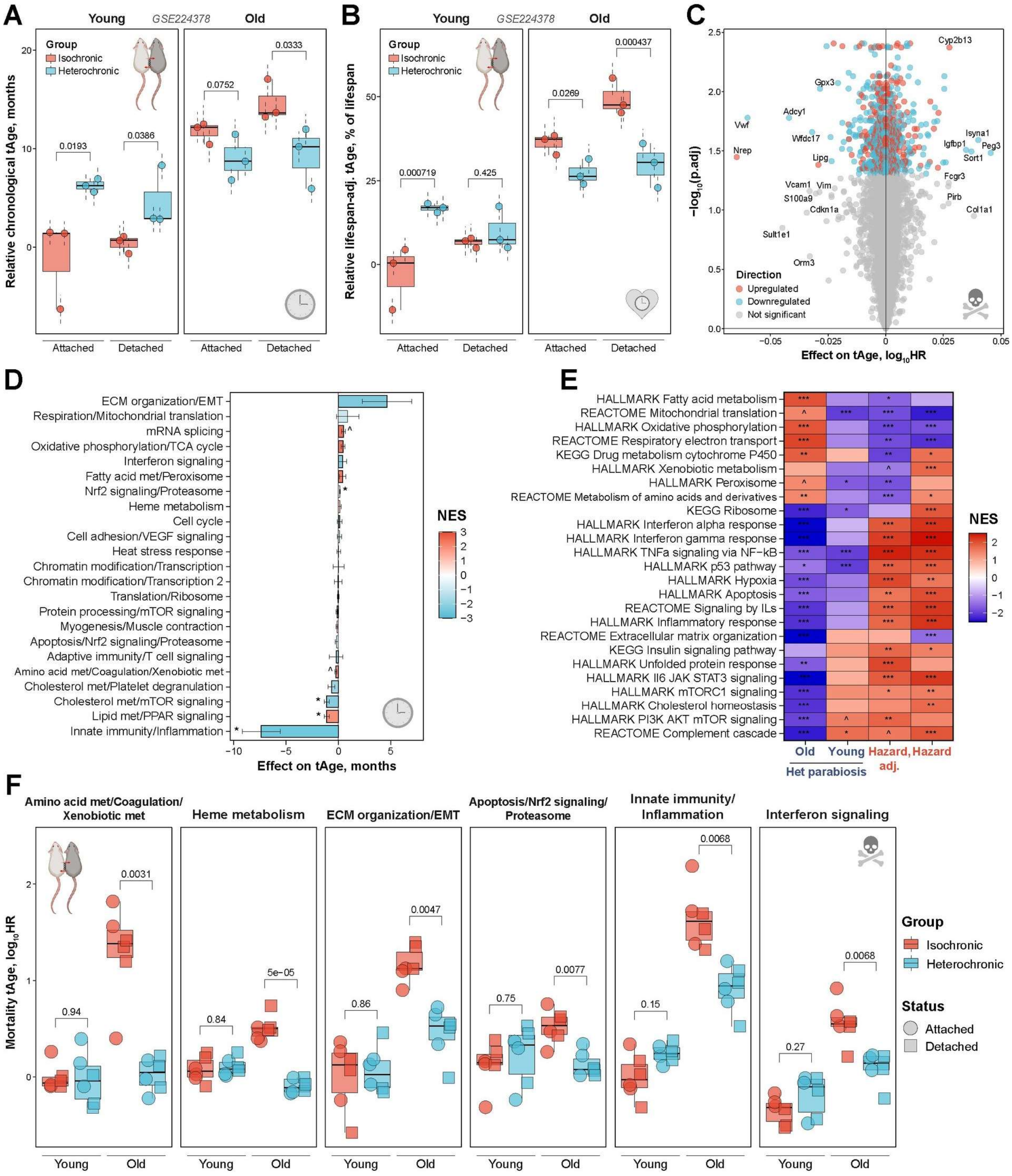


**Extended Data Fig. 19. Genes and functions associated with the molecular rejuvenation effect of heterochronic parabiosis.**

A-B. Chronological (A) and lifespan-adjusted (B) transcriptomic age (tAge) of livers from 3-month-old (left) and 20-month-old (right) mice subjected to isochronic or heterochronic parabiosis for 3 months, as assessed with the rodent multi-tissue Bayesian Ridge (BR) chronological and lifespan-adjusted clocks. tAges between the groups were compared with a mixed effect model, and corresponding BH-adjusted p-values are shown in text. Data are tAges ± SE.

C. Volcano plot of gene expression contributions to mortality tAge difference in old mice subjected to heterochronic parabiosis, according to the rodent multi-tissue EN mortality clock. The effect of the corresponding gene on tAge (logFC * clock coefficient) and BH-adjusted p-value of its change in heterochronic parabiosis group compared to age-matched isochronic controls are shown on x and y axes, respectively. Statistically significant differentially expressed genes (BH-adjusted p-value < 0.05) are colored in blue or red if they are downregulated or upregulated in heterochronic group, respectively.

D. Contributions of transcriptomic modules to chronological tAge difference in old mice subjected to heterochronic parabiosis, estimated with the rodent multi-tissue Elastic Net (EN) chronological clock. Individual modules are shown in rows and named after representative functions. Statistical significance for each module was assessed with ANOVA and indicated with asterisks. Bars are colored based on normalized enrichment scores (NES) from gene set enrichment analysis (GSEA), reflecting if genes associated with a particular module are generally up- (red) or downregulated (blue) in old mice subjected to heterochronic parabiosis compared to the age-matched isochronic parabiosis group. Modules significantly enriched for up- or downregulated genes (BH-adjusted p-value < 0.05) are visualized with thick bars. Data are means ± SE.

E. Functional enrichment (GSEA) of gene expression changes induced in young and old mice subjected to heterochronic parabiosis compared to the age-matched isochronic parabiosis group, and signatures of mortality. Only functions significantly enriched by at least one signature are shown (BH-adjusted p-value < 0.05). The whole list of enriched functions is in Supplementary Table 5C. Het: Heterochronic.

F. Mortality tAge in young and old mice subjected to isochronic and heterochronic parabiosis for 3 months estimated with representative module-specific multi-tissue clocks. The shape of dots reflects an attachment status. Difference between the groups was assessed with the ANOVA, and corresponding BH-adjusted p-values are shown in text.

HR: Hazard Ratio; ECM: Extracellular matrix; EMT: Epithelial-Mesenchymal Transition; met: metabolism. ^ p.adj < 0.1; * p.adj < 0.05; ** p.adj < 0.01; *** p.adj < 0.001.

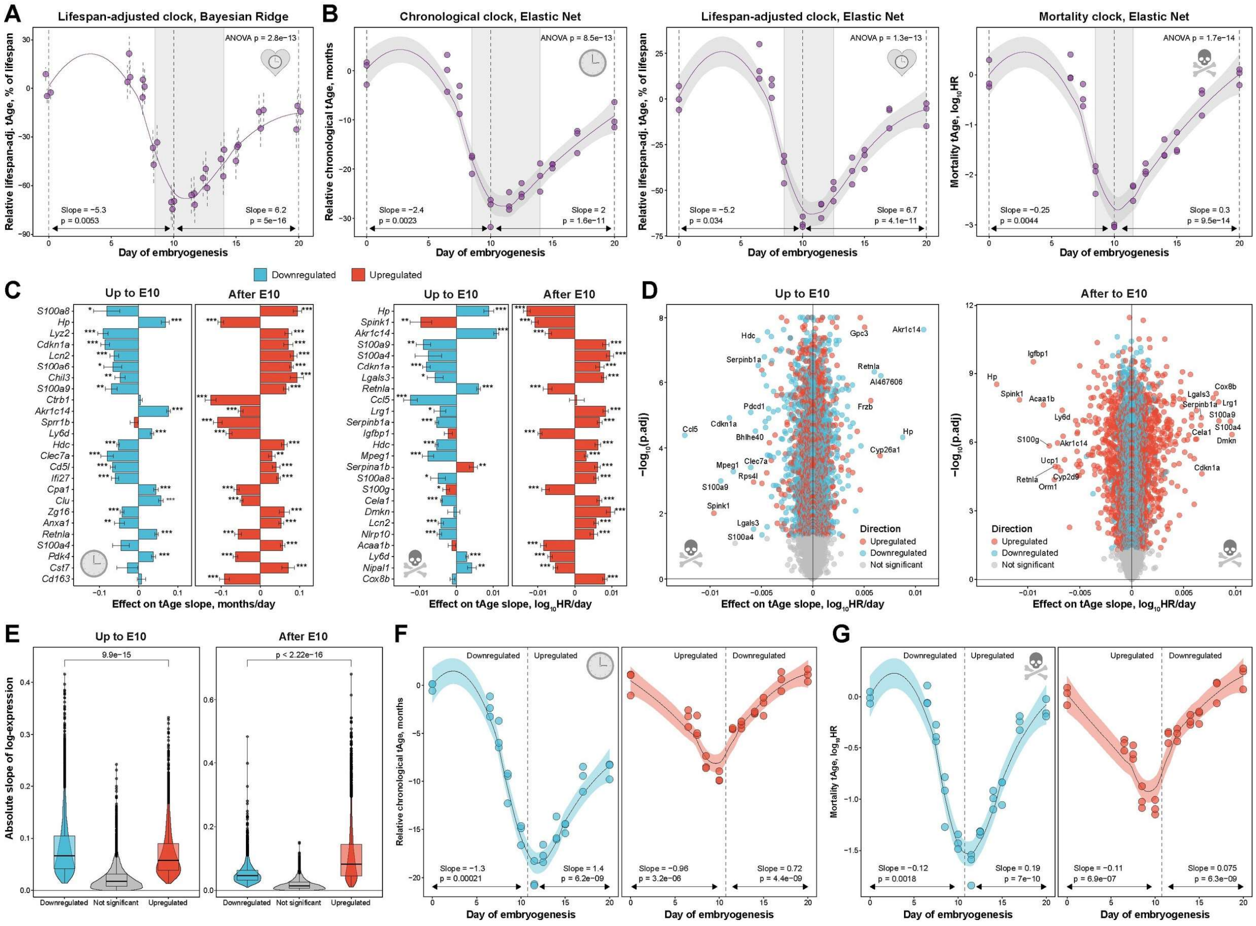


**Extended Data Fig. 20. Dynamics of transcriptomic age during mouse embryonic development.**

A. Lifespan-adjusted transcriptomic age (tAge) of mouse embryos during development, assessed with the mouse multi-tissue Bayesian Ridge (BR) lifespan-adjusted clock. Overall change of tAge was assessed with the mixed effect ANOVA, whereas slopes of tAge change up to day 10 and after day 10 were assessed with a mixed effect linear model. Corresponding p-values and slope estimates are shown in text. Loess regression curve is shown with a purple line. Dotted line reflects time point with the minimum average tAge, and 95% confidence interval that contains putative minimum of embryo's biological age (ground zero state) is shown with shaded grey rectangular. Data are tAges ± SE.

B. Chronological (left), lifespan-adjusted (middle) and mortality (right) tAge of mouse embryos during development, assessed with the mouse multi-tissue Elastic Net (EN) clocks. Overall change of tAge was assessed with ANOVA, whereas slopes of tAge change up to day 10 and after day 10 were assessed with a linear regression model. Corresponding p-values and slope estimates are shown in text. Purple line and shaded grey area around it reflect loess regression curve and its 95% confidence interval, respectively. Dotted line shows time point with the minimum average tAge, and 95% confidence interval that contains putative minimum of embryo's biological age (ground zero state) is shown with shaded grey rectangular.

C. Top genes driving pro- or anti-aging (left) and mortality (right) transcriptomic changes in mouse embryos up to day 10 and after day 10 of development, according to the rodent multi-tissue EN chronological (left) and mortality (right) clocks. Top 25 genes with the highest average absolute effect on tAge slope (slope * clock coefficient) are shown. Genes up- and downregulated during given stage of embryogenesis are colored in red and blue, respectively. Statistical significance of logFC for each gene is indicated with asterisks. Data are tAge slope estimates ± SE.

D. Volcano plot of gene expression contributions to mortality tAge change in mouse embryos up to day 10 and after day 10 of developemnt, according to the rodent multi-tissue EN mortality clock. The effect of the corresponding gene on tAge (slope * clock coefficient) and BH-adjusted p-value of its change during the given period are shown on x and y axes, respectively. Statistically significant differentially expressed genes (BH-adjusted p-value < 0.05) are colored in blue or red if they are downregulated or upregulated during the corresponding stage of embryogenesis, respectively.

E. Absolute slope of gene expression changes in mouse embryos up to day 10 (left) and after day 10 (right) of development for genes that are significantly (BH-adjusted p-value < 0.05) upregulated (red) and downregulated (blue) during the given stage of embryogenesis as well as for genes without statistically significant change of expression (grey). Median scale of expression changes between up- and downregulated genes was compared with Wilcoxon rank sum test, corresponding p-values are shown in text.

F-G. Trajectories of chronological (F) and mortality (G) tAges during mouse embryogenesis calculated only for genes downregulated or upregulated up to day 10 and after day 10 of development, assessed with the mouse multi-tissue EN clocks. tAges based on genes downregulated up to day 10 or upregulated after day 10 are shown in blue, while tAges based on genes with the opposite behavior are shown in red. Black lines and shaded areas around them reflect loess regression curves and their 95% confidence intervals, respectively. Slopes of tAge change up to day 10 and after day 10 were assessed with a linear regression model. Corresponding p-values and slope estimates are shown in text.

HR: Hazard Ratio. * p.adj < 0.05; ** p.adj < 0.01; *** p.adj < 0.001.

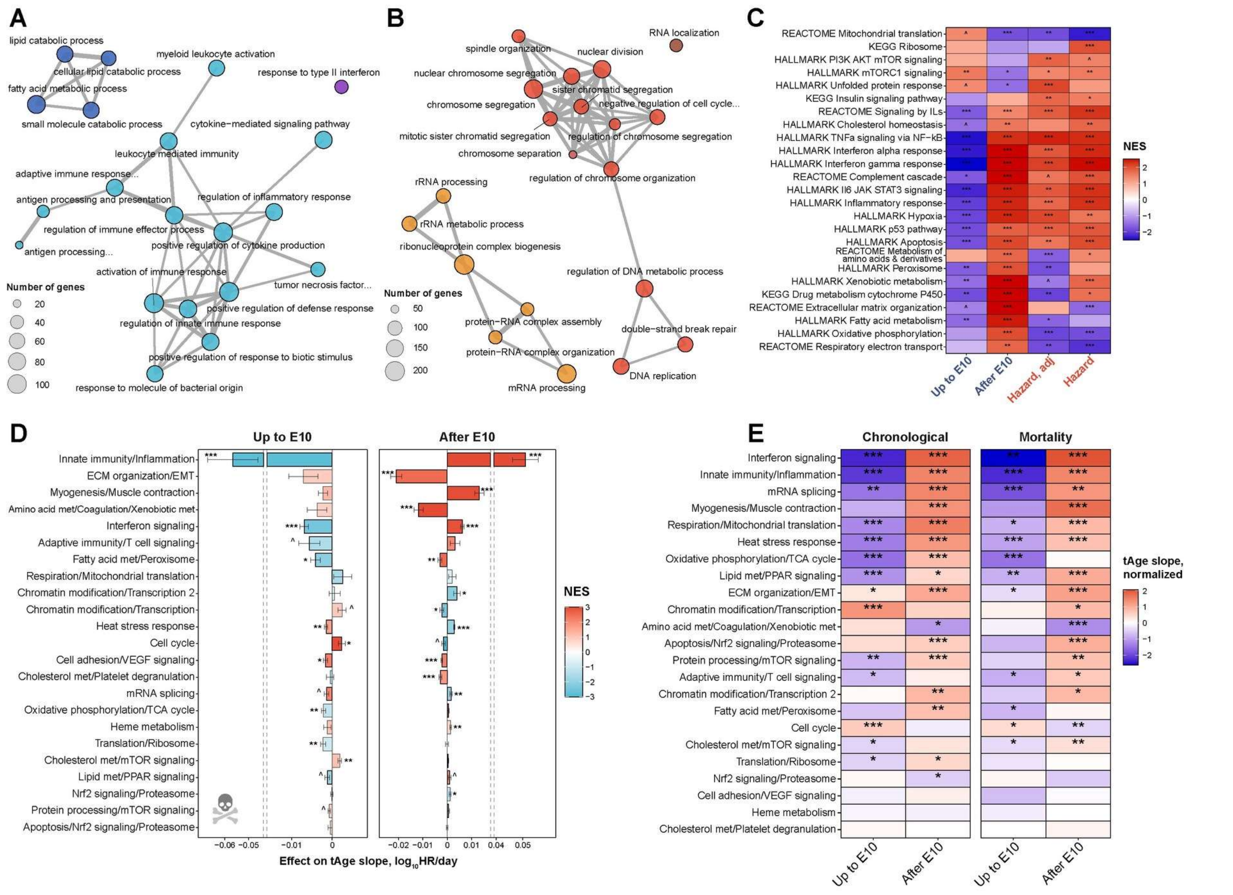


**Extended Data Fig. 21. Functional characterization of aging- and mortality-associated gene expression changes during embryogenesis.**

A-B. Network of functional terms enriched for genes downregulated up to day 10 of mouse embryogenesis and upregulated afterwards (A) or vice versa (B). Statistical significance of enrichment was assessed with Fisher's exact test. Each dot reflects a functional term from GO BP ontology that is significantly enriched (BH-adjusted p-value < 0.05) for the corresponding gene list. Number of genes affected during embryogenesis and associated with each term is indicated with dot size. Semantically similar terms are connected by edges. Colors represent distinct clusters of enriched terms. The whole list of enriched functions is in Supplementary Table 6.

C. Functional enrichment (GSEA) of gene expression changes during mouse embryogenesis up to day 10 and after day 10 of development, and signatures of mortality. Only functions significantly enriched by at least one signature are shown (BH-adjusted p-value < 0.05). The whole list of enriched functions is in Supplementary Table 5D. NES: Normalized Enrichment Score.

D. Contributions of transcriptomic modules into slope of mortality tAge change in mouse embryos up to day 10 (left) and after day 10 (right) of development estimated with the multi-tissue Elastic Net (EN) mortality clock. Statistical significance for each module was assessed with the linear regression and indicated with asterisks. Bars are colored based on NES from GSEA, reflecting if genes associated with a particular module are generally up- (red) or downregulated (blue) during the corresponding stage of embryogenesis. Modules significantly enriched for up- or downregulated genes (BH-adjusted p-value < 0.05) are visualized with thick bars. Data are slopes ± SE. HR: Hazard Ratio.

E. Normalized slope of chronological (left) and mortality (right) tAge change in mouse embryos up to day 10 or after day 10 of development assessed with all module-specific multi-tissue chronological (left) and mortality (right) clocks. Color and asterisks reflect size and statistical significance of tAge change during the given stage of embryogenesis, assessed with the linear regression model. Positive and negative slopes of tAge are shown in red and blue, respectively.

ECM: Extracellular matrix; EMT: Epithelial-Mesenchymal Transition; met: metabolism. ^ p.adj < 0.1; * p.adj < 0.05; ** p.adj < 0.01; *** p.adj < 0.001.

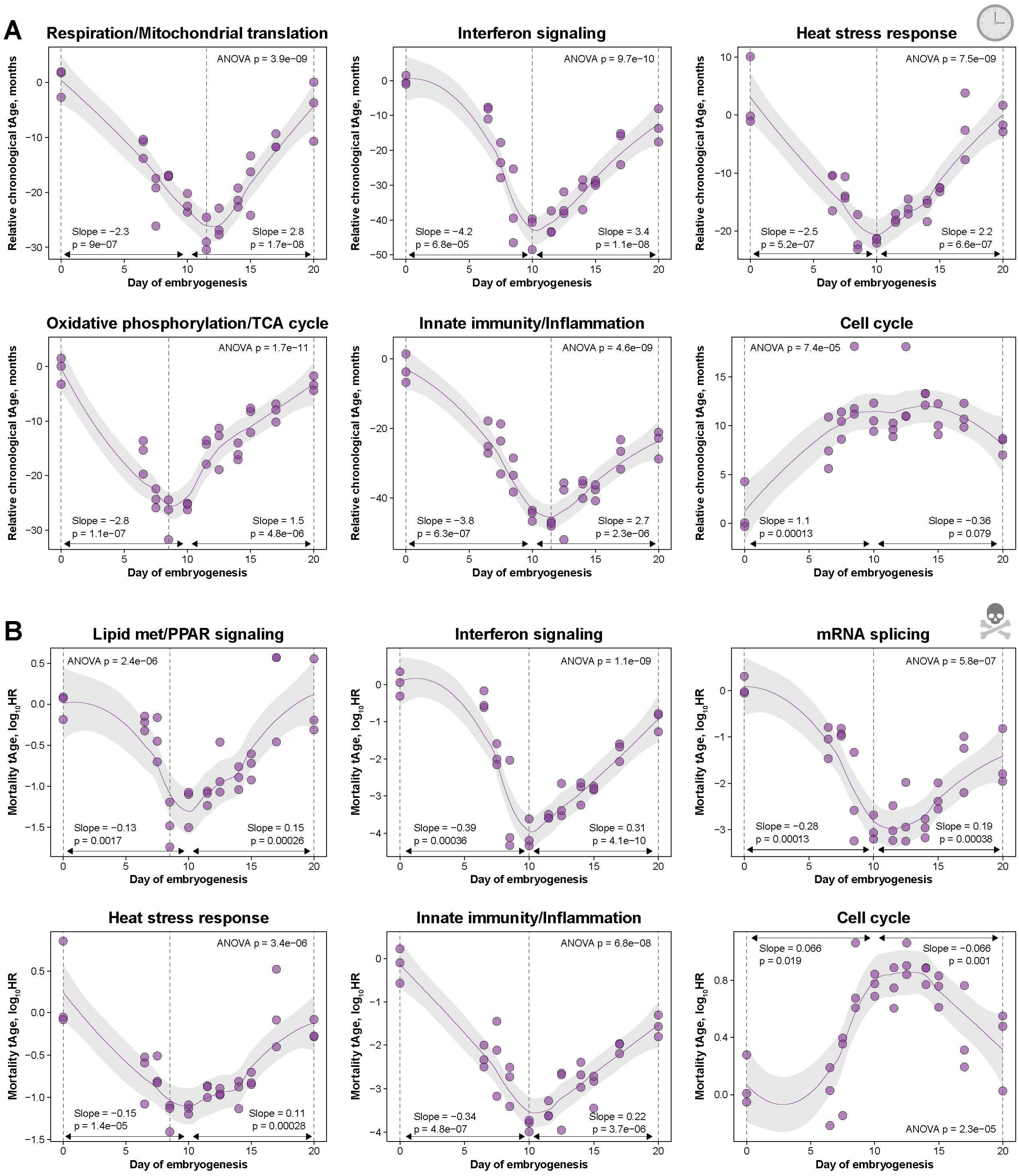


**Extended Data Fig. 22. Aging- and mortality-associated molecular changes during embryogenesis characterized by representative module-specific transcriptomic clocks.**

A-B. Chronological (A) and mortality (B) tAge of mouse embryos during development estimated with representative module-specific multi-tissue clocks. Slope of change up to day 10 and after day 10 was assessed with the linear regression. Corresponding p-values and slope estimates are shown in text. Purple line and shaded grey area around it reflect loess regression curve and its 95% confidence interval, respectively. Results for all module-specific chronological and mortality clocks are in Extended Data Fig. 21E.

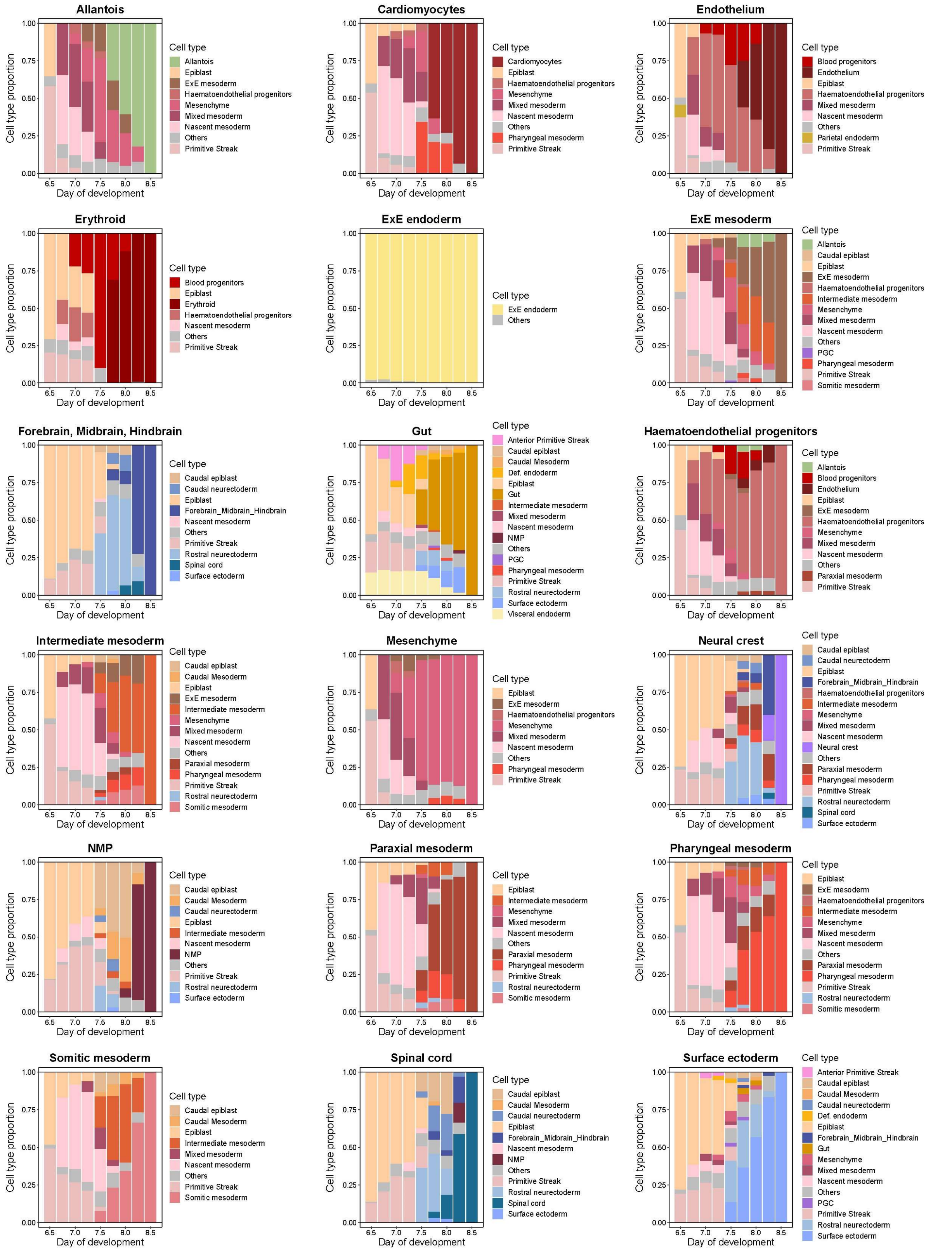


**Extended Data Fig. 23. Proportions of cell types contributing to lineages of cell types presented at day 8.5 of mouse embryonic development.** For every cell lineage and day of development, bars represent sums of probabilities of being an ancestor for each annotated cell type presented at this stage of development. Top cell types that cover more than 90% of the total probability at the given day are shown, while remaining cell types are combined in the "Others" category. ExE: Extraembryonic; PGC: Primordial germ cells; NMP: Neuromesodermal Progenitors.

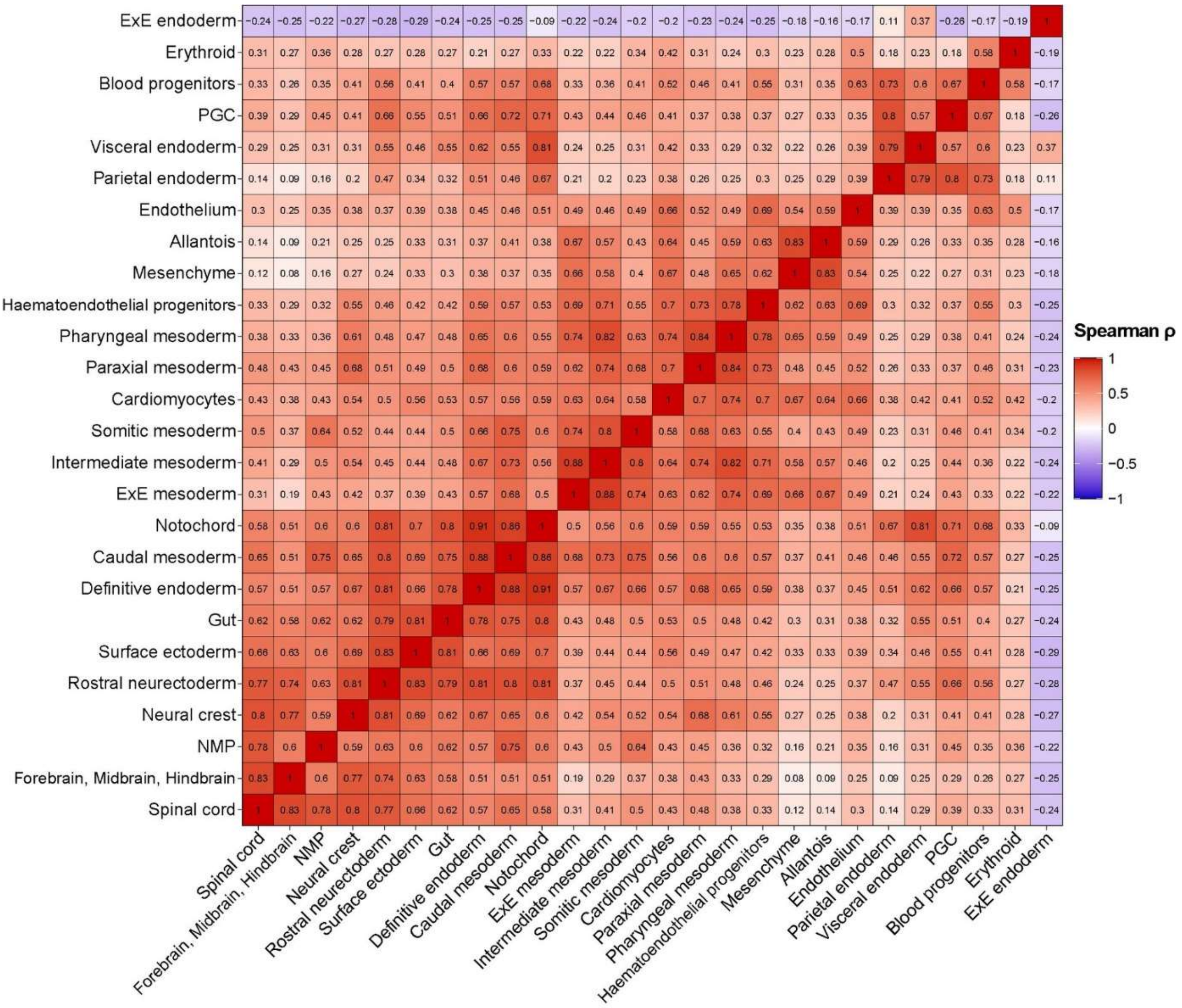


**Extended Data Fig. 24. Spearman correlation of cell lineage trajectories between days 6.5 and 8.5 of mouse embryonic development.** For every pair of cell types presented at day 8.5, pairwise correlation coefficient between vectors of probabilities that cells are included in the corresponding lineage was calculated. Clustering was performed with complete hierarchical method based on correlation distances. ExE: Extraembryonic; NMP: Neuromesodermal Progenitors.

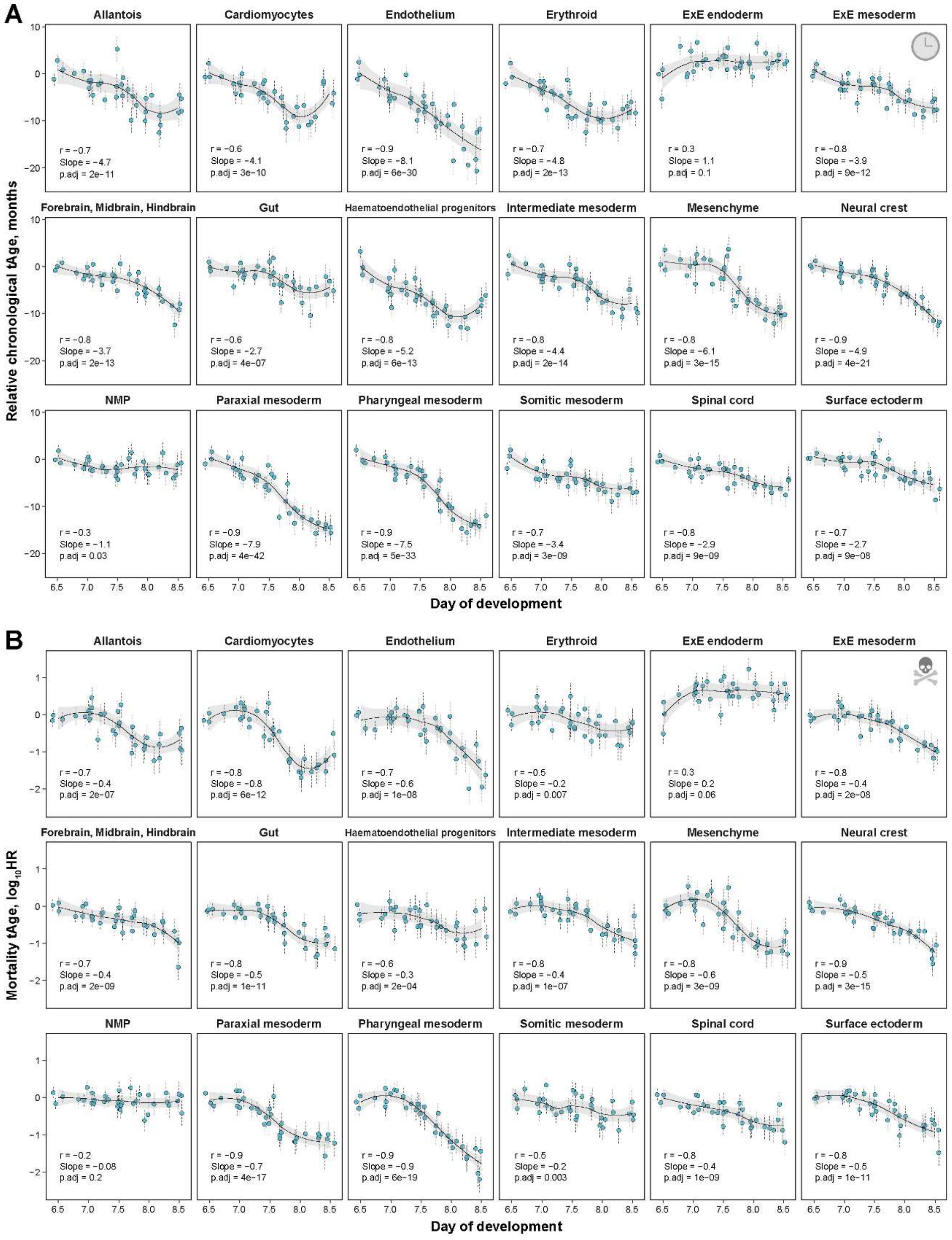

**Extended Data Fig. 25. Transcriptomic age trajectory of individual cell lineages between days 6.5 and 8.5 of mouse embryonic development.**

A-B. Chronological (A) and mortality (B) transcriptomic ages (tAges) of metacells representing lineages of cell types presented at day 8.5 of embryogenesis, assessed with the mouse multi-tissue Bayesian Ridge (BR) chronological (A) and mortality (B) clocks. Slope of tAge change was assessed with a mixed effect linear model. Pearson correlation coefficient, slope estimate and corresponding p-value are shown with text. Black line and shaded grey area around it reflect loess regression curve and its 95% confidence interval, respectively. Data are tAges ± SE. HR: Hazard Ratio; ExE: Extraembryonic; NMP: Neuromesodermal Progenitors.

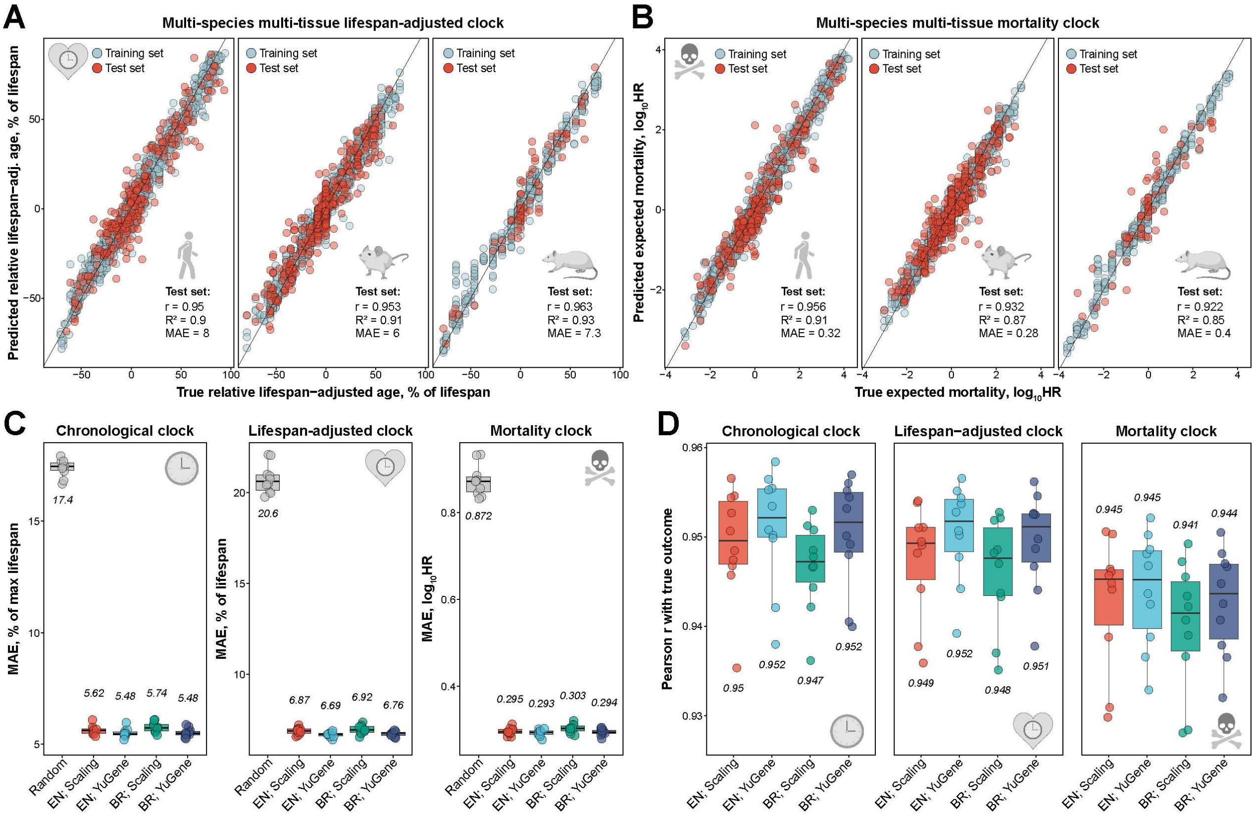


**Extended Data Fig. 26. Quality of multi-species multi-tissue transcriptomic clocks of chronological age, lifespan-adjusted age and mortality.**

A-B. Accuracy of prediction of relative lifespan-adjusted age (A) and expected mortality (B) with multi-tissue transcriptomic clocks trained with Elastic Net (EN) model on human, mouse and rat data (with scaling normalization). Accuracy of the trained model was assessed separately for humans (left), mice (middle) and rats (right). Training and test sets are denoted by color. Pearson correlation coefficient, $R^2$ and mean absolute error (MAE) for test sets are shown in text. HR: Hazard Ratio.

C. Mean absolute error (MAE) of predictions of relative chronological age (left), lifespan-adjusted age (middle) and expected mortality (righ) on 10 randomly chosen test sets with multi-species multi-tissue transcriptomic clocks based on EN or Bayesian Ridge (BR) model and scaling or YuGene normalization. Median estimate of quality across 10 runs is provided in text. MAE of random prediction is shown in grey. HR: Hazard Ratio.

D. Pearson correlation between real outcomes and predictions of relative chronological age (left), lifespan-adjusted age (middle) or expected mortality (righ) on 10 randomly chosen test sets with multi-species multi-tissue transcriptomic clocks based on EN or BR model and scaling or YuGene normalization. Median estimate of quality across 10 runs is provided in text.

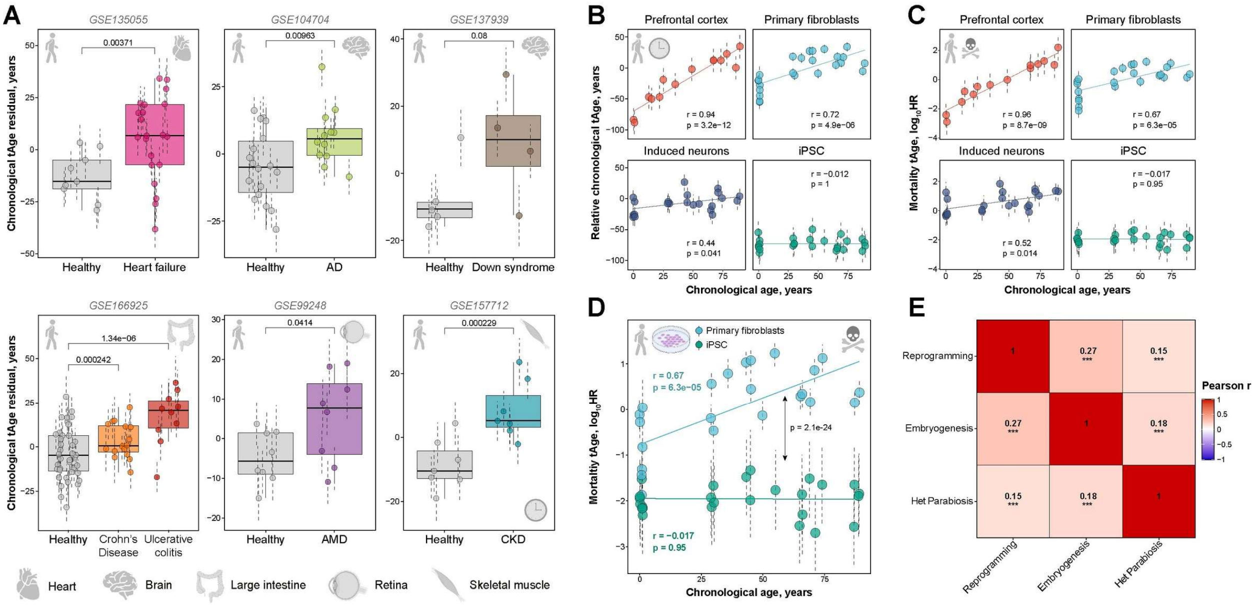


**Extended Data Fig. 27. Association of transcriptomic age with human diseases and cellular reprogramming.**

A. Chronological transcriptomic age (tAge) residual of tissues from healthy people (grey) and patients diagnosed with age-related diseases (indicated with color) adjusted for chronological age and sex, assessed with the multi-species multi-tissue Bayesian Ridge (BR) chronological clock. tAge between the groups were compared with mixed effect model, and corresponding BH-adjusted p-values are shown in text. Organs are depicted with icons. GEO IDs of corresponding datasets are provided in text. Data are tAges ± SE.

B-C. Association between chronological age (on x axis) and chronological (B) or mortality (C) transcriptomic age (on y axis) for various human cell types. tAges were calculated with the multi-species multi-tissue BR chronological (B) and mortality (C) clocks. Association between chronological age and tAge for each cell type was assessed with a mixed effect model. Corresponding p-value and Pearson correlation coefficient are shown in text. Data are tAges ± SE. HR: Hazard Ratio. AD: Alzheimer's Disease; AMD: Age-Related Macular Degeneration; CKD: Chronic Kidney Disease.

D. Mortality tAge of primary fibroblasts and reprogrammed iPSCs from patients of different chronological ages, assessed with the multi-species multi-tissue BR mortality clock. Association between chronological age and tAge for each cell type was assessed with a mixed effect model. Corresponding p-value and Pearson correlation coefficient are shown with colored text. tAges of primary fibroblasts and corresponding iPSCs were compared with a mixed effect model, where patient ID was included as a factor covariate. The corresponding p-value is provided in black. Data are tAges ± SE. HR: Hazard Ratio.

E. Pearson correlation between weighted gene expression signatures of mortality of early mouse embryogenesis, old mouse heterochronic parabiosis and human iPSCs models. The union of top 2,000 differentially expressed genes (with the lowest p-value) for each pair of signatures was used to calculate the correlation coefficient. Correlation coefficient and statistical significance are indicated with text and asterisks, respectively. * p.adj < 0.05; ** p.adj < 0.01; *** p.adj < 0.001.

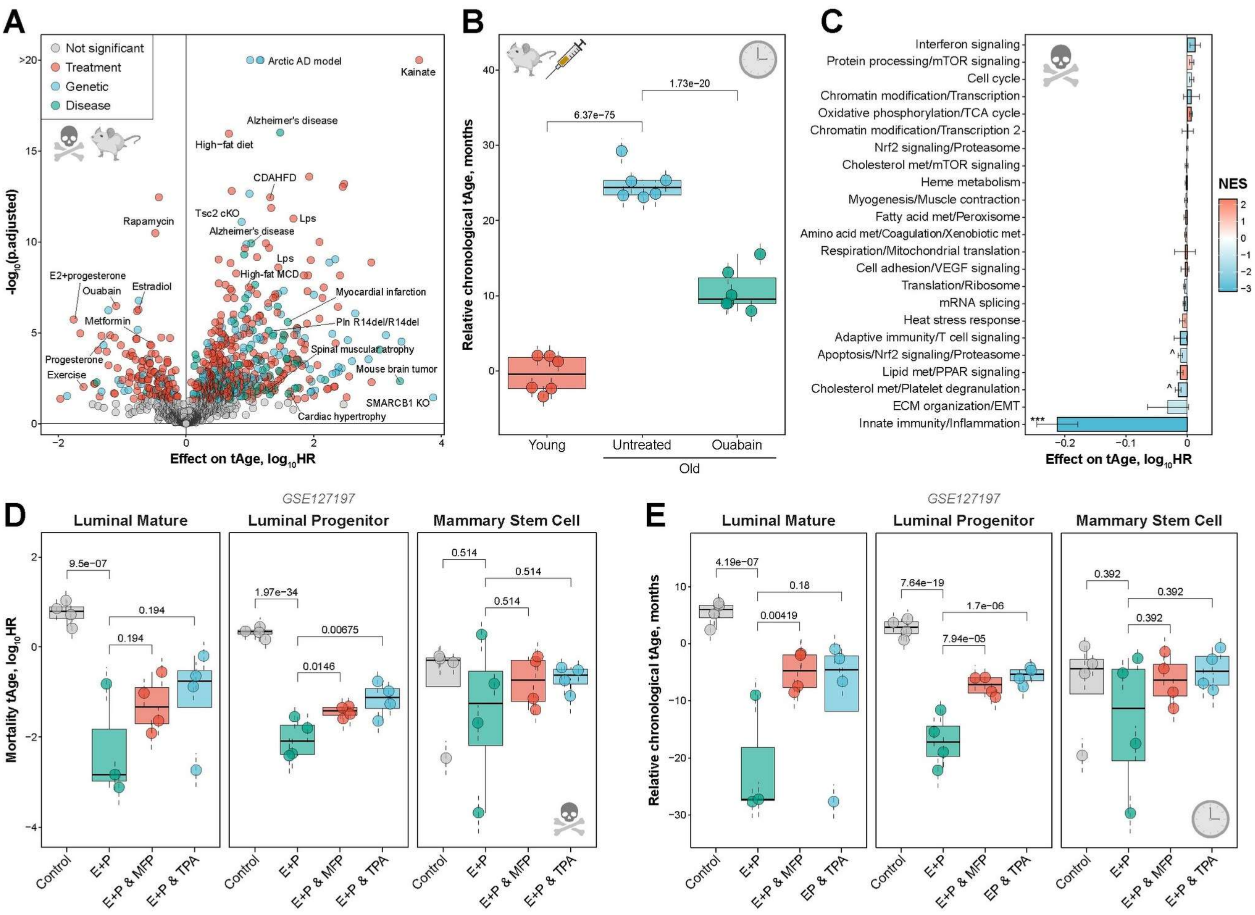


**Extended Data Fig. 28. Transcriptomic clocks allow to identify and characterize novel interventions that affect mammalian aging and mortality.**

A. Mouse interventions identified with the rodent multi-tissue Elastic Net (EN) mortality clock in the GEO screening performed on the dataset collection from Clockbase. Every dot represents an intervention from the particular dataset. tAge difference between control and intervention samples and the corresponding BH-adjusted p-value are plotted on x and y axes, respectively. Significant perturbations (BH-adjusted p-value < 0.05) are indicated with color.

B. Chronological transcriptomic age (tAge) of livers from 7-week-old (Young) and 24-month-old (Old) mice treated with saline (Untreated) or ouabain (Ouabain), examined with the rodent multi-tissue Bayesian Ridge (BR) chronological clock. tAges between the groups were compared with a mixed effect model, and corresponding BH-adjusted p-values are shown in text. Data are tAges ± SE.

C. Contributions of transcriptomic modules to mortality tAge difference between old untreated mice and old mice treated with ouabain, estimated with the rodent multi-tissue EN mortality clock. Individual modules are shown in rows and named after representative functions. Statistical significance for each module was assessed with ANOVA and indicated with asterisks. Bars are colored based on normalized enrichment scores (NES) from gene set enrichment analysis (GSEA), reflecting if genes associated with a particular module are generally up- (red) or downregulated (blue) in mice subjected to ouabain. Modules significantly enriched for up- or downregulated genes (BH-adjusted p-value < 0.05) are visualized with thick bars. Data are means ± SE. HR: Hazard Ratio.

D-E. Mortality (D) and chronological (E) transcriptomic ages (tAges) of various cell types from mammary glands of 3-month-old ovariectomized mice untreated (grey) or treated with estrogen and progesterone either alone (E+P, green) or in combination with progesterone receptor antagonists mifepristone (MFP, red) and telapristone (TPA, blue). tAges were estimated with the rodent multi-tissue BR mortality (D) and chronological (E) clocks. tAges between the groups were compared separately for each cell type with a mixed effect model, and corresponding BH-adjusted p-values are shown in text. Data are tAges ± SE. HR: Hazard Ratio.

Lps: Lipopolysaccharides; AD: Alzheimer's Disease; CDAHFD: choline-deficient, L-amino acid-defined, high-fat diet; MCD: Methionine- and choline-deficient diet; KO: Knockout; E2: Estradiol.
